# Convergent Control of NREM Sleep and Anesthesia by Prefrontal Layer 5 Extratelencephalic Neurons

**DOI:** 10.1101/2025.10.10.681639

**Authors:** Andrzej Z. Wasilczuk, Daiki Takekawa, Joseph Cichon, Xiaoyan Zhang, Ethan B Blackwood, Jeffrey Parra-Munevar, Mean-Hwan Kim, William Spain, Aaron J. Krom, Hirotaka Kinoshita, Brockton J. Keen, E. Railey White, Max B. Kelz, Nikolai Dembrow, Alex Proekt

**Affiliations:** Neuroscience of Unconsciousness and Reanimation Research Alliance, Perelman School of Medicine, University of Pennsylvania, Philadelphia, PA, USA; Department of Anesthesiology and Critical Care, Perelman School of Medicine, University of Pennsylvania, Philadelphia, PA, USA; Department of Anesthesiology, Graduate School of Medicine, Hirosaki University, Hirosaki, Aomori, Japan; Mahoney Institute of Neurosciences, Perelman School of Medicine, University of Pennsylvania, Philadelphia, PA, USA; Neuroscience Graduate Group, Perelman School of Medicine, University of Pennsylvania, Philadelphia, PA, USA; Department of Neurobiology and Biophysics, University of Washington, Seattle, WA 98195, USA; Synapse and Circuits Dynamics Laboratory, Department of Brain Sciences, DGIST, Daegu, South Korea; Epilepsy Center of Excellence, Department of Veterans Affairs Medical Center, Seattle, WA 98108, USA; Department of Anesthesiology, Intensive Care, and Pain Management, Tel Aviv Sourasky Medical Center, Tel Aviv, Israel; Sagol Brain Institute, Tel Aviv Sourasky Medical Center, Tel Aviv, Israel; Chronobiology and Sleep Institute, Perelman School of Medicine, University of Pennsylvania, Philadelphia, PA, USA

## Abstract

The cortical mechanisms that actively suppress consciousness remain poorly understood. Here, we identify a specific subset of prefrontal cortical (PFC) neurons that are activated under anesthesia while most cortical neurons are suppressed. Chemogenetic activation of these excitatory neurons enhances anesthetic potency and deepens NREM sleep, whereas their inhibition blunts anesthetic effects and disrupts sleep. We designate these cells NREM- and Anesthesia-Promoting (NAP) neurons. NAP neurons correspond to layer 5 extratelencephalic (L5 ET) neurons enriched in hyperpolarization-activated cyclic nucleotide-gated (HCN1) expression and characteristic HCN-mediated electrophysiological properties. Mechanistically distinct anesthetics act directly and specifically on PFC L5-ET neurons to increase their excitability. NAPs have sparse cortical projections and predominately innervate subcortical targets including anterior and reticular thalamic nuclei. Using a transgenic mouse line (*Colgalt2-cre*), we confirm that PFC L5 ET neurons are preferentially activated under anesthesia and during deep recovery NREM sleep. Finally, we show that activation of thalamic-projecting Colgalt2-expressing PFC neurons is sufficient to promote deep NREM sleep. These findings identify a novel prefrontal corticothalamic circuit that provides a shared cortical substrate for naturally occurring sleep and anesthetic-induced unconsciousness.

## Introduction

The state of consciousness is closely tied to thalamocortical oscillations^1^. In states of diminished consciousness such as slow wave (NREM) sleep and anesthesia, EEG exhibits slow (<4 Hz) oscillations associated with rhythmic discharges of cortical and thalamic neurons^1–4^. In contrast, during wakefulness, thalamic and cortical neurons exhibit sustained irregular activity patterns associated with desynchronized EEG^1–3^. Modulation stemming from a network of mutually inhibitory arousal and sleep promoting nuclei in the brain stem, hypothalamus, and the basal forebrain^5–7^ strongly influence the oscillation patterns exhibited by the thalamocortical system.

Anesthesia is distinct from sleep. However, some of the underlying physiological processes under anesthesia and during slow wave sleep are shared. While at high concentrations many inhalational and intravenous anesthetics elicit burst suppression, an activity pattern not typically observed in healthy brains, at intermediate concentrations sufficient to induce unconsciousness, conventional non-dissociative anesthetics induce slow (<4 Hz) EEG rhythms^8^. EEG^9^ and single-neuron activity across cortex and thalamus^10^ under anesthesia-induced slow EEG oscillations resemble NREM sleep, indicating convergence at the level of circuit dynamics.

Corroborating evidence shows that anesthetics activate sleep-promoting and suppress wake-promoting subcortical nuclei^11–18^. There are, however, key distinctions between sleep and anesthesia. For instance, many anesthetics directly reduce the excitability of cortical neurons^19–21^, leading to widespread decrease of cortical activity under anesthesia^22–27^. Direct modulation of biophysical properties of cortical neurons by sleep promoting neuromodulators is complex and cell type specific^28^. However, much like under anesthesia, activity of most cortical neurons is also reduced in NREM sleep^29–31^.

With the notable exception of heat-sensing hypothalamic^32^ and hypothalamic-projecting midbrain^33^ and brainstem^34^ neurons, the vast majority of all identified subcortical NREM-promoting neurons are inhibitory. Likewise, apart from supraoptic^35^ and habenular glutamatergic neurons^36^, the vast majority of neurons known to potentiate the effects of anesthetics are also inhibitory^11,17,18^. Recent studies suggest that the cortex also promotes NREM sleep^37^ and several subtypes of inhibitory cortical NREM-promoting neurons^38–42^ have been identified. Furthermore, zolpidem, a commonly used sleep aid that potentiates inhibitory GABAergic transmission^43^, hastens the onset of sleep in part by acting upon the frontal cortex^44^. Thus, suppression of consciousness by both cortical and subcortical circuits appears to be primarily established through inhibition.

There are, however, compelling reasons to believe that a diminished level of consciousness may be actively brought about by excitatory neurons. For instance, NREM sleep increases following chemogenetic potentiation of excitatory synapses in the prefrontal cortex (PFC)^45–47^. Conversely, disruption of synaptic transmission in Layer 5 (L5) pyramidal neurons disrupts sleep homeostasis and reduces NREM^48^. These results are consistent with in vivo^49,50^, cortical slab^51^, and slice work^52,53^ suggesting that slow oscillations that define states of unconsciousness originate in L5 and are critically dependent upon specific biophysical properties of L5 neurons^54^. L5 neurons, however, are a broad category composed of distinct neuronal subtypes^55,56^ with distinct properties and synaptic connectivity in different cortical regions^57–59^. The specific excitatory cortical circuits that suppress consciousness, however, have yet to be identified.

Here, using targeted recombination in active populations (TRAP)^60^, we identify a specific L5 excitatory neuron subtype in the PFC preferentially activated under anesthesia. Activation of these neurons elicits deep NREM sleep with enhanced slow-wave activity and potentiates anesthetic-induced unconsciousness, whereas their inactivation partially antagonizes anesthetic effects and disrupts NREM sleep. Therefore, we refer them as NREM- and Anesthesia-Promoting (NAP) neurons. Biophysical and transcriptomic analyses classify NAPs as excitatory L5 extratelencephalic (ET) pyramidal neurons, a finding confirmed using a transgenic L5 ET mouse line^61^. We show that mechanistically distinct anesthetics directly increase the excitability of L5-ET PFC neurons and suppress the excitability of a complementary set of intratelencephalic L5 (L5-IT) neurons. Consistent with NAPs being L5-ET neurons, their projections predominantly target subcortical nuclei known to play a key role in controlling of level of consciousness. While previous work identified PFC hypothalamus connection as a key circuit that promotes sleep,^41,62^ we show that NAPs lack strong interactions with the hypothalamus. Rather, we show that L5-ET neurons which target the anterior and reticular thalamic nuclei are sufficient to promote deep NREM sleep. These findings define a novel prefrontal corticothalamic circuit that actively promotes slow-wave activity in both natural and pharmacologically induced unconscious states, providing a mechanistic framework for cortical suppression of consciousness by excitatory neurons.

## Results

### Isoflurane Anesthesia Recruits a Subset of Layer 5 Prefrontal Neurons

To identify neurons active under anesthesia, we performed a whole brain screen for c-Fos expression (Figure 1a), widely used to map antecedent neuronal activity^63^. A steady state isoflurane (iso) dose was chosen to elicit consolidated slow wave activity.^64^ Following optical clearing and immunolabeling for c-Fos,^65–67^ brains were imaged with light-sheet microscopy, enabling detection of individual c-Fos positive neurons (Figure 1a). Automated registration^68,69^ to the Allen reference atlas,^70^ allowed quantification of c-Fos expression in neurons across brain regions^70^. This screen identified numerous subcortical loci (Figure 1b), consistent with previous work showing that anesthetics activate sleep-promoting subcortical structures^12,71–74^. Surprisingly, we also observed a region of c-Fos^+^ neurons localized to the infralimbic (ILA), prelimbic (PL), and anterior cingulate (ACA) regions of the prefrontal cortex (PFC; Figure 1b-c). With the exception of piriform cortex, likely activated by the pungent odor of isoflurane,^75,76^ the rest of the neocortex had sparse c-Fos staining (Figure 1d, PFC 0.1664 ± 0.0669, primary motor cortex (M1) 0.0506 ± 0.0257, and primary somatosensory cortex (S1) 0.0589 ± 0.0304, mean c-Fos^+^ neurons/voxel ± std). Mean PFC c-Fos density did not differ between male and female mice (unpaired two-tailed t-test, t(26)=0.66, p=0.52). The overall fraction of PFC c-Fos^+^ neurons was significantly higher under isoflurane than during wakefulness (Sup. Figure 1, Wake 0.0579 ± 0.0291, Iso 0.1081 ± 0.0405; mean ± std), suggesting that a subset of PFC neurons is preferentially activated under isoflurane anesthesia.

**Figure 1:**
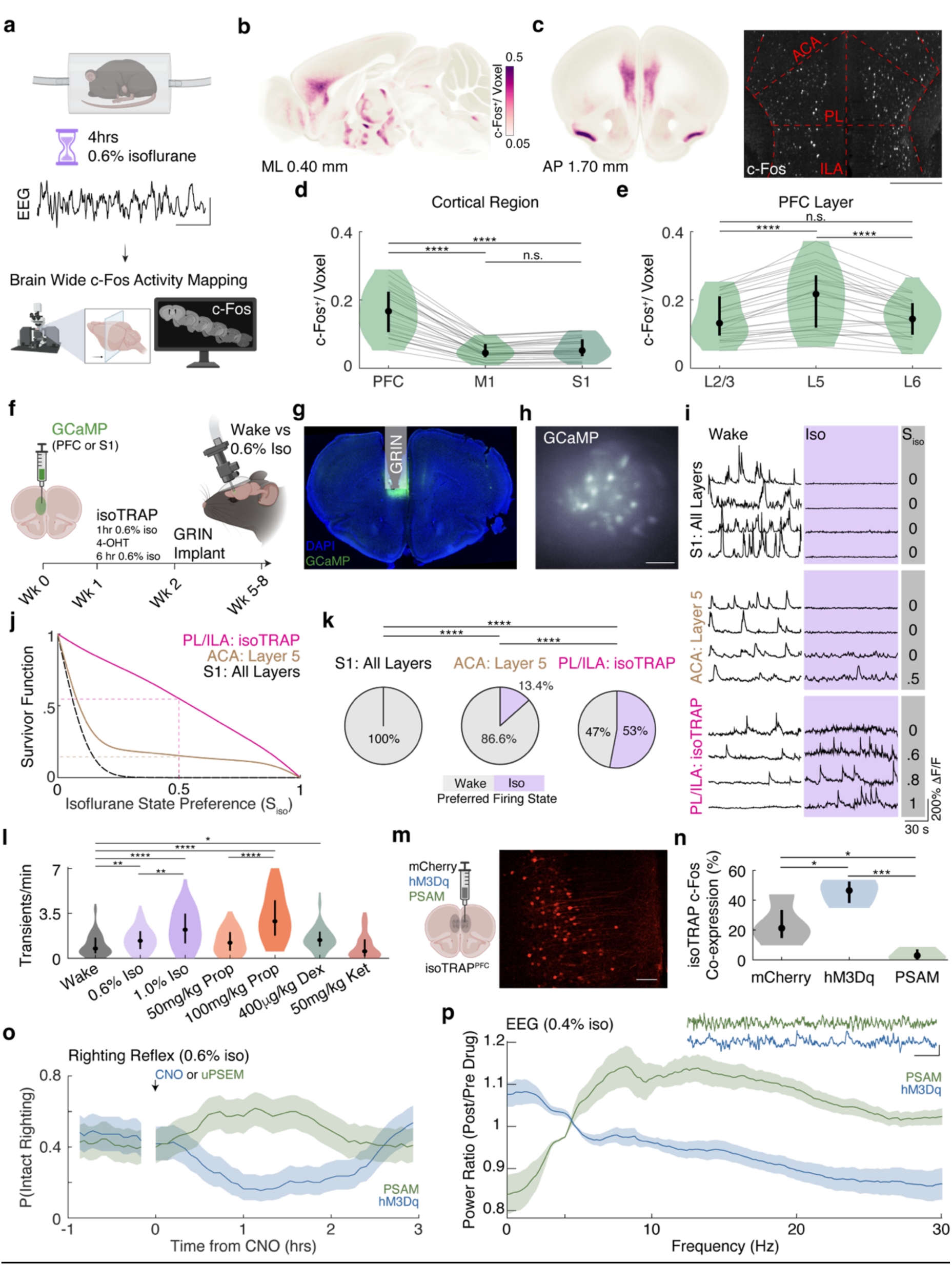
Identification of novel anesthetic active Layer 5 prefrontal cortical neurons that bidirectionally regulate anesthetic sensitivity. **a**) Experimental schematic for whole brain c-Fos activity mapping. Example EEG trace at 0.6% isoflurane. EEG is dominated by δ-oscillations. Scale bar represents 25 μV and 1 sec. **b**) Average sagittal (ML 0.4mm) and **c**) coronal (AP 1.70mm) sections showing c-Fos density (nuclei/voxel) (28 mice (13M 15F), including 10 mice: 5M 5F, from Wasilczuk et al, PNAS 2024). Inset in **c**) shows representative prefrontal coronal optical section (2 μm thick) with approximate regional boundaries for anterior cingulate (ACA), prelimbic (PL), and infralimbic (ILA) cortex indicated with red lines. Scale bar represents 500μm. Distributions of c-Fos densities across different cortical regions (d) and layers of the PFC (**e**) (defined as ILA, PL, ACA) shown as violin plots (gray lines show individual animals, black circle shows the median, vertical bar shows interquartile range). Mean c-Fos density (nuclei/voxel) in the PFC is significantly higher than primary motor (M1) or primary somatosensory (S1) cortex (F(2, 54)=166.0, p<0.0001, repeated measures 1way ANOVA; Tukey’s multiple comparison test: PFC greater than M1, p<0.0001, PFC greater than S1, p<0.0001, M1 no different from S1, p=0.48). Mean PFC density (nuclei/voxel) expression in L5 is significantly higher than L2/3 or L6 (F(2, 54) = 44.48, p<0.0001, repeated measures 1way ANOVA; Tukey’s multiple comparison test: L5 greater than L2/3, p<0.0001, L5 greater than L6, p<0.0001, L2/3 no different from L6, p=0.99). **f**) Experimental schematic and timeline for isoTRAP used for *in vivo* calcium imaging. **g**) Representative localization of GRIN lens in the PL/ILA. **h**) Example miniscope field of view. Scale bar represents 100 μm. **i**) Representative calcium traces (ΔF/F) during wakefulness (white) and under 0.6% isoflurane (purple) in distinct neuronal populations are shown. Gray numbers indicate isoflurane state preference. **j**) Survivor function (P) of Isoflurane State Preference (S_iso_) computed for the three neuronal groups in i. P(0.5) reflects the fraction of neurons that fired at higher frequency under isoflurane (dashed lines). (PL/ILA isoTRAP: 5 mice (3M 2F), 83 neurons; S1 All Layers: 6 mice (3M 3F), 215 neurons; ACA: Layer 5; 4 mice (2M 2F), 119 neurons) **k**) Pie charts showing the fraction of neurons that fired at higher frequency under isoflurane (purple). A chi-square test confirms that PL/ILA isoTRAP had higher preference for firing under isoflurane (χ^2^(2)=136.7, p<0.0001 two sided. Post hoc Fisher’s exact tests (two sided) with Bonferroni correction; S1: All Layers vs ACA: Layer 5, p<0.0001; S1: All Layers vs PL/ILA: isoTRAP, p<0.0001; ACA: Layer 5 vs PL/ILA: isoTRAP, p<0.0001). **l**) PL/ILA isoTRAP calcium activity across multiple anesthetics and doses. Median spontaneous calcium transients of isoTRAP neurons is significantly higher under 0.6% isoflurane (iso), 1.0% iso, 100mg/kg propofol (prop), and 400 μg/kg dexmedetomidine (dex) compared to wakefulness. (H(8)=87.98, p<0.0001, Kruskal Wallis test; Dunn’s multiple comparisons test compared to Wake: 0.6% iso, p=0.0036, 1.0% iso, p<0.0001, 50 mg/kg prop, p=0.99, 100mg/kg prop, p<0.0001, 400 μg/kg dex, p=0.029, 50 mg/kg ketamine (ket), p=0.99). Calcium transient rate was dose dependent for both isoflurane and propofol (1.0% iso greater than 0.6%, p=0.0044; 100mg/kg propofol greater than 50mg/kg propofol, p<0.0001). Wake: 6 mice (4M 2F), 95 neurons; 0.6% iso: 10 mice (5M 5F), 161 neurons; 1.0% iso: 4 mice (2M 2F), 72 neurons; 50 mg/kg propofol: 3 mice (1M 2F) 42 neurons; 100 mg/kg propofol: 4 mice (2M 2F) 32 neurons; 400 μg/kg dexmedetomidine: 3 mice (1M 2F), 51 neurons; 50mg/kg ketamine: 3 mice (1M 2F), 48 neurons. **m**) Experimental schematic for chemogenetics using isoTRAP protocol. Representative section showing viral expression in PL/ILA (right side corresponds to midline; scale bar is 100 μm). **n**) Validation of chemogenetic manipulations. Activation of hM3Dq-expressing neurons increases mean fraction of c-Fos expressing neurons relative to mCherry controls. PSAM-mediated inactivation reduced c-Fos expression relative to mCherry controls. (4 mice (2M 2F) per group, F(2,9) = 17.67, p=0.0008, 1way ANOVA; Tukey’s multiple comparisons test: hM3Dq greater than mCherry, (p=0.0342) and PSAM (p=0.0006), PSAM less than mCherry, p=0.0417). **o**) Population righting reflex (PRP) following PL/ILA: isoTRAP chemogenetic activation (10 mice: 4M 6F) or inhibition (9 mice: 9M) (mean and 95% CI estimated from Binomial distribution). **p**) Power ratios following isoTRAP chemogenetic activation (9 mice:6 M 3F) or inhibition (6 mice: 6F) compared to pre intervention control period. (mean and 95% CI estimated from Jackknife resampling across mice). 30 min data analyzed before and after drug injection [30 min pre & 1-1.5hr post] for ratio calculation. Raw EEG following chemogenetic intervention is also shown. Scale Bar is 25 μV and 1 sec. Statistical significance is denoted by *p<0.05, **p<0.01, ***p<0.001, ****p<0.0001. Schematics from a, f, m were created in BioRender.

PFC c-Fos^+^ neurons were predominantly found in Layer 5 (L5) (Figure 1e, L2/3: 0.1465 ± 0.0614, L5: 0.2051 ± 0.0885, and L6: 0.1477 ± 0.0569 c-Fos^+^ neurons/voxel mean± std). Similarly to the PFC overall, L5 c-Fos density did not differ significantly between male and female mice (unpaired two-tailed t-test, t(26)=0.43, p=0.67). Enrichment of c-Fos in L5 was further confirmed in Rbp4-cre x Ai6 mice, which selectively express ZsGreen fluorescent protein in L5 neurons (Sup. Figure 1 ; Isoflurane 0.06370 ± 0.0304, Wake 0.03636 ± 0.02101 fraction of c-Fos^+^ neurons, mean ± std). Thus, in contrast to most cortical neurons, a subpopulation of Layer 5 neurons in the PFC appears to be preferentially activated under isoflurane anesthesia. Because, as we show subsequently, this specific subset of L5 PFC neurons is **<u>N</u>**REM sleep and **<u>A</u>**nesthesia **<u>P</u>**romoting, we hereafter refer to them as NAPs.

To obtain direct neurophysiological evidence for selective activation of NAPs under anesthesia, we used TRAP^77^ during steady state isoflurane exposure (isoTRAP) to specifically express GCaMP in NAPs (Figure 1f). To avoid capturing neurons activated during induction or emergence from anesthesia, 4-hydroxytamoxifen (4-OHT) was administered 1 hour after the initiation of isoflurane exposure, which was maintained for subsequent 6 hours (Figure 1f). Because most NAPs are located in the PL/ILA—beyond the reach of conventional two-photon imaging —activity of PL/ILA of TRAPed neurons was imaged using a miniature head-mounted microscope via GRIN lens (Figure 1 f-h). For comparison, we also performed pan-neuronal two-photon imaging in S1 (all layers) and in L5 of the ACA (excitatory neurons at least 400 microns from cortical surface).

We confirmed that NAPs were preferentially active under isoflurane anesthesia. In contrast, calcium transients in S1 and ACA L5 neurons were predominantly suppressed (Figure 1i-k). To quantify activation under isoflurane, we computed a state preference index, S_iso_ (Methods). Survivor plots for S_iso_ revealed that more NAPs were preferentially active under isoflurane relative to both S1 and to the excitatory L5 ACA neurons (Figure 1j-k, NAPs: 53%, S1: 0%, L5 ACA: 13.4% of neurons preferentially activated under isoflurane). Activity of ACA GABAergic interneurons (PV: Parvalbumin, SST: somatostatin, VIP: vasoactive intestinal polypeptide) was also preferentially decreased under isoflurane (Sup. Figure 2). Thus, isoTRAP achieved a ∼4-fold enrichment of anesthesia-active neurons relative to L5 ACA. These results confirm that, NAPs are a subpopulation of Layer 5 PFC neurons activated under isoflurane anesthesia.

**Figure 2:**
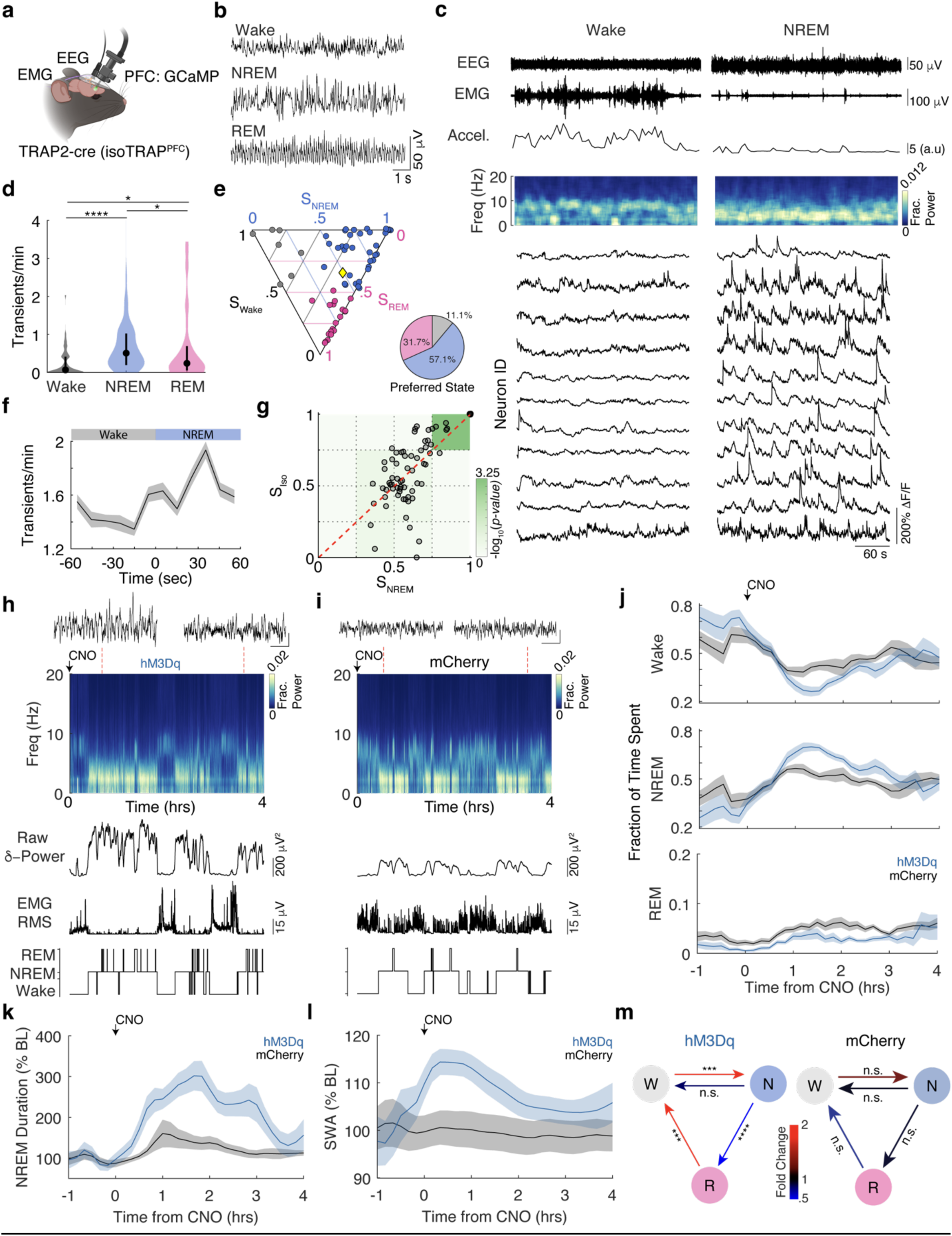
Anesthetic active PFC neurons are preferential active during NREM sleep and drive NREM sleep when chemogenetically activated. **a)** Experimental schematic for simultaneous EEG/EMG and PL/ILA calcium activity recordings. **b)** Representative 10 s EEG recordings classified as Wake, NREM, and REM by a neural network. **c)** 5-minute recordings of EEG, muscle tone, movement, and NAP calcium activity from a representative mouse during wakefulness (left) and NREM sleep (right). Muscle tone and movement were computed as root mean squared (RMS) of EMG and accelerometer signals respectively. EEG spectrograms are expressed as fraction total of power per frequency. **d)** Frequency of calcium transients (transients/minute) during Wake, NREM, and REM sleep shown as violin plot. Calcium transient frequency was highest in NREM sleep (5 mice (3 M 2 F), 63 total cells, χ^2^(2)=30.28, p<0.0001, Friedman test. Post hoc Dunns multiple comparisons test: NREM>Wake, p<0.001; NREM>REM, p=0.0293; REM>Wake, p=0.0131). **e)** Distribution of state preferences shown as simplex plot (dots show individual neurons, colored according to the state in which maximal firing was observed Wake: gray, NREM; blue, or REM: pink). Yellow diamond shows mean of all neurons. Pie chart quantifies proportion of NAPs with maximal activity in each state. Distribution of neurons with state preference was significantly non-uniform (χ²(2) = 10.97, p<0.0041, Chi-Square Test). **f)** Average NAP calcium activity aligned to Wake ◊NREM transition a (mean and 95% CI estimated from Jackknife resampling over Wake◊NREM transitions; 43 transitions from 3 mice (1M 2F). **g)** Relationship between NREM state preference and isoflurane state preference in individual NAP neurons tracked during both natural sleep-wake transitions and under 1.0% isoflurane anesthesia. Each point represents a single NAP neuron (4 mice (2M 2F), 72 neurons; Note: 7 neurons have S_iso_=S_NREM_=1; 1.0 % isoflurane data is reanalyzed from figure 1l). Red line is best fit line (Pearson’s r=0.70, p<0.0001). Permutation analysis (100,000 permutations, bin width 0.25) revealed selective enrichment of neurons with strong preference for both NREM sleep and isoflurane (top right bin, p=0.0005, state preference index ≥0.75 for both measures), with no significant enrichment in any other bin. **h-i)** Spectrogram, delta power, EMG RMS, hypnogram, and zoomed-in EEG traces (from periods marked by red line, scale bar is 25 μV and 1 sec) from a representative hM3Dq (chemogenetic excitation) and mCherry (control) transfected mouse respectively. CNO (3mg/kg, IP) injections in both groups occurred in the middle of the dark phase (ZT 18). **j)** Fraction of time spent in each of the three states (Wake, NREM, REM) following CNO in mCherry (grey): 9 mice (1M 8F) and hM3Dq (blue): 11 mice (7M 4F) cohorts. Two-way ANOVA for the fraction of time spent in NREM (F(4,90)=93.07, p<0.0001) and Wake (F(4,90)=73.75, p<0.0001) before and after CNO revealed statistically significant effects. Šídák’s multiple comparisons test revealed that hM3Dq transfected mice spent greater fraction of time in NREM sleep than the mCherry control one hour after CNO administration, p<0.0001). After 3 hours following CNO administration differences in NREM sleep between the cohorts were no longer observed (p=0.6293). ANOVA performed for the fraction of time spent in REM, did not reveal significant effect (F(4,90)=1.786, p=0.1385). **k)** NREM bout duration following CNO. Two-way ANOVA performed for NREM bout durations (normalized to baseline values in each cohort individually) revealed significant differences between the hM3Dq and mCherry control groups (F(4,90)=104.8, p<0.0001). Šídák’s multiple comparisons test revealed that NREM bout durations in hM3Dq cohort was longer than in mCherry controls 1-2 hours post CNO, p<0.0001). **l)** Slow-wave activity (SWA) following CNO. Two-way ANOVA on SWA (normalized to pre-CNO values in each cohort individually) revealed that SWA differed significantly between hM3Dq and mCherry groups (F(4,90)=28.50, p<0.0001). Šídák’s multiple comparisons confirmed that SWA increased more in hM3Dq than in mCherry control 1-2 hours following CNO, p<0.0001). All data in **j-l** are shown as mean and 95% CI estimated from Jackknife resampling across mice. **m)** Transition probabilities after CNO injection (0.5-2.5 hrs) in hM3Dq (left) and mCherry (right) cohorts were normalized by transition probabilities zeitgeber time–matched (ZT 18.5-20.5) period 24 hrs prior to CNO injection. (7 hM3Dq mice, 8mCherry mice) CNO induced changes in the hM3Dq cohort F (1.504,9.023)=97.41, p<0.0001, 2way ANOVA with Geisser-Greenhouse Correction; Šídák’s multiple comparisons test: W-N, p=0.0004; N-W, p=0.7084; N-R, p<0.0001; R-W, p=0.0003;). No significant difference was found for control and post CNO period in mCherry mice (F (2.599, 18.19) = 0.2672, p=0.8215, 2way ANOVA with Geisser-Greenhouse Correction). Red values indicate a greater transition probability in the CNO condition, whereas blue values indicate a greater probability in the baseline condition. Statistical significance is denoted by *p<0.05, **p<0.01, ***p<0.001, ****p<0.0001. Schematics from a was created in BioRender.

### Slow-Wave Promoting Anesthetics Dose-Dependently Recruit NAPs

Anesthetics exert their effects on the brain via a multitude of molecular mechanisms and differ among distinct anesthetic agents^78^. To determine whether recruitment of NAPs is specific to isoflurane or generalizes across multiple mechanistically distinct anesthetics, we recorded calcium activity from NAPs during exposure to multiple anesthetics (Figure 1l). Relative to wakefulness, NAP activity was significantly increased during isoflurane and propofol exposure as well as during dexmedetomidine-induced sedation. Furthermore, NAP activity increased in a dose-dependent manner for both isoflurane (0.6% vs 1.0%) and propofol (50 vs 100 mg/kg). In contrast, ketamine (50 mg/kg IP), a dissociative agent which minimally affects the amplitude of δ-oscillations^79^, did not significantly alter NAP calcium transient rate relative to wakefulness. These findings demonstrate the *in vivo* dose-dependent activation of NAPs by mechanistically distinct anesthetics and sedatives that enhance slow-wave activity.

### NAPs Bidirectionally Modulate Anesthetic Potency

To test whether NAPs causally modulate anesthetic sensitivity, we selectively expressed chemogenetic modulators (hM3dq: excitatory, PSAM: inhibitory) or control mCherry virus in the PFC of TRAP2-cre mice (Figure 1m). c-Fos immunostaining confirmed that bidirectional chemogenetic modulation was successful in vivo (Figure 1n). Behavioral assays using repeated righting reflex near the EC50 (0.6% isoflurane) dose revealed that hM3Dq activation of NAPs during steady state isoflurane exposure significantly increases anesthetic potency, causing more than a 50% decrease in righting probability. In contrast, PSAM-mediated inhibition of NAPs during steady state isoflurane exposure significantly decreased anesthetic potency (Figure 1o). To determine whether NAP activity also causally contributes to electrophysiological anesthetic endpoints, we exposed mice to steady state 0.4% isoflurane, a dose that produces EEG activity similar to NREM sleep while avoiding burst suppression (Sup. Figure 3) and chemogenetically activated or inhibited them. NAP activation increased the power of EEG slow wave activity and decreased power at higher frequencies. (Figure 1p). Conversely, PSAM-mediated inactivation of NAPs produced the opposite effect on the EEG. Together, these findings demonstrate that NAP activity bidirectionally controls both behavioral and electrophysiological anesthetic endpoints.

**Figure 3:**
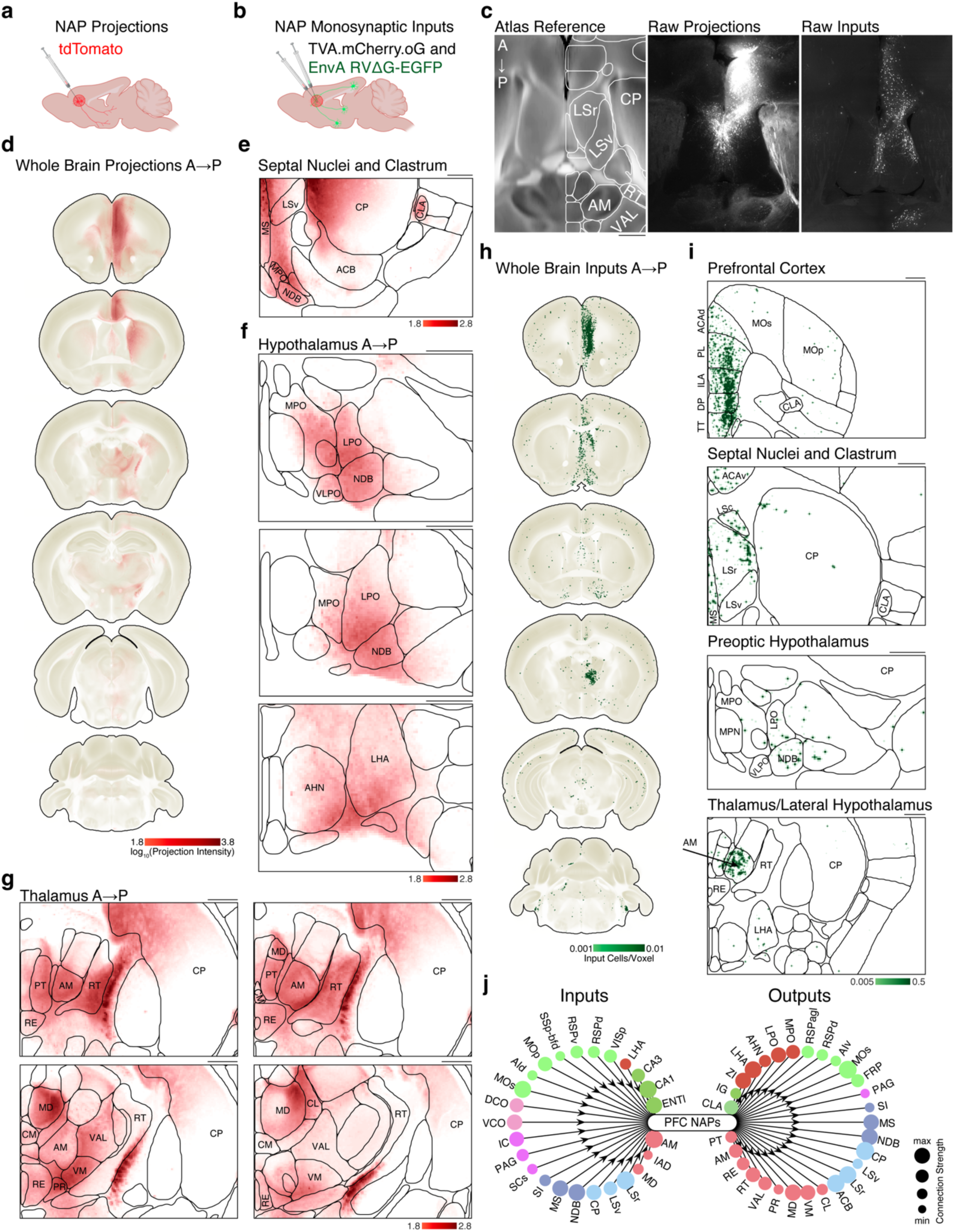
Whole brain mapping of NAP outputs and monosynaptic inputs. **a)** Experimental schematic for anterograde tracing outputs of NAPs (3 mice, 3M). **b)** Experimental schematic for tracing monosynaptic inputs into NAPs (4 mice, 4F). **c)** Transverse section through the reference atlas with overlaid summary structure outlines (left), representative transverse slices (20μm maximal projection) showing anterograde projections (middle), and monosynaptic inputs (right) highlighting bidirectional communication of NAPs with anteromedial nucleus (AM) and projections to the reticular nucleus (RT). **d)** Coronal sections of the average (n=3 mice) brain-wide distribution of NAP projections **(e–g)** Region-specific insets of NAP projections to septal nuclei and claustrum **(e)**, hypothalamus **(f)**, and thalamus **(g)**. Colormap **(d-g)** shows log_10_(projection intensity per voxel). **h)** Coronal sections of the average (n=4 mice) brain-wide distribution of monosynaptic inputs into NAPs. **i)** Region-specific insets highlight inputs from the PFC, septal nuclei and claustrum, preoptic hypothalamus, and thalamus/lateral hypothalamus. Colormap **(h-i)** shows the density of inputs (cells/voxel). Outlines in e, f, g, and i represent boundaries of relevant summary structures as defined by the Allen reference atlas. Fiber tracts and ventricles are not outlined. **j)** Connectivity plots summarizing relative strength of strongest (top 10%) inputs to (left) and outputs of (right) of NAPs. Node size encodes connection strength (scaled to the maximum value separately for inputs and outputs). Scale bars represent 500 μm across all subpanels. Schematics from a, b were created in BioRender.

### NAPs are Preferentially Active During NREM Sleep

To test if NAPs also participate in natural sleep, we recorded their calcium activity in conjunction with EEG/EMG recordings in freely moving mice across naturally occurring sleep and wake states (Figure 2a) in undisturbed, home cage conditions. Vigilance states were classified as Wake, NREM, or REM (Figure 2b) using a custom neural network classifier (Sup. Figure 4). NAPs were largely silent during periods of consolidated wakefulness (Figure 2c), but displayed large, coordinated calcium transients during NREM sleep. Some NAPs also exhibited increased activity during REM sleep (Sup. Figure 5). NAPs exhibited highest calcium transient rates during NREM sleep (Figure 2d). Median calcium transient rates during NREM sleep were significantly higher than REM (2.1-fold difference) and wakefulness (7.5-fold difference). NAP activity in REM was higher than wakefulness (3.6-fold difference). The distribution of NAP state preferences, visualized as a simplex plot, was non-uniform (χ²(2)=10.97, p=0.0041, Chi-square Test), with a strong bias toward NREM sleep (Figure 2e). Further, NAP calcium activity increased at the onset of NREM and progressively grew peaking at ∼30 seconds after NREM onset (Figure 2f). Thus, NAP calcium activity is also preferentially recruited into NREM sleep.

**Figure 4:**
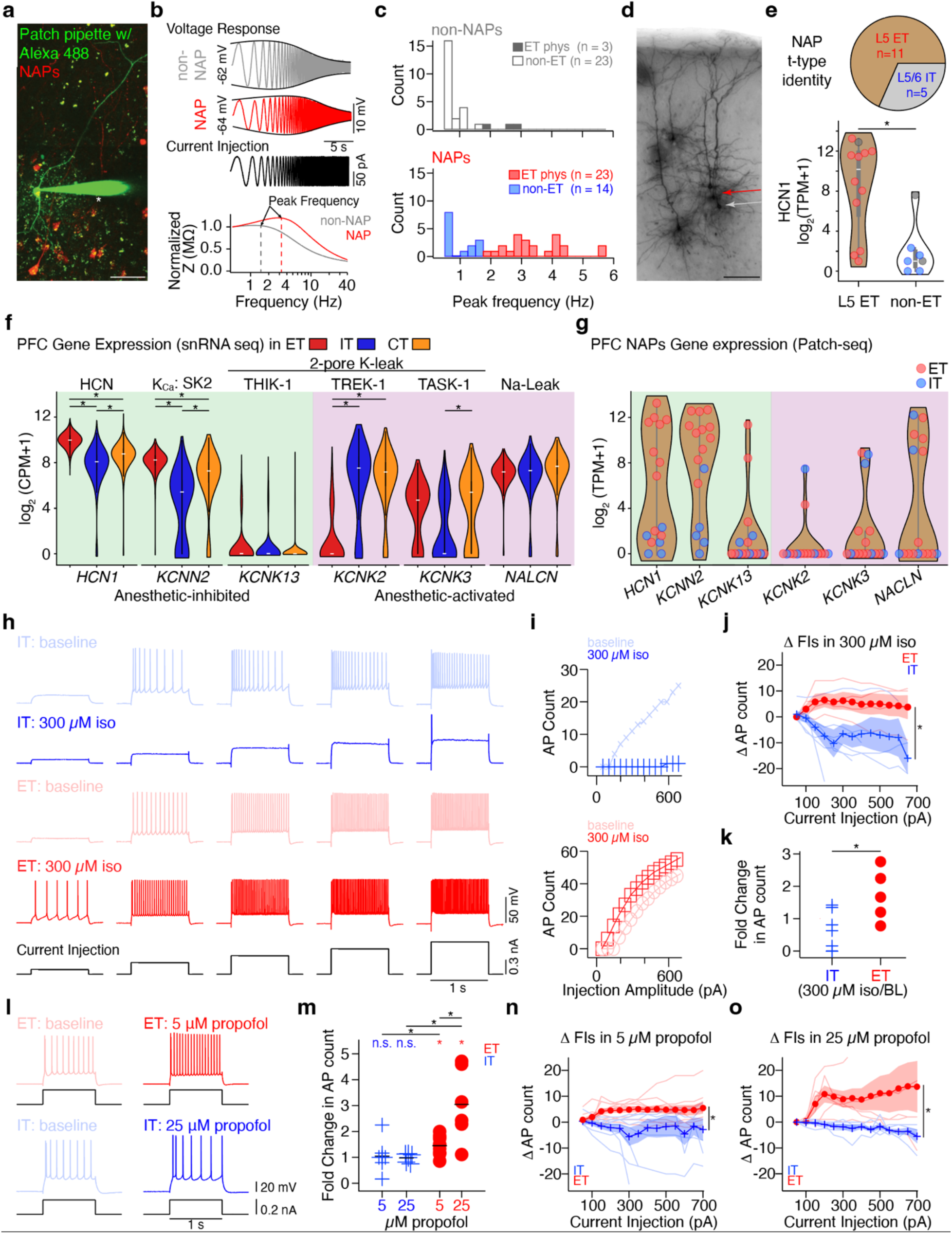
NAPs are enriched in a molecularly and physiologically defined L5 ET cell type in PFC whose excitability is directly enhanced by isoflurane and propofol. **a)** 2-photon Z-stack of NAPs (red) in medial prefrontal cortex, with patch recording (electrode marked with *) from a labeled L5 pyramidal neuron filled with Alexa 488 (green). Neurons lacking red fluorescence were classified as non-NAPs. Scale bar: 100 μm. **b)** Example voltage response of a NAP (red, cell shown in a) and a non-NAP (grey) neuron to sinusoidal chirp current injection (black). Below, normalized impedance (Z) amplitude profile calculated from the voltage responses above; dashed lines indicate peak resonant frequency. **c)** Histogram of the distribution of peak resonant frequency for non-NAP (top, grey) and NAP (bottom, red) neurons. Frequencies above 2 Hz (dashed line) were classified as ET-like physiology (red, filled bars); frequencies below were classified as non-ET (blue, open bars). The proportion of neurons with ET-like physiology differed significantly between groups (χ²(1) = 16.15, p < 0.0001). **d)** Pyramidal neuron morphology of NAPs and non-NAPs, revealed by histological processing of neurobiotin fills. Arrows denote the neurons recorded in (a, b); red = NAP, grey = non-NAP. Scale bar: 100 μm. **e)** Patch-seq validation that NAPs are primarily ET neurons (top). Molecular subclass identity of NAP nuclei (n = 16), aligned to the Allen Institute taxonomy. Physiological and molecular classification of L5 ET neurons matched 100% (n = 16). Bottom, violin plots of HCN1 transcript log2 transcripts per million (TPM+1) in non-labeled neurons (n = 3, grey) and NAPs (n = 16), split into molecularly and physiologically defined L5 ET (red, n = 9) and L5-6 IT-like (blue, n = 7) groups. Mean HCN1 transcript counts were significantly higher in L5 ET neurons (unpaired t-test, t = 2.49, p = 0.026, d = 1.22). **f)** Violin plots comparing gene expression across PFC L5 ET (n = 1656), L4-6 IT (n = 11,301), and L6 CT (n = 7425) nuclei from the Allen Institute mouse brain dataset^55^ for key ion channels inhibited (left, green box) or activated (right, purple box) by anesthetics. Cell type specific expression was compared using two-sided Wilcoxon rank-sum test with a Benjamini-Hochberg false discovery rate correction done within each pairwise comparison. Differences labeled as statistically significant had an adjusted p value < 0.05 and effect size (r) of at least 0.3. **g)** Violin plots of the same ion channel genes from patch-seq recordings of NAPs (n = 16), showing broad concordance with the snRNA-seq data in (f). **h)** Example voltage traces from an L5 IT neuron (top) in response to 1 s depolarizing current injections (Δ150 pA, black traces, bottom) at baseline (light blue) and after bath perfusion of 300 μM isoflurane (blue). Below, voltage responses from an example L5 ET neuron at baseline (light red) and in isoflurane (red). **i)** Frequency-current (F-I) plots for the representative IT (top, blue) and ET (bottom, red) neurons, showing action potential (AP) count in response to 1 s depolarizing current steps (50–700 pA, 50 pA intervals) at baseline and in isoflurane. **j)** Average change in F-I (ΔFI) caused by 300 μM isoflurane in ET (red circles, n = 4) versus IT (blue crosses, n = 6) neurons was statistically significantly different (linear mixed-effects model, LMM, fit by restricted maximum likelihood, p<0.01). Shading indicates SEM; light traces show individual cells. **k)** Fold change in AP count caused by 300 μM isoflurane in ET (n=5) versus IT (n=6) neurons, measured using a fixed current injection sufficient to drive 5–9 APs at baseline. **l)** Example responses to propofol: 5 μM onto an ET neuron (top, red) and 25 μM onto an IT neuron (bottom, blue). **m)** As in (k), fold change in AP count caused by propofol for IT (blue crosses, 5 μM: n = 8; 25 μM: n = 6) and ET (red circles, 5 μM: n = 11; 25 μM: n = 6) neurons. Both propofol concentrations significantly increased AP count from baseline in ET but not IT neurons (Wilcoxon Signed test, p = <0.05); Changes in ET excitability differed significantly from IT at both concentrations (Wilcoxon rank sum, p < 0.05) and were greater at 25 μM than at 5 μM (Wilcoxon Rank test, p = <0.05). **n, o)** As in (j), average ΔFI caused by (n) 5 μM and (o) 25 μM propofol; both concentrations produced significantly different responses between ET and IT neurons (LMM fit by restricted maximum likelihood, p<0.01). Significance: *p < 0.05, ***p < 0.0001.

**Figure 5:**
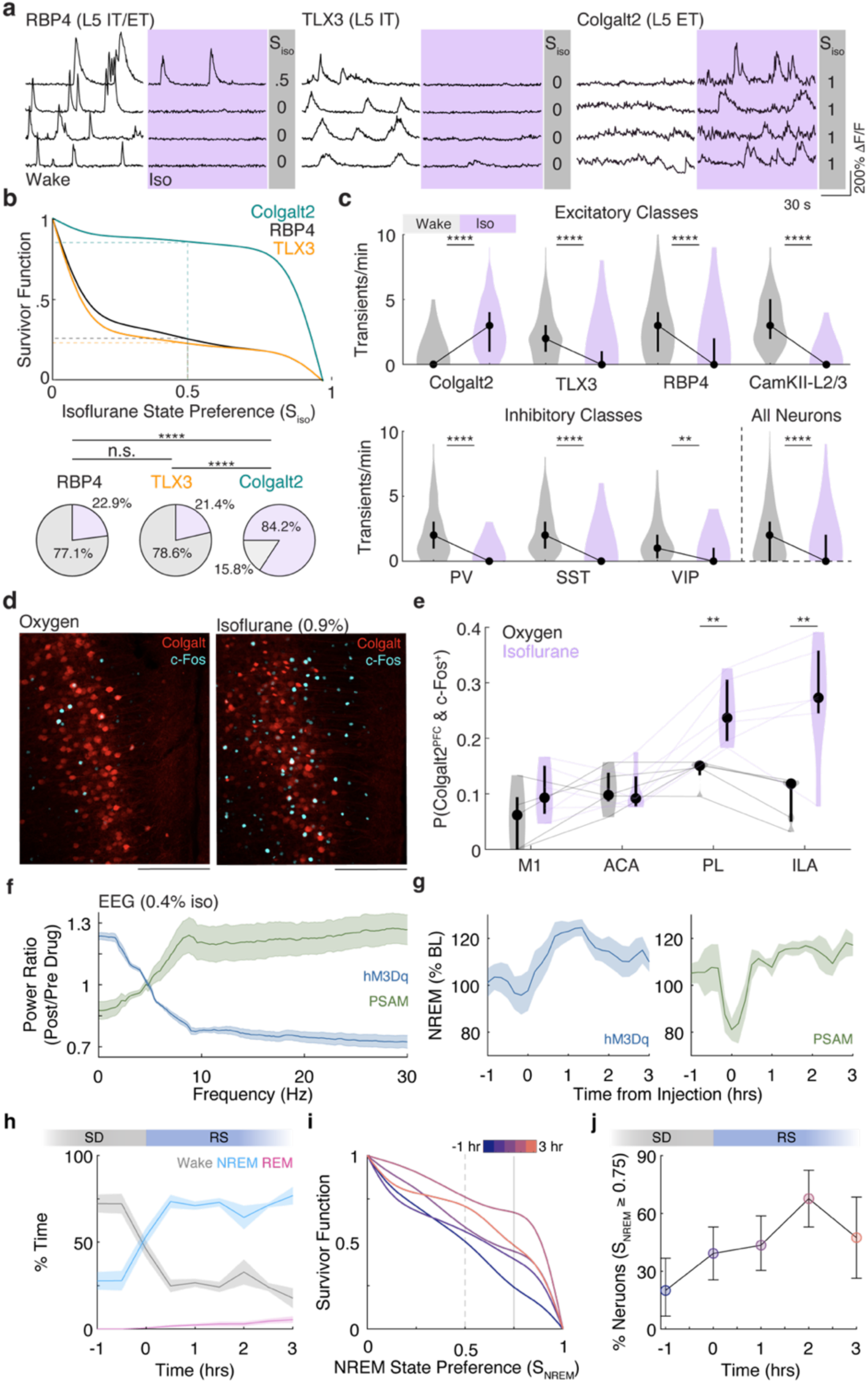
Prefrontal Colgalt2-cre neurons are recruited during anesthesia and sleep and bidirectionally modulate brain state. **a)** Representative two-photon calcium activity traces recorded from cre-driver lines that label distinct Layer 5 subtypes in the ACA during wakefulness and under 0.6 % isoflurane (purple). Numbers shown at right margin (gray) indicate Isoflurane State Preference (S_iso_) for each neuron (RBP4 4 mice (2M 2F), 170 neurons; TLX3 4 mice (3M 1F), 192 neurons; Cogalt2 4 mice (4 M), 190 neurons). **b)** Survivor function (P) of S_iso_ for all neurons in L5 subtypes. P(0.5) reflects the fraction of neurons that had higher activity under isoflurane (dashed lines) shown also as pie charts (purple: isoflurane; gray: wake). Isoflurane preference differed significantly between the L5 subtypes (χ^2^(2)=195.7, p<0.0001. Post hoc Fisher’s exact tests with Bonferroni correction; RBP4 vs TLX3, p=0.7998; RBP4 vs Colgalt2, p<0.0001; TLX3 vs Colgalt2, p<0.0001). **c)** Calcium transient rates across major excitatory (top), inhibitory (bottom) neuron subtypes. Colgalt2 is the only molecularly defined subtype that increases calcium activity under anesthesia (Wilcoxon signed-rank test: Colgalt2 Wake < Iso, p<0.0001; TLX3 Wake > Iso, p<0.0001; RBP4 Wake > Iso, p<0.0001; CamKII-L2/3 Wake > Iso, p<0.0001; PV Wake > Iso, p<0.0001; SST Wake > Iso, p<0.0001; VIP Wake > Iso, p=0.0039; All PFC Wake > Iso, p<0.0001). Furthermore, Colgalt2 neurons were significantly more active under isoflurane than all other neuron subtypes. (H=277.8, df=8, p<0.0001, Kruskal-Wallis test. Dunn’s Multiple Comparisons Test: Colgalt2 vs TLX3, RBP4, CaMKII-L2/3, PV, SST, VIP, All PFC, all p<0.0001). **d)** Representative images of tdTomato-labeled Colgalt2-cre PL/ILA neurons (red) and c-Fos immunoreactivity (cyan) in mice exposed to oxygen (top, 5 mice (5M)) or 0.9% isoflurane (bottom, 6 mice (6M)). Scale bars represent 200 μm. **e)** Proportion of c-Fos^+^ Colgalt2 neurons (tdTomato^+^) across PFC subdivisions in oxygen (gray) or isoflurane (purple) conditions. Two-way ANOVA revealed significant effects of condition (F(1,9)=13.02, p=0.0057) and region (F(1.47,13.23)=12.66, p=0.0016), as well as a significant condition x region interaction (F(1.47, 13.23)=6.216, p=0.018). Post hoc Šidák comparisons revealed significantly increased c-Fos expression following isoflurane in the infralimbic (p=0.0083) and prelimbic cortex (p=0.0045), whereas no significant differences between oxygen and isoflurane conditions were observed in M1 (p=0.181) or ACA (p=0.972). **f)** Power ratios following Colgalt2-cre prefrontal chemogenetic activation (6 mice, 6M) or inhibition (6 mice, 6M, 0.4 % isoflurane, 30 min pre & 1-1.5hr post CNO or uPSEM for ratio calculation; mean and 95% CI estimated from Jackknife resampling across mice). **g)** Fraction of time spent in NREM sleep following PFC chemogenetic activation (CNO in hM3Dq-expressing Colgalt2 mice, 9 mice (9M), injection at ZT 18, left) or chemogenetic inhibition (uPSEM in PSAM-expressing Colgalt2 mice,6 mice (6M), injection at ZT 6, right). Chemogenetic activation increased NREM sleep: (Repeated measures one-way ANOVA: F(2,16)=4.115, p=0.036) but not REM sleep (F(2,16)=0.71, p=0.51), with post hoc Dunnett’s multiple comparisons revealing NREM sleep significantly increases 1-2 hours post CNO administration (p=0.021) and returning to baseline level after 3 hours (p=0.29). Conversely, chemogenetic inhibition decreased NREM sleep (Repeated measures one-way ANOVA: F(4,20)=5.067, p=0.0055) and REM sleep (F(4,20)=3.057, p=0.041). Dunnett’s multiple comparisons test showed that NREM sleep decreased during the first 30 min (p=0.043) following UPSEM administration and returned to baseline levels after 1 hour (p>0.05). Mean and 95% CI estimated from Jackknife resampling across mice normalized to pre injection period are shown. **h)** Fraction of time spent in Wake (gray), NREM (blue), and REM (pink) during novel object induced partial sleep deprivation (lasting 4 hours, last hour shown) and subsequent recovery in mice expressing GCaMP in prefrontal Colgalt2 neurons and implanted with miniscope targeting PL/ILA (6 mice (6M); mean and 95% CI estimated from Jackknife resampling across mice for each group shown). **i)** Survivor function of S_NREM_ as a function of time relative to termination of the sleep deprivation phase. **j)** Percentage of Colgalt2 neurons with NREM state preference greater than 0.75 as a function of time as in **i**. Fraction of Colgalt2 PL/ILA neurons with strong NREM preference was non-uniform sleep deprivation and recovery sleep (χ^2^(4)=15.32, p<0.0041). (Fraction of neurons and 95% CI estimated from Binomial distribution is shown). Statistical significance is denoted by **p<0.01 and ****p<0.0001

### NAPs are Jointly Recruited During NREM Sleep and Anesthesia

To determine if the same individual NAP neurons are preferentially recruited during both NREM sleep and anesthesia, we tracked a subset of NAPs across sleep-wake states and during 1.0% isoflurane anesthesia without altering the imaging plane (Figure 2g). The NREM and isoflurane state preference indices were strongly correlated, indicating that neurons active during NREM sleep tended to also be active during isoflurane anesthesia. To further quantify this, we generated a two-dimensional histogram of NREM and isoflurane state preference indices and compared the occupancy of each bin to a permutation-derived null distribution. This analysis revealed statistically significant enrichment of neurons with strong preference for both NREM sleep and isoflurane anesthesia (state preference index greater than 0.75 for both measurements). Thus, while NREM sleep and isoflurane anesthesia are distinct physiological and behavioral states, our results suggest that NAPs are selectively recruited into both.

### NAPs Promote NREM Sleep

We next asked whether NAP activity causally contributes to sleep. Cre-dependent hM3Dq or mCherry control viruses were injected into the PFC of TRAP2-cre mice and recombination was induced using the isoTRAP protocol (Figure 2h-i). Vigilance states were recorded as above for 24 hours prior to injection of the hM3Dq agonist clozapine-N-oxide (CNO) in the middle of the active period (Zeitgeber Time 18). Chemogenetic activation of NAPs selectively promoted NREM over wakefulness (Figure 2j). NAP activation significantly decreased Wake time and increased NREM sleep compared to their own baseline and to the mCherry controls, suggesting that CNO effect were mediated by hM3Dq rather than potential off target effects^74,80–82^. While there is no clear REM equivalent under isoflurane anesthesia, REM sleep was observed following NAP activation (Sup. Figure 6) and followed bouts of NREM sleep. REM was modestly increased in both hM3Dq and mCherry groups but was not significantly different between them, suggesting a specific effect of NAP activation on NREM. Chemogenetically induced NREM sleep did not appear localized to the PFC as sleep-state classification remained highly concordant across the anterior-posterior axis (Sup. Figure 7). Together, these findings indicate that NAP activation selectively promotes NREM sleep while suppressing wakefulness.

**Figure 6:**
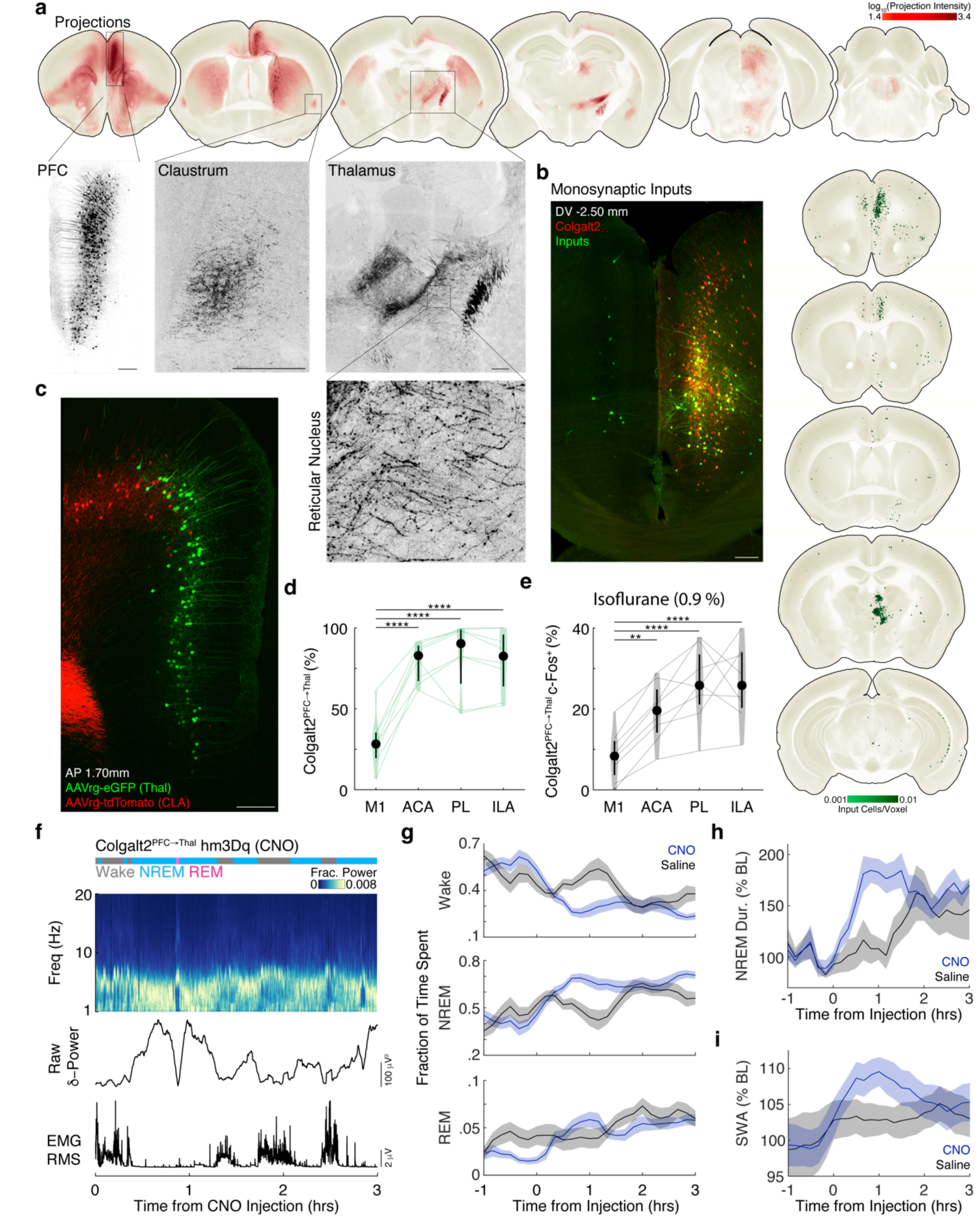
Thalamus-projecting Colgalt2 PFC neurons are NAPs. **a)** Coronal sections of the mean (3 mice (3M)) brain-wide distribution of tdTomato-labeled projection intensity of PFC Colgalt2 neurons. Representative confocal images shown for the PFC (injection site), claustrum, and thalamus; two major projection targets that overlap with the NAP projection network. An enlarged view of projections within the thalamic reticular nucleus is also shown. Colormap shows log_10_(projection intensity per voxel). **b)** Representative transverse light-sheet image (left) through infralimbic and prelimbic cortices showing Colgalt2 starter neurons expressing helper virus (red) and monosynaptic inputs (green). Mean brain-wide distribution of monosynaptic inputs into prefrontal Colgalt2 neurons (right, 3 (3M)). Colormap shows the density of inputs (cells/voxel). **c)** Representative confocal image showing retrograde labeling of Colgalt2 neurons projecting to the claustrum (red) or thalamus (green). **d)** Fraction of all labeled Colgalt2 neurons with thalamic projections across PFC subdivisions (8 mice (8M)). Repeated-measures one-way ANOVA (F(3,21)=36.42, p<0.0001) followed by Tukey’s post hoc comparison shows that thalamus projecting neurons are enriched in the ACA, PL, and ILA relative to M1 (all p<0.0001). **e)** Fraction of c-Fos^+^ Colgalt2 thalamic-projecting neurons after 2-hour 0.9% isoflurane. Repeated-measures one-way ANOVA (F(3,21)=20.40, p<0.0001) followed by Tukey’s post hoc comparison shows enrichment of c-Fos in thalamus projecting neurons in ACA (p=0.0027), PL (p<0.0001), and ILA (p<0.0001) relative to M1. **f)** Hypnogram, spectrogram, delta power, and EMG RMS from an example Colgalt2-cre mouse following chemogenetic activation of thalamus-projecting Colgalt2 PFC neurons. **g)** Fraction of time spent in Wake, NREM and REM following CNO (8 mice (8M)) or saline (7 mice (7M)). Time spent in NREM significantly increased following CNO compared to saline up to 1.5 hours following injection (Repeated-measures two-way ANOVA, F(4,24)=4.21, p=0.01). **h)** NREM bout duration and **i)** SWA were significantly higher following CNO compared to saline. **g-i)** mean and 95% CI estimated from Jackknife resampling across mice shown. Scale bar represents 200 μm (**a-c**).

### NAPs Enhance NREM Stability

We next examined how NAP activation changes the characteristics of NREM and the dynamics of fluctuations between states of sleep and wakefulness. Chemogenetic excitation of NAPs significantly increased NREM bout duration relative to baseline and to the mCherry cohort (Figure 2k). The number of both NREM and REM bouts following NAP activation was reduced relative to mCherry controls (2-way repeated measures ANOVA; main effect: F(1,18) = 8.753, p=0.0084, NREM: 6.55 ± 3.50 vs. 10.00 ± 3.81; REM: 2.00 ± 1.18 vs. 4.89 ± 2.52; bouts/hr; hM3Dq vs. mCherry; mean ± std). Altogether, these results suggest that the increase in total sleep time was due to the prolongation of NREM sleep bouts.

Consistent with deeper, more consolidated sleep, slow-wave activity (SWA) during NREM was also significantly elevated following NAP activation (Figure 2l). In addition, NAP activation significantly reduced locomotor activity (Sup. Figure 8) and decreased body temperature^83^ (hM3Dq Δ_temp_ -0.81 ± 0.12 °C, mCherry Δ_temp_ - 0.01 ± 0.01 °C; mean ± SD; p < 0.0001, unpaired t-test). Thus, NAP activation is associated with behavioral, electrophysiological, and thermoregulatory signatures of deep NREM sleep.

To further quantify the dynamics of sleep-state transitions, Wake, NREM, and REM transitions were modelled as a Markov process. Transition probability matrices were estimated 0.5-2.5 hours following CNO injection and compared to ZT-matched times from the previous day. NAP activation increased the likelihood of transitioning from Wake to NREM and reduced transitions from NREM to REM. (Figure 2m). These changes in transition probabilities serve to stabilize NREM sleep. In contrast, mCherry controls showed no significant changes in transition probabilities following CNO injection. Thus, NAP activation leads to deep NREM sleep characterized by prolonged bout duration and increased slow wave activity.

### NAPs Connect with Subcortical Structures Involved in Control of Arousal and Sleep

The PFC is a highly heterogenous structure, projecting to and receiving inputs from many brain regions.^84–86^ Moreover, PFC connectivity is cell-type dependent.^59,87^ To characterize NAP connectivity, we performed brain-wide anterograde projection and monosynaptic input tracing to identify their downstream targets and upstream inputs (Figure 3).

Although the PFC is broadly connected with other cortical areas,^88^ NAP projections were largely restricted to subcortical regions, with the notable exception of the retrosplenial cortex (Figure 3d, Supplemental Table 1). NAPs innervated subcortical structures known to be involved in sleep and anesthesia including the claustrum^89–94^, the nucleus of the diagonal band^95^, and the lateral and preoptic hypothalamus^11,16–18,96^ (Figure 3e-f). NAPs also projected strongly to anterior thalamic nuclei, with minimal labeling of posterior, sensory-related thalamic regions. Within the anterior thalamus, dense labelling was observed in the reticular nucleus, anteromedial, ventromedial and intralaminar nuclei (Figure 3g)—regions critical for the generation of slow waves during sleep and anesthesia.^97–104^

Replication-deficient rabies monosynaptic input tracing^105^ revealed that the majority of NAP inputs were local L5 PFC neurons, with sparse cortical afferents from retrosplenial, motor, and somatosensory areas (Figure 3h-I, Supplemental Table 1). The most prominent subcortical inputs came from anterior thalamic structures, particularly the anteromedial thalamus (Figure 3c-i). Weaker inputs were observed from the septal nuclei, nucleus of diagonal band, lateral hypothalamus, and CA1 region of the hippocampus (Figure 3i).

### NAPs are enriched in a molecularly and physiologically defined L5 ET cell type

The PFC consists of numerous distinct cell types distinguished by their biophysical properties and synaptic connectivity.^57,56,106^ The specific recruitment of NAP activity during NREM sleep and under anesthesia relative to the nonspecifically labeled L5 PFC neurons raised a possibility that NAPs represent a distinct L5 subtype. Morphological features of NAPs suggested that they are pyramidal neurons (Figure 1n, 4a) and their projections are largely subcortical (Figure 3). L5 pyramidal neurons are classified into two broad classes based on whether their projections are contained within the telencephalon (IT) or extend beyond it, referred to as extratelencephalic (ET). IT and ET neurons are also distinguished by their biophysical properties^106–110^. A key physiological property that distinguishes L5 ET neurons from L5 IT neurons is their somatic subthreshold resonance frequency within the high δ–frequency band (>2 Hz).^107,111^ To determine whether NAPs are enriched in a specific biophysically defined L5 neuronal subtype, we performed targeted whole cell patch clamp recordings in ex vivo slices under quiescent, controlled conditions in the presence of synaptic blockers.

Consistent with our anatomical data showing that NAPs have extratelecephalic projections, most NAPs were enriched in δ-resonance (23/37) compared to a much smaller proportion of unlabeled L5 pyramidal neurons (3/26) which were classified as IT based on their physiological properties (Figure 4b-c). The somatic resonance profile of ET neurons is caused by hyperpolarization activated cyclic nucleotide gated (HCN1) channels^107,112^. In addition to impedance profiles consistent with greater HCN channel expression (Sup. Figure 9a-f), most NAPs also had other hallmarks of HCN channels as demonstrated by their response to step current injections: including larger voltage ‘sag’ and rebound responses (Sup. Figure 9g-l). To assess their excitability, we examined the suprathreshold firing properties of NAPs using depolarizing current injections (Sup. Figure 9m-v). The initial interspike interval (ISI) was frequently shorter in NAPs compared to neighboring unlabeled neurons, consistent with previous observations of onset doublets in L5 ET neurons in other neocortical brain regions.^113^ Together these data indicate that NAPs are enriched in L5 ET-like physiological properties.

To validate our cell-type categorization using the intrinsic biophysical properties, we collected the nuclei in a subset of NAPs (n=16) and performed single-cell RNA sequencing to map their transcriptomic subclass. NAP nuclei were mapped using correlation-based clustering to Allen Institute cell type taxonomy. Consistent with the physiological data, most NAP nuclei were molecularly defined as the L5 ET subclass (11/16) (Figure 4e). The remainder of nuclei mapped to the L5-6 IT subclass (5/16). No nuclei mapped to the near-projecting (NP), or L6 CT or L6b molecular subclasses. Given the ∼35% prevalence of L5 ET transcriptomic subclass in infragranular layers of ventral PFC,^55^ the probability of observing the L5 ET subclass in 11 out of 16 NAPs by chance is 0.0049. Consistent with HCN dependent properties being a defining feature of L5 ET neurons, HCN1 gene expression was higher in neurons with L5 ET-physiological properties compared to non-ET neurons (Figure 4e). The concordance between physiological cell type classification (ET versus IT) and molecular mapping was 100% (n = 16). Thus, converging gene expression and neurophysiological data indicate that PFC neurons isoTRAPed under isoflurane anesthesia are predominantly L5 ET neurons. Remarkably, while NAPs are engaged in strong bidirectional interactions with the thalamus, not a single NAP was mapped to the Layer 6 corticothalamic (CT) subtype.

### L5 ET excitability is enhanced by two molecularly distinct anesthetics

Having determined that NAPs are primarily comprised of the L5 ET neuron subclass, we sought to identify potential molecular mechanisms through which anesthetics may directly act upon NAPs to contribute to their selective recruitment under anesthesia. While anesthetics are well known to act upon synaptic transmission, many ion channels are also direct molecular targets of anesthetics^114–116^. Thus, we first asked whether known anesthetic-targeted channels were differentially expressed across subclasses of PFC pyramidal neurons. We leveraged a publicly available single-nucleus gene expression dataset^55^ to compare gene expression across L4-6 IT, L6 CT, and L5 ET neuron types in the PFC (Figure 4f), as well as in NAPs from our patch-seq data (Figure 4g). Anesthetics generally suppress neuronal activity; however, the direct actions of anesthetics on several ion channel targets can be pro-excitatory. For example, both isoflurane^117,118^ and propofol^119–121^ inhibit HCN1 channels, whose gene expression is significantly enriched in PFC L5 ET neurons compared to IT and CT types (Mann-Whitney U, r = 0.379 for ET vs IT; r = 0.374 for ET vs CT, both with a Benjamini-Hochberg adjusted p < 0.001). HCN inhibition has complex effects on excitability by hyperpolarizing neurons while simultaneously increasing their input resistance, making them more responsive to synaptic inputs^107,109^. Other ion channels targeted by anesthetics also have pro-excitatory effects. For example, inhibition of Ca2+-activated potassium (SK2) channels^122^, two-pore domain Halothane-Inhibited K+ (THIK-1) channels^123^, and potentiation of Na+ leak current (NALCN)^124^ can also increase excitability. Among these pro-excitatory targets, *KCNN2* (SK2, inhibited by anesthetics) gene expression is significantly higher in PFC ET neurons relative to IT and CT neurons (Figure 4f, Mann-Whitney U, r = 0.374 for ET vs IT; r = 0.252 for ET vs CT, both with a Benjamini-Hochberg adjusted p < 0.001). In addition to comparing pro-excitatory targets of anesthetics, we also compared gene expression of several anesthetic-activated two-pore potassium channels that inhibit neuron excitability (Figure 4f, Sup. Figure 10a-b). Among these *KCNK2* (TREK-1) expression was significantly lower in L5 ET neurons (Mann-Whitney U, r = 0.297 for ET vs IT; r = 0.371 for ET vs CT, both with a Benjamini-Hochberg adjusted p < 0.001).

Given the differences in expression of anesthetic-targeted ion channel genes among the major subtypes of PFC pyramidal neurons, we sought to determine whether anesthetics differentially effect the excitability of PFC ET versus IT neurons. Because our physiological classification of IT vs ET neurons was perfectly aligned with single nucleus gene expression based neuron subtype identification (Figure 4b-e), in the following experiments we relied solely on the physiological criteria to distinguish between IT and ET neurons. To specifically isolate the direct effects of anesthetics, experiments were performed in the presence of a cocktail of synaptic receptor blockers, conditions under which PFC neurons are otherwise quiescent. We delivered 300 μM isoflurane (see Methods for details on calibration and delivery of this volatile anesthetic) and tested for changes in excitability measured as the number of action potentials in response to the depolarizing current injections. This isoflurane concentration is similar to that measured in the brains of mice anesthetized with 1% isoflurane^74^. Interestingly, the excitability of IT and ET neurons was indeed differentially affected by isoflurane. While the excitability of IT neurons was largely suppressed under isoflurane relative to baseline, ET neurons were generally rendered more excitable (Figure 4h-k). The suppressive effect of 300 μM isoflurane on IT neurons was most pronounced at larger depolarizing current injections (Figure 4i-j), during which most IT neurons failed to sustain action potentials over the 1s depolarization. In contrast, the increase in excitability in ET neurons occurred across the entire range of current injections (Figure 4i-j). Given that we had also observed NAP activation in vivo under propofol induced anesthesia (Figure 1l), we next tested whether propofol would similarly exhibit cell type specific effects on PFC ET versus IT neurons. As we had observed with isoflurane, the effect of propofol on excitability depended upon cell type (drug/baseline ratio values, N=31; F_(1,27)_ = 9.96, compared to 17/10,000 permutations, p = 0.0017). Propofol made ET neurons more excitable (Figure 4l-o) to depolarizing steps across a range of depolarizing current injections (5 μM: n=11, p < 0.005; 25 μM: n = 6, p < 0.05; Wilcoxon sign tests). The increase in ET excitability was greater at higher concentrations of propofol (5 μM: n=11; 25 μM: n = 6; p <0.05, Wilcoxon Rank test). For IT neurons, no significant changes in excitability were observed at either concentration of propofol (5 μM: n=8, p = 0.875; 25 μM: n = 6, p = 1; Wilcoxon sign tests). In propofol, but not isoflurane, changes to the subthreshold properties of ET neurons were consistent with reduction in HCN-dependent properties (decreased resonance, increased input resistance and membrane hyperpolarization, Sup. Figure 10c-e). Consistent with the direct actions of anesthetics on multiple ion channels, the observed changes in excitability could not be explained by effects on a single ion channel.

### Prefrontal Colgalt2-Expressing Neurons Recapitulate NAP Activity and Function During Anesthesia

Based on the convergence of biophysical and gene expression data, we hypothesized that a molecularly and biophysically defined L5 ET PFC neuron subtype would recapitulate functional, physiological, and anatomical features of NAPs. To test this, we systematically recorded spontaneous neuronal activity in the ACA using calcium imaging during wakefulness and under isoflurane in cre-driver lines that label distinct L5 subtypes, including RBP4-cre (L5 IT/ET), TLX3-cre (L5 IT) and Colgalt2-cre (L5 ET)^125^ (Figure 5a).

Consistent with results in Figure 1, only a small subset of the broad L5, RBP4-expressing neurons exhibited calcium transients under isoflurane. Calcium transients of L5 IT (TLX3) neurons were similarly reduced. In contrast, most (84.2%) L5 ET neurons (Colgalt2) were more active under isoflurane than during wakefulness (Figure 5b). In fact, L5-ET cohort was more enriched for anesthetic-preferring neurons than those originally TRAPed under anesthesia (p<0.0001, Fischer’s Exact test). Of all the major molecularly defined subtypes of excitatory and inhibitory PFC neurons, L5-ET neurons marked in Colgalt2-cre mice were the only population that increased activity under isoflurane anesthesia (Figure 5c).

To independently validate that L5-ET prefrontal neurons are recruited during anesthesia, we quantified c-Fos expression in tdTomato labeled Colgalt2 PFC neurons following exposure to either oxygen or 0.9% isoflurane (Figure 5d). Isoflurane significantly increased the fraction c-Fos^+^ Colgalt2 neurons, particularly within the PL/ILA regions (Figure 5d-e). Thus, calcium imaging and c-Fos expression converge to demonstrate that Colgalt2-expressing neurons are preferentially active during isoflurane anesthesia, distinguishing them from other major neuronal subtypes.

To determine whether prefrontal L5 ET neurons causally modulate anesthetic potency, we expressed chemogenetic actuators (hM3Dq, excitatory; PSAM, inhibitory) in the PFC of Colgalt2-Cre mice. EEG recordings under isoflurane revealed that chemogenetic activation of prefrontal Colgalt2 neurons increased slow-wave activity and reduced higher-frequency power (Figure 5f). Conversely, PSAM-mediated inhibition produced the opposite effect on the EEG. Thus, Colgalt2-expressing prefrontal neurons, similarly to PFC neurons TRAPed under isoflurane, bidirectionally modulate anesthetic EEG signatures.

### Colgalt2 Neurons Are Preferentially Recruited During Recovery NREM and Bidirectionally Regulate NREM Sleep

We next investigated the activity of Colgalt2 neurons during sleep and wake and their causal contribution to NREM sleep. Consistent with the causal role of Colgalt2 neurons in NREM, their chemogenetic activation increased time spent in NREM sleep (Figure 5g, left), while their inhibition produced the opposite effect on NREM (Figure 5g, right).

Activation of isoTRAPed PFC neurons resulted in prolonged bouts of deep NREM sleep (Figure 2). Because deep NREM sleep is typically seen during recovery from sleep deprivation, we investigated the possibility that Colgalt2 PFC neurons are preferentially recruited during recovery sleep. To test this hypothesis, we performed miniscope calcium imaging in the PL/ILA PFC regions of freely behaving Colgalt2-Cre mice implanted with EEG/EMG (as in Figure 2) during 4 hours of mild non-stressful sleep deprivation and the subsequent recovery phase. During the sleep deprivation phase, elicited by introducing novel objects into the home cage, mice spent ∼25% time asleep. During this phase, similarly to isoTRAPed neurons (Figure 2), 53% of Colgalt2 neurons preferentially fired during NREM sleep (Figure 5i, Fischer’s exact test, p=0.82). Remarkably, after sleep deprivation phase ended, and recovery NREM sleep ensued, the fraction of Colgalt2 neurons exhibiting strong NREM preference increased (Figure 5i). Specifically, fraction of neurons exhibiting strong NREM preference (S_NREM_ ≥ 0.75), increased over 3-fold (Figure 5j). Thus, Colgalt2 PFC neurons are preferentially activated during recovery NREM sleep and their activity causally contributes to NREM.

### Connectivity Profile of PFC Colgalt2 Neurons

We next performed whole-brain imaging to characterize projection targets and monosynaptic inputs of prefrontal Colgalt2 neurons. Similar to isoTRAPed neurons, and consistent with their identification as L5-ET neurons, Colgalt2 neurons predominately projected to subcortical structures with the retrosplenial cortex representing the only notable cortical projection target (Figure 6a, Supplemental Table 2). Colgalt2 PFC neurons also densely innervated the striatum and claustrum. Most notably, Colgalt2 neurons exhibited a striking thalamic projection pattern that closely recapitulated that of isoTRAPed neurons. Projections were concentrated within anterior and reticular thalamic nuclei, with minimal labelling in posterior sensory thalamic regions (Figure 6a insets). Additional prominent projections were observed in the ventrolateral periaqueductal gray in the midbrain and the laterodorsal tegmental nucleus, dorsal tegmental nucleus, and pontine central gray areas of the hindbrain. In stark contrast to isoTRAPed neurons, however, Colgalt2 neurons exhibited little to no projection to the preoptic or lateral hypothalamus.

Replication-deficient rabies monosynaptic input tracing revealed that the majority of monosynaptic inputs to prefrontal Colgalt2 neurons arose from neighboring Colgalt2 neurons, indicating extensive local recurrent connectivity (Figure 6b, left), consistent with the strong reciprocal connectivity among prefrontal L5-ET neurons in the PFC^57^. The dominant subcortical input arose from anterior thalamic structures, particularly the anteromedial thalamic nucleus (Figure 6b, right, Supplemental Table 2). These major input pathways closely mirrored the isoTRAP input architecture. Together, these findings demonstrate that prefrontal Colgalt2 neurons recapitulate the core connectivity patterns of NAPs originally defined by the isoTRAP protocol while lacking the prominent hypothalamic connectivity.

### Isoflurane Preferentially Recruits a PFC Colgalt2 Thalamic-Projecting Subpopulation

The weak connectivity between Colgalt2 PFC neurons and the hypothalamus helps narrow down the likely downstream targets through which NAPs may act to promote NREM sleep to the claustrum and thalamus. To begin to dissect these pathways we first sought to determine whether the claustrum and thalamic projecting Colgalt2 PFC neurons are distinct populations. To accomplish this, we selectively labelled claustrum- and thalamus-projecting Colgalt2 neurons using distinct retrograde viruses and examined their spatial distribution and overlap across the prefrontal cortex (Figure 6c). Colgalt2 neurons projecting to the thalamus were found throughout the infralimbic, prelimbic, and anterior cingulate regions of the PFC. In contrast, Colgalt2 neurons projecting to the claustrum were most concentrated in the adjacent motor and cingulate areas. The fraction of Colgalt2 claustrum projecting neurons was progressively diminished in the more ventral prelimbic and infralimbic regions (Figure 6c-d). Only 1.7% (76 of 4471) of Colgalt2 PFC neurons were co-labeled with both retroviruses. Thus, claustrum and thalamus projecting Colgalt2 PFC neurons are largely distinct, anatomically-segregated populations.

Preferential enrichment of c-Fos^+^ Colgalt2 neurons under isoflurane in the PL/ILA regions, rather than the ACA or M1 strongly implies that thalamus projecting neurons, rather than those that project to the claustrum, are preferentially activated under anesthesia (Figure 5e). To test this hypothesis further, we quantified c-Fos expression in thalamus-projecting Colgalt2 neurons following two-hours 0.9% isoflurane exposure (Figure 6e). The expression of c-Fos under isoflurane in the thalamus projecting Colgalt2 neurons in the PL/ILA regions was similar to that observed in the Colgalt2 neuron population as a whole. In contrast, c-Fos expression in claustrum-projecting Colgalt2 neurons following isoflurane exposure was uniformly lower across all PFC regions (Sup. Figure 11). Thus, altogether, calcium imaging in PL/ILA during recovery sleep, c-Fos expression under anesthesia, and the anatomical segregation between the claustrum and the thalamus projecting sub-populations of Colgalt2 neurons, suggest that the thalamus projecting neurons labeled in the Colgalt2 mouse line in the ventral PFC are NAPs.

### PFC Colgalt2 Thalamic-Projecting Neurons Promote NREM Sleep

To determine whether thalamus-projecting Colgalt2 subpopulation contributes to sleep regulation, we selectively expressed hM3Dq in the thalamus-projecting Colgalt2 neurons by bilaterally injecting a cre-dependent retrograde AAV-hM3Dq into the anterior thalamus of Colgalt2-Cre mice. Because Colgalt2 mouse line preferentially labels the PFC, and out targeting of the anterior thalamic nuclei, this results in predominant viral expression in the PFC (Sup. Figure 12). Following CNO administration, chemogenetic activation of thalamus-projecting Colgalt2 neurons significantly increased time spent in NREM sleep while reducing time spent in wakefulness relative to the saline controls (Figure 6 f-g). Time spent in REM was not affected. Further, chemogenetic activation increased NREM bout duration (Figure 6h) and slow-wave activity during NREM sleep (Figure 6i). These results are congruent with those observed in the isoTRAP cohort (Figure 2). Together, these findings demonstrate that selective activation of thalamus-projecting Colgalt2 neurons is sufficient to promote deep consolidated NREM sleep, thus identifying a projection-specific, molecularly defined neuronal population as NAPS – a key node of the shared prefrontal circuit linking sleep and anesthesia.

## Discussion

Here we showed that increased activity of L5-ET corticothalamic neurons in the PFC leads to the suppression of arousal. L5-ET neurons are selectively activated in naturally occurring and pharmacologically induced states of diminished consciousness in contrast to the reduced activity of all other major subtypes of cortical neurons. L5-ET activation is sufficient to promote deep NREM sleep and potentiate effects of anesthesia, while their inhibition partially antagonizes anesthesia and suppresses NREM sleep. Unlike most other types of PFC neurons that communicate extensively with other cortical regions, NAPs are primarily synaptically engaged with anterior thalamic nuclei involved in the regulation of sleep and arousal.

Our findings that PFC plays a key role in potentiating slow cortical rhythms are consistent with the fact that slow waves during NREM sleep^126^ and under anesthesia^9^ are initiated frontally. Our results also align with prior work showing that enhancing excitatory synapses in the PFC promotes NREM sleep^46^, while partially silencing L5 synapses^48^ impairs sleep. Yet, other work proposed that the frontal dominance of slow waves in NREM sleep is coordinated by the thalamus^101^. Specifically it has been suggested, that burst firing in centromedian thalamic nucleus precedes and entrains cingulate UP states and that the anterodorsal thalamic nucleus play a key role in synchronization of UP states across the neocortex^101^. Consistent with these prior findings, we show that NAPs bidirectionally interact with these thalamic nuclei. While we show that *in vivo* NAP activation coincides with the transition from wakefulness to NREM sleep, calcium imaging and EEG-based sleep state classification lack the millisecond temporal resolution needed to determine the precise temporal sequence of cortical and thalamic burst firing and slow waves. The fact that NAP activation is sufficient to promote NREM characterized by large amplitude slow oscillations and bidirectionally control their amplitude under anesthesia, however, strongly suggests that the PFC plays a key role in producing slow oscillations that define states of reduced arousal.

While we demonstrate that activation of corticothalamic L5-ET neurons is sufficient to drive deep NREM sleep, we do not at present know whether the interactions with the thalamus *per se* are critical for this effect. Our observations and previously published data^127^ both suggest that the axons of L5-ET corticothalamic neurons collateralize, with one branch innervating the thalamus while the other travels through the internal capsule to innervate hindbrain structures also thought to be involved in the control of arousal. The degree to which hindbrain and thalamic projections of these neurons contribute to the generation of slow oscillations and the circuit mechanisms responsible for the generation of slow waves awaits future investigation.

Although NAPs show strong bidirectional interactions with the thalamus, our limited transcriptomic analyses revealed no L6 corticothalamic (L6 CT) neurons, consistent with predominant localization of NAPs in L5 rather than L6. Interestingly, recent work has begun to elucidate unique features of PFC-thalamic loops that distinguish them from other cortical regions. In the PFC, L5 ET neurons directly excite the reticular nucleus (RT) of the thalamus^89^, a connection observed predominantly for L6 CT neurons elsewhere in the cortex^129^. Consistent with this work, we observe strong NAP projections to the reticular nucleus (Figure 6). Burst firing of RT neurons is correlated with UP states observed in the LFP under anesthesia and L5 projections to the RT play a key role in this corticothalamic synchronization^128^. Large calcium transients observed in NAPs under anesthesia and during NREM sleep may therefore drive RT bursts linked to the generation of slow waves^130,131^. Previous findings suggest that L5 cortical neurons synapse on a specific subtype of RT neurons that selectively project to “higher-order” thalamic nuclei^127^ and that molecularly distinct RT neuron subtypes have different effects on NREM sleep^132^. Determining whether NAPs engage a specific RT neuron subtype and the investigation of the physiology of these synapses may reveal functional implications of the unique PFC-thalamic loop in the control of sleep and arousal.

Recent work revealed that inhibition of RT neurons by lateral hypothalamic (LH) GABAergic neurons triggers awakening from sleep^133^. Thus, NAPs and wake promoting hypothalamic neurons converge on the RT and exert opposing effects on its activity, suggesting a key role of this circuit in regulation of levels of consciousness, a hypothesis supported by computational models^134^. NREM-promoting PFC SST neurons inhibit LH GABAergic neurons^41^, and may therefore release RT neurons from inhibition. Together these results outline the specific cell types that comprise a unique PFC-thalamic loop integrated with ascending modulatory hypothalamic nuclei to control the level of consciousness.

A subset of isoTRAPed neurons and neurons labeled in the Colgalt2-cre mouse line have a strong projection to the claustrum – a structure involved in coordinating synchronous slow wave activity across evolution from reptiles to humans^89–91,94^. Wide projections of the claustrum throughout the neocortex have been used to suggest that claustrum plays a key role in the regulation of conciousness^135^, a controversial claim^136^. As we show, NAP neurons projecting to claustrum and thalamus are distinct and anatomically segregated into different subregions of the PFC. While our results point to the importance of the thalamus projecting PFC L5 ET neurons, the contribution of the subpopulation projecting to the claustrum remains unclear at present.

While activity TRAP under isoflurane identified PFC neurons with strong projections to the hypothalamus, this projection was weak in the Colgalt2-cre mouse line. Previous work^62^ showed that glutamatergic neurons that project from the PL/ILA to the hypothalamus control REM sleep. By anatomical definition these hypothalamus projecting neurons are ET; however the fact that they are not labeled by the Colgalt2-cre mouse line strongly implies that they are molecularly distinct from the thalamic projecting L5 ET neurons that drive NREM sleep. The nature of this molecular distinction, its relationship to the biophysical properties, and the connectivity between the PFC glutamatergic neurons that control different phases of sleep is a fascinating direction for future investigation.

Previous work on subcortical circuits mediating effects of anesthesia shows that anesthetics activate sleep-promoting neurons^11,14,15,17,18,74,96,137–140^. Our results here show that the convergence between sleep and anesthesia circuits extends to the neocortex. Most known actions of anesthetics on cortical neurons^19,20^ and synapses^141,142^ suppress neuronal activity. We identify PFC L5 ET neurons as the only major cortical neuronal subtype that increases activity under anesthesia *in vivo*, pointing to their unique properties. Remarkably, we find that both isoflurane and propofol increased the excitability of synaptically isolated L5-ET neurons in PFC slices. The effects of propofol and isoflurane were different on the IT neuron population. Propofol left L5-IT excitability nearly unchanged whereas isoflurane profoundly suppressed the ability of IT neurons to fire action potentials in response to current injections. Consistent with previous demonstrations that propofol^119–121^ inhibits HCN1 channels, we found that propofol-induced changes in L5-ET were consistent with a reduction in HCN-dependent properties: reduced somatic resonance frequency, resting membrane potential hyperpolarization, and increased input resistance (Sup. Figure 10). In contrast, we did not observe a shift in these properties in isoflurane, which similarly has been shown to inhibit HCN1 channels^117,118^. Because HCN1 channels are enriched in the apical dendrites of L5-ET neurons, we cannot exclude the possibility that our somatic recordings failed to capture isoflurane-induced changes to these ion channels in more remote dendritic compartments. It is important to note, however, that while direct cell-type specific effects of mechanistically distinct anesthetics on L5-ET excitability may contribute to their unique activation under anesthesia *in vivo*, the relationship between drug effects in synaptically isolated slices and *in vivo* activity is complex.

Several known molecular anesthetic targets are differentially expressed in L5-ET PFC neurons and this differential expression may account of the cell-type specificity of anesthetic effects on NAP excitability. Undoubtedly, this list of potential targets is incomplete. Our results on both the similarities and differences between the effects of propofol and isoflurane on the excitability of distinct PFC glutamatergic neuronal subtypes are consistent with the view that that anesthetics are likely to have multiple ion channel targets and no single shared receptor is likely to account for effects of all anesthetics. While we show that both isoflurane and propofol increase the excitability of L5-ET neurons, it is not clear whether these mechanistically distinct drugs converge on the same molecular targets to produce this effect. Excitability is a coarse measure that depends on the interactions between many ion channels as well as other variables such as cell morphology. Our preliminary analysis of the finer-grained features of anesthetic effects on ET and IT physiology revealed significant heterogeneity in both neuron populations even in highly reduced preparation (Sup. Figure 10). This suggests that even within a specific subtype of PFC glutamatergic neurons, anesthetics may act upon distinct molecular targets. This is not entirely surprising as both L5-ET and, especially, L5-IT neurons are broad subtypes that consist of several interrelated neuronal populations.

Future work may identify NAPs as a specific molecular subset of L5-ET neurons. Together with the availability of extensive cell-type specific gene expression datasets^55,56^, this may have clinical implications for developing novel sedatives or for alleviating PFC dysfunction after anesthesia^143,144^. It may appear surprising that low affinity, molecularly promiscuous general anesthetics may exert cell type specific effects. However, modern single-nucleus gene expression profiling revealed that distinct neuronal types are defined by a unique combination of expressed genes rather than a single marker^55^. Therefore, to pharmacologically affect a specific cell type, a drug must be able to differentially affect combinations of targets rather than exhibit high selectivity for a single receptor. A proof of principle for this idea has been obtained by specifically designing a drug cocktail that selectively affects a specific subset of locus coeruleus neurons^145^.

Interestingly, previous work in humans and in animal models singled out anesthetic induced suppression of the PFC as critical for disruption of consciousness. Consistent with this view, loss of consciousness induced with variety of anesthetics in humans is associated with changes in frequency and coherence of frontal EEG signals^146,147^ and disruption of fronto-parietal functional connectivity^148–150^. Cholinergic activation of the PFC can precipitate emergence from anesthesia^151^ while nonspecific inhibition of the PFC delays anesthetic emergence^152^ and decreases wakefulness^153^. Further, cell-type agnostic chemogenetic inhibition of glutamatergic neurons in the PFC potentiated anesthetic effects while nonspecific excitation reduced anesthetic potency^21^. These findings have been interpreted as lending support to the Global Neuronal Workspace theory of consciousness^154^ and implicate the PFC as a key arousal node^155^. Interestingly, together with previous findings,^41^ this work strongly implies that the PFC contains both arousal promoting and arousal suppressing microcircuits composed of distinct neuronal subtypes, consistent with the fact that PFC is a highly heterogeneous structure^156^. Understanding the interplay between these microcircuits is essential for deeper insights into mechanisms that control the level of consciousness.

It has been proposed^24^ that the primary effect of anesthetics at the cellular level is the decoupling of dendritic from somatic activity in L5 neurons. Consistent with this hypothesis, both IT and ET L5 neurons exhibit synchronous somatic calcium fluctuations dissociated from dendritic signals^25^. Rather than focusing on relatively small amplitude fast calcium dynamics which may reflect UP and DOWN states^25^, here we focused on large calcium transients thought to be related to nonlinear dendritic integration events^157^ associated with burst firing^158^. Our findings reveal that L5 ET PFC neurons are the only major cortical neuronal subtype to preferentially exhibit large calcium bursts under anesthesia, as corroborated by c-Fos expression. We did not directly address the relationship between the dendritic and somatic signals in NAPs. Hence, it remains unclear whether uncoupling of dendritic and somatic signals also occurs in the PFC L5 ET neurons. However it is notable that L5 ET and L5 IT neurons exhibit distinct dendritic physiology that could contribute to the cell-type differences that we observe^109,110^.

All animal behaviors are hierarchically organized into orderly sequence of phases^159,160^. Prior to the onset of sleep, animals exhibit a set of preparatory behaviors^41,161^. Upon their completion, sleep itself proceeds in orderly stages from progressively deepening NREM sleep to REM^100^. It now seems clear that the PFC contains distinct neuronal populations that control all sleep stages: SST neurons that initiate preparatory behaviors^41^, L5 ET neurons that promote NREM sleep, and the hypothalamic projecting PFC glutamatergic neurons that initiate REM^62^. Understanding the dynamics of the PFC microcircuits that interlink these neuronal populations will reveal the mechanics of the orderly progression through different behavioral states that comprise sleep.

There is, however, a deeper level at which sleep needs to be understood. The decision to sleep, the simplest decision made by animals across evolution, must be integrated within the broader behavioral hierarchy. Even relatively “simple” animals must decide whether it is safe to sleep, and this requires executive function that weighs homeostatic and circadian sleep pressures against other behavioral needs. It is therefore unsurprising that the PFC emerges as a central cortical hub that controls sleep. In most studies, including this one, sleep is studied in a highly contrived lab environment where the decision of whether to sleep or to stay awake lacks behavioral consequences. Understanding how the PFC implements the decision to sleep at the expense of all other behaviors will require interrogation of the PFC circuits in more naturalistic settings.

## Materials and Methods

All experiments were approved by the Institutional Animal Care and Use Committee at the University of Pennsylvania (IACUC protocol # 803844, 804902, and 807237). All ex-vivo slice experiments were performed in accordance with Institutional Animal Care and Use Committee at the University of Washington (IACUC protocol # 4586-01). Male and female mice aged 12-46 weeks were equally distributed across all experiments and cre-lines except for the Colgalt2-cre mouse line. Only male Colgalt2-cre mice were used as Cre-recombinase expression in this line is driven by a Y chromosome-linked transgene. This broad age range reflects the extended preparation time required for isoTRAP labeling, GRIN lens implantation and window clearing, and, for some cohorts, interinstitutional transfer and mandatory quarantine before experimentation. All experiments were conducted in accordance with the National Institutes of Health guidelines. Mice were housed on a 12h:12h reverse light dark cycle (ZT0/lights on at 9:00pm) with *ad libitum* access to food and water. For experiments examining effects of chemogenetic inhibition on sleep/wake, mice were instead housed on a standard 12h:12h light dark cycle (ZT0/lights on at 9:00am). Table 1 lists all the mouse strains used in this work. Table 2 lists all viruses used in this work.

**Table 1:** Mouse strains used in this work.

| Mouse Line | Source | Description/Purpose |
| --- | --- | --- |
| C57BL/6J | JAX: 000664 | Wild type |
| TRAP2-cre (*Fos*<sup>tm2.1(icre/ERT2)Luo/J</sup>) | JAX: 030323 | Activity-tagging neurons |
| PV-cre (B6;129P2-*Pvalb*<sup>tm1(cre)Arbr/J</sup>) | JAX: 008069 | Interneuron line |
| SST-cre (*Sst*<sup>tm2.1(cre)Zjh/J</sup>) | JAX: 013044 | Interneuron line |
| VIP-cre (*Vip*<sup>tm1(cre)Zjh/J</sup>) | JAX: 010908 | Interneuron line |
| Ai6 (B6.Cg-Gt(*ROSA*)26*Sor*<sup>tm6(CAG-ZsGreen1)Hze/J</sup>) | JAX: 007906 | Fluorescent reporter |
| RBP4-cre (Tg(*Rbp4*-cre)KL100Gsat/Mmcd) | MMRC: 031125 | Layer 5 line |
| TLX3-cre (B6.FVB(Cg)-Tg(Tlx3-cre)PL56Gsat/Mmucd) | MMRC: 041158 | Layer 5 IT-neuron line |
| Colgalt2-cre (Tg(*Glt25d2*-cre)NF107Gsat/Mmucd) | MMRC: 036504 | Layer 5 ET-neuron line |
| Thy1h-eyfp (B6.Cg-Tg(Thy1-EYFP)-HJrs/J) | JAX:003782 | Layer 5 ET-neuron line |
| Etv1-egfp Tg(Etv1-EGFP)-BZ192Gsat/Mmucd) | MMRC: 011152-UCD | Layer 5 IT-neuron line |

**Table 2:** Viruses used in this work.

| Virus Name | Source | Concentration (GC/ml) | Description/Purpose |
| --- | --- | --- | --- |
| *Chemogenetics* |  |  |  |
| AAV8-hSyn-DIO-hM3Dq-mCherry | Addgene: 44361 | 2.1 x 10<sup>13</sup> | Chemogenetic excitation |
| AAV9-Syn-flex-PSAM4-GlyR-IRES-EGFP | Addgene: 119741 | 2.1 x 10<sup>13</sup> | Chemogenetic inhibition |
| AAV8-hSyn-DIO-mCherry | Addgene: 50459 | 2.2 x 10<sup>13</sup> | Chemogenetic control |
| AAVrg-hSyn-DIO-hM3Dq-mCherry | Addgene: 44361 | 1.9 x 10<sup>13</sup> | Chemogenetic excitation |
| *Anatomic Tracing* |  |  |  |
| AAV9-CAG-FLEX-tdTomato | Addgene: 28306 | 3.1 x 10<sup>13</sup> | Anterograde tracing |
| AAV9.CMV.FLEX.TVA.mCherry.2A.oG | Salk: 102985 | $1.37 \times 10^{13}$ | Monosynaptic input tracing: helper adenovirus |
| EnvA G-deleted Rabies-EGFP | Salk: 32635 | $6.1 \times 10^8$ | Monosynaptic input tracing: modified rabies virus |
| AAVrg-pCAG-FLEX-EGFP-WPRE | Addgene: 51502 | $2.1 \times 10^{13}$ | Retrograde tracing viral vector |
| pAAVrg-CAG-FLEX-tdTomato | Addgene: 28306 | $2.1 \times 10^{13}$ | Retrograde tracing viral vector |
| *Calcium Imaging* |  |  |  |
| pGP-AAV9-Syn-Flex-JGCaMP8s | Addgene: 162377 | $2.3 \times 10^{13}$ | 1P miniscope imaging |
| AAV9-CAG-Flex-GCaMP6f-WPRE-SV40 | Addgene: 100835 | $2.1 \times 10^{13}$ | 2P imaging, specific neuronal populations |
| AAV1-hSyn-GCaMP6f-WPRE-SV40 | Addgene: 100837 | $2.2 \times 10^{13}$ | 2P imaging: pan-neuronal |
| pENN-AAV9-CamKII-Cre-WPRE-SV40 | Addgene: 105551 | $1.1 \times 10^{13}$ | 2P imaging: pyramidal neurons |

### General surgical procedures

All surgeries were performed following IACUC guidelines for rodent survival surgery. Mice were induced with 2.50% isoflurane in 100% oxygen and maintained at 1.50% isoflurane for the duration of the surgery. After a surgical plane of anesthesia was reached, as measured by the lack of a toe pinch response, mice were placed in a stereotactic frame (Kopf Model 940). Core body temperature was regulated with a closed-loop heating pad (CWE Inc, TC1000), and ophthalmic ointment was applied to protect the eyes. All surgeries were performed with aseptic technique and sterile instruments. All animals were administered analgesics and antibiotics along with fluids intraperitoneally prior to surgical incision.

### Stereotaxic Viral Delivery

The skull was first leveled to accommodate stereotactic coordinates.^162^ Burr holes were created with a stereotaxic microdrill (Stoelting, 51449) using 0.5 mm diameter drill bit (Fine Science Tools, 190070-05). Viruses were delivered using a 32 gauge, 10μL syringe (Hamilton, 80008) mounted on a stereotactic microinjection syringe pump (Stoelting Co, 53311). PFC virus injection coordinates were: AP 1.70mm, ML 0.30mm, DV -3.00mm relative to bregma. Thalamic virus injection coordinates were: AP -1.00mm, ML 0.75mm, DV -3.50mm relative to bregma. Claustrum virus injection coordinates were: AP 0.50mm, ML 3.10mm, DV -4.15mm relative to bregma. Prior to and following viral injection, the syringe was held in position for 10 minutes to lessen tissue trauma and to avoid virus reflux up the syringe tract.^163^ Following injection, burr holes were filled with bone wax (Fine Science Tools, 19009-00) and the incision was sutured closed. Mice were allowed a minimum of 1 week recovery time prior to any other experimentation.

*Chemogenetic actuator viruses:* Bilateral (300nL per side) injections of either AAV8-hSyn-DIO-hM3Dq-mCherry, AAV9-Syn-flex-PSAM4-GlyR-IRES-EGFP or AAV8-hSyn-DIO-mCherry (excitation, inhibition, and control respectively) were targeted at PFC coordinates from above. For thalamus-targeted AAVrg chemogenetic experiments, bilateral (150nL per side) injections of AAVrg-hSyn-DIO-hM3Dq-mChery were delivered. *Anterograde Projection viruses:* Unilateral 300 nL injection of AAV9-CAG-FLEX-tdTomato was performed. *Monosynaptic input viruses:* 300 nL of AAV9.CMV.FLEX.TVA.mCherry.2A.oG helper virus was injected into the PFC unilaterally. After 3 weeks of viral expression, 1 μL EnvA G-deleted Rabies-EGFP was injected at the same stereotaxic coordinates. *Retrograde Tracing Viruses:* Unilateral 150 nL injection of AAVrg-pCAG-FLEX-EGFP-WPRE or AAVrg-FLEX-tdTomato were injected into the thalamus or claustrum, respectively. *Calcium imaging in the PFC:* For GRIN mediated imaging of the PFC, 300 nL of pGP-AAV9-Syn-Flex-JGCaMP8s was injected into the PFC unilaterally.

### EEG/EMG Electrode Implantation

Electrodes were constructed and implanted as previously described.^164^ Briefly, ten epidural EEG leads (Digikey, 853-93-012-10-001000-ND) were implanted 0.65mm lateral to bregma in both hemispheres, ranging from 2.30 mm anterior to bregma to -2.90 mm posterior to bregma. 2 stainless steel EMG wires were soldered onto two separate headpiece connecter leads and were implanted in the dorsal neck muscles (A-M Systems, 793500). Each insulated wire coating was stripped prior to soldering, followed by a reinsulating nail polish coating. Headpieces were secured using dental cement (A-M Systems, 525000 and 526000) and two anchor screws (McMaster-Carr, 91800A050), positioned at 2.50 mm lateral and 2.00 mm posterior to Bregma. Following surgery, mice were singly housed and were allowed to recover from the implant for a minimum of one week prior to investigation. For EEG recordings in TRAP2-cre mice undergoing chemogenetic modulation, electrode implantation was performed at least one week following the isoTRAP protocol (see below).

### Combined EEG and PFC Calcium Imaging in Awake Behaving Animals

For simultaneous EEG and one-photon miniscope recordings, we constructed a custom 2×2 headpiece connector by trimming the 2x6 EEG connector to accommodate the miniscope. Modified EEG electrodes were made using stainless steel screws (McMaster-Carr, 91800A052) connected to headpiece via silver wire wrap (A-M Systems, 787000) and solder. Two stainless steel EMG wires were soldered to separate leads of the headpiece connector as above.

Standard V4 variant 2 baseplate (Open-Ephys, OEPS-7416) was modified to enable real-time visualization of GCaMP expressing neurons within the field-of-view (FOV) during GRIN lens implantation. Specifically, we removed the portion of the baseplate cap that normally fits inside of the metal baseplate. Then we drilled a 3.0 mm diameter recessed hole in the center of the modified cap to a depth of 1.2mm, the approximate working distance of the miniscope. A second, precision reamed through-hole with a 0.51mm diameter was drilled, centered about the recessed hole. To ensure precise centering of the GRIN lens within the baseplate, a second baseplate cap with a similar centered, 0.51 mm precision reamed through hole served as a template alignment tool. This template cap was secured to a metal baseplate in its standard orientation and was used to align the modified baseplate cap with a precision 0.51mm pin gauge (McMaster-Carr, 2326A312) inserted through both template cap and modified cap. The recessed circular portion of the modified cap was confirmed to be oriented upwards and then secured to the underside of the metal baseplate with cyanoacrylate adhesive (McMaster-Carr, 7520A12). With the alignment cap and pin gauge removed, a 0.5mm GRIN lens (Inscopix, 1050-004600, 8.4mm length) was press-fit, flush with the top of the recessed area and secured with cyanoacrylate adhesive.

GRIN-lens and EEG/EMG electrode implantation surgery was performed at least one week following the completion of the isoTRAP protocol. Prior to surgical incision, mice were administered 0.2 mg/kg dexamethasone IP. Burr holes were drilled centered at AP -2.0 mm and ± 2.0 mm for EEG screws, AP +3.0 mm, ML 1.7 mm for an additional anchor screw, and AP +2.0 mm, ML −0.3 to −0.4 mm for GRIN lens. The mediolateral coordinate for the GRIN lens was adjusted for each mouse to avoid injury to the sagittal sinus and reduce intraoperative bleeding risk. EEG and anchor screws were secured, followed by EMG wire insertion in the dorsal neck muscles. Next followed stereotactic aspiration of brain tissue above infralimbic area to a depth of ≈1.5mm using a 25G blunt needle, as previously described.^165^ The cavity was perfused with sterile saline and irrigated thoroughly until the effluent became clear. For modified baseplate/GRIN implantation, we utilized the stereotaxic miniscope holder designed by the Aharoni Lab (https://github.com/Aharoni-Lab/Miniscope-v4/tree/master/Miniscope-v4-Holder). The miniscope was mounted onto the modified baseplate with the GRIN lens, allowing real-time visualization of the FOV during implantation. The lens was slowly lowered to a depth of approximately 3.0 mm while monitoring the FOV. The final position was determined when individual cells around this depth (DV -2.8mm to -3.0 mm) became visible. GRIN lens was then secured by layering cyanoacrylate gel adhesive rapidly cured with cyanoacrylate accelerator, cyanoacrylate adhesive, and dental cement, with adequate drying times between. Following surgery, mice were singly housed and were allowed to recover from the implant for a minimum of three weeks to allow for window clearance prior to additional studies. Dexamethasone, cefazolin, and meloxicam were administered for the first 3 post-operative days.

### *In vivo* two photon imaging

#### Calcium indicator Expression

Two-photon in-vivo calcium imaging studies were performed using AAV9-CAG-Flex-GCaMP6f-WPRE-SV40 in specific neuronal populations labeled in transgenic mouse lines (Table 1). To attain pan-neuronal imaging, pyramidal neurons were labeled using pENN-AAV9-CamKII-GCaMP6f-WPRE-SV40 and AAV9-CAG-Flex-GCaMP6f-WPRE-SV40, or with AAV1-Syn-GCaMP6f-WPRE-SV40. GCaMP6f transfection was achieved using intracranial AAV injections of neonatal mice as previously described.^166,167^

#### Cranial Window Implantation

1.5-3 months following viral transfection head-mount and acute cranial windows implant surgery was performed as previously described.^168^ Briefly, periosteal tissue across the skull was removed without damaging either temporal or occipital muscles. A mounting bracket consisting of two parallel metal bars was attached to the animal’s skull to aid in head restraint and reduce motion artifacts during imaging. Mice were screened for GCaMP expression, after which a small craniotomy was made over either the primary somatosensory cortex (S1; 0.5 mm posterior and 1.5 mm lateral to bregma) or the anterior cingulate area (ACA; 0.5–1.0 mm anterior to bregma and 0.3–0.5 mm lateral to the midline) based on stereotaxic coordinates. A round glass coverslip (Thomas Scientific, 1217N66) approximately matching the size of the craniotomy was secured in place using cyanoacrylate adhesive and adhesive resin cement (Parkell, S380). Mice were habituated to head restraint twice daily for 30 minutes on postoperative day 1 and 2 in a custom-built body support to reduce stress.^168^ No obvious distress was observed during the habituation periods across all experimental setups.

#### 2-Photon Imaging

On the day of imaging, awake mice were positioned in the custom head holder device under the two-photon microscope. In vivo two-photon imaging was performed with an Olympus DIY RS two-photon system (tuned to 910–920 nm) equipped with a Coherent Discovery NX laser. Pyramidal neurons in ACC region were recorded for 2-min sessions under awake conditions and once again after 0.6% isoflurane while maintained normothermic with a heating pad. Isoflurane imaging began 30 minutes after anesthetic induction to ensure steady-state conditions. Motion-related artifacts were typically <2 µm. Vertical movements were rare and minimized using two metal bars affixed to the skull (described above) in combination with a custom-built body support.^168^ All experiments were performed using a ×20 Olympus objective (XLUMPLFLN; NA = 1.00, 2.0 mm working distance) immersed in aCSF, with ×2 digital zoom. Images were acquired at a frame rate of 2–4 Hz (2-μs pixel dwell time). Image acquisition was performed using Olympus Fluoview software and analyzed post hoc using ImageJ software version 2.1.0.

### Isoflurane TRAP (isoTRAP) Exposure Protocol to Label Neurons Active Under Anesthesia

This protocol was performed 1-2 weeks following stereotaxic injection of cre-dependent virus into the PFC (see above) to permanently label neurons active under isoflurane anesthesia with a construct of interest. TRAP2-cre mice were habituated to gastight, temperature-controlled, 200mL cylindrical chambers with 100% oxygen flowing at 200 mL/min for two hours per day for three days prior to experimentation, as previously described.^169^ After 1-hour pretreatment with 0.6% isoflurane, mice were injected with 4-hydroxytamoxifen (4-OHT, 40mg/kg IP) to drive nuclear expression of Cre and enable cre-mediated recombination in isoflurane active neurons. Time exposed to open air during IP injections was less than 20 seconds per mouse.^74^ Mice remained exposed to 0.6% isoflurane for another 6 hours following tamoxifen injection. Anesthetic concentrations were confirmed using a Riken FI-21 refractometer (AM Bickford). Mice were allowed a minimum of 1 week recovery to allow for viral expression prior to any further experimentation. A stock concentration of 5mg/mL of 4-hydroxytamoxifen (Hello Bio, HB6040) solution was prepared by dissolving the 10mg of drug in 0.5 mL 100% ethanol, heating the suspension at 60 C for 10 minutes, then adding 0.5 mL kolliphor oil (Sigma-Adrich, C5135), and lastly adding 1.0 mL 0.1X PBS with 10 minute vortexing period between each step.^170^

### Chemogenetic Agonist Preparation and Dosing

Clozapine-N-oxide, CNO (Sigma-Aldrich, C0832), was dissolved in sterile saline to a final concentration of 0.3 mg/mL. CNO dosing for all experiments was 3.0 mg/kg IP.^171^ uPSEM 817 tartrate (Tocris, 6866) was dissolved in sterile saline to a final concentration of 0.1 mg/mL. uPSEM dosing for all experiments was 1.0 mg/kg.^172^ All drug preparations were prepared fresh for each experiment and all drugs were administered intraperitoneally (IP).

### Repeated Righting Reflex Assessment

Anesthetic was delivered and monitored using the same procedure as above. Righting reflex (RR) assessments were performed under steady state isoflurane 0.6 % conditions^82,173^ as described previously. RR was then assessed every 3 minutes. Successful righting probability was estimated as fraction of all trials on which righting was achieved in sliding windows of 10 trials (window step 1 trial). 95% confidence bounds were estimated using the binomial distribution.

For each chemogenetic manipulation experiment, mice first underwent a control injection of saline. Following the saline injection, with isoflurane maintained at 0.60%, the righting reflex was assessed every 3 minutes for 1 hour.^74,82,173,174^ Mice expressing hM3Dq DREADDs received an IP CNO injection, while mice expressing PSAM-GlyR received an IP uPSEM injection, at the conclusion of the final saline righting reflex assessment. After a 15-minute re-equilibration period, mice were subjected to an additional 3 hours of righting reflex assessments at 3-minute intervals.

### EEG/EMG and Miniscope Recordings

#### EEG recording apparatus

A 32-channel headstage (Intan Technologies, C3324) linked to an acquisition system (Intan Technologies, C3004, controlled with RHX software) was used for all neurophysiological recordings. EEG, EMG and 3-axis headstage accelerometer signals were acquired continuously at 1KHz per channel. The headstage interfaced with the EEG and EMG electrodes via a custom-designed PCB adaptor. All filtering and processing of the EEG signals was performed *post-hoc*. A slip-ring commutator (Digikey, 152-1175-ND) attached to a spring counter-balanced lever arm (Instech, CMMI/65) was used to connect the EEG headstage to the acquisition system using SPI cable adapter boards (Intan Technologies, C3430).

#### GRIN-mediated in vivo Ca imaging apparatus

Miniscope (Open Ephys, v4.4, OEPS-7407) was used to acquire calcium signals sampled at 20 Hz. 5 minutes of continuous recordings were interspersed with 5 minutes of rest to avoid excessive photobleaching and heat damage. LED power was maintained at a constant level. Minimal sufficient LED power (1<mW) was used in all recordings. For combined EEG/Miniscope recordings the signals were connected to a motorized commutator (LabMaker, Open-MAC Commutator) enabling unrestricted movement during recordings.^175^ Calcium signals were recorded with a Miniscope-DAQ (Open Ephys, v3.3, OEPS-9050) controlled with Miniscope DAQ QT Software v1.11 (Aharoni Lab, https://github.com/Aharoni-Lab/Miniscope-DAQ-QT-Software/releases).

#### Video tracking of behavior

An infrared LED (Adafruit, 387) was soldered to the EEG headstage and tracked using a USB infrared camera (SVPRO OV2710), modified to operate exclusively under infrared conditions, allowing continuous monitoring of mouse movements. Video camera frames were acquired at 5 Hz. Real-time LED centroid tracking was used to estimate ambulatory activity as described previously.^176^

#### Signal Synchronization

Miniscope images, video camera frames, anesthetic gas concentration, and light: dark cycle, were synchronized to the electrophysiological signals via an I/O expansion module (Intan Technologies, E6500). Synchronization was achieved using TTL pulses delivered by an Arduino Uno (Mouser Electronics, A000066) and managed via custom built MATLAB (MathWorks, 2024a) interface.

#### Anesthesia EEG Experiments

EEG recordings under anesthesia were performed in a gas-tight, temperature controlled, cylindrical recording chamber as described previously in tether-habituated mice.^177^ Briefly, fresh gas was delivered at 8 L/min (approximately one full volume exchange per minute). Body temperature was maintained at 37°C via partial submersion of the chamber in a circulating 37°C water bath. Anesthetic gas concentrations were sampled using a Poet IQ 602-6A Gas Monitor (Criticare Systems). For chemogenetic modulation experiments, mice expressing either excitatory (hM3Dq) or inhibitory (PSAM) receptors in the PFC were exposed to 0.4% isoflurane for 1 hour. Then, hM3Dq-expressing and PSAM-expressing mice received CNO or uPSEM respectively (CNO and uPSEM preparation and dosing in: Chemogenetic Agonist Preparation and Dosing section). EEG recordings continued for another 3 hours following chemogenetic agonist delivery.

#### Miniscope Anesthesia Experiments

Mice implanted with a GRIN lens were exposed to either 0.6% isoflurane, 1.0% isoflurane, 50 mg/kg propofol, 100 mg/kg propofol, 400 µg/kg dexmedetomidine, or 50 mg/kg ketamine (all non-volatile anesthetics were delivered IP), and calcium activity was recorded for at least 60 min beginning at anesthetic administration. In the 1.0% isoflurane exposure experiments, an extended (∼1.5 hours) baseline period was recorded to quantify calcium transient activity during wakefulness and NREM sleep while maintaining the same neurons in the field of view during subsequent isoflurane exposure, enabling direct assessment of whether individual neurons exhibited similar activity state preferences during NREM sleep and isoflurane relative to wakefulness.

#### Sleep/Wake Recordings

All sleep/wake EEG recordings were conducted in a dedicated room in custom-built sound-attenuated recording chambers that fit each animal’s home cage. A 12:12, light-dark cycle synchronized to that used in the animal facility was maintained in each recording chamber. Animals were given access to food and water ad-libitum. Prior to each recordings mice were habituated to the apparatus and tethering for at least 24 hours. This habituation period was followed by 1-24 hrs of baseline recordings. Chemogenetic activation of neurons of interest was performed at ZT=18 (middle of the dark phase) while chemogenetic inhibition was performed at ZT=6 (middle of light phase) and followed by at least 3 hours of continuous recordings. For sleep/wake EEG experiments with simultaneous miniscope recordings, mice expressing GCaMP were recorded in the same apparatus for a period between 4 and 24 hours.

#### Miniscope Sleep Deprivation and Recovery Sleep Recordings

Simultaneous miniscope calcium imaging recordings and EEG were recorded during a 4-hour novel object-mediated sleep deprivation period followed by a recovery sleep period of at least 3 hours. During sleep deprivation, novel objects were introduced into the recording cage every 10-20 min to maintain wakefulness. To promote recovery sleep, the object was removed from the recording cage, and mice were left undisturbed for the remainder of the recording session.

### Analysis of Electrophysiological Signals

All analysis of electrophysiological signals was performed *post-hoc* using custom MATLAB code. EEG signals were decimated to 125 Hz, high pass filtered (4^th^ order Butterworth filter, cutoff frequency of 0.5Hz) and mean re-referenced. EMG signals were high pass filtered (4^th^ order Butterworth filter, cutoff frequency of 300Hz) and differentially referenced. All filtering was performed in both directions to minimize phase delays (MATLAB *filtfilt* function). Multitaper power spectral estimates (35 tapers, 5 second window, 2.5 second step) were computed using Chronux.^178^ EMG and accelerometer RMS was computed in the same temporal windows. A single EEG channel was selected for subsequent sleep wake analysis based on visual inspection for absence of large-scale artifacts and with impedance ≤ 100 kΩ.

To assess changes in spectral power in anesthesia EEG experiments before and after chemogenetic modulation, we computed the ratio of spectral power at each frequency. 30 minutes of artifact free EEG was used to estimate power spectrum for each mouse for the 30 minutes prior to chemogenetic agonist injection or 60-90 minutes following injection. 95% confidence intervals for resulting power ratios were estimated using jackknife resampling across mice.

To assess changes in sleep/wake, we developed a feedforward, fully connected neural network classification model, using MATLAB’s *fitcnet* function. Input features were extracted from 5-second windows of EEG, EMG, and accelerometer data. Dimensionality of the EEG spectrogram was reduced using non-negative matrix factorization ^179^, to six features (91.4% of spectral variance captured). These were augmented with EMG RMS and accelerometer RMS to form an eight-dimensional input vector. The network was trained in a supervised manner using manually scored labels (Wake, NREM, REM) from a subset of data recorded from 16 mice. Training set contained approximately equal proportions of all states. Network classification was validated using a leave-one-mouse-out cross-validation strategy. The neural network classifier achieved an overall training accuracy of 95% and overall testing accuracy (on left out data) of 93% across all folds. (See Sup. Figure 4)

Total time spent in each state, NREM bout duration, and slow wave activity (SWA) were calculated and compared for each state and compared between groups. SWA was defined as the faction of power within the 0.5– 4 Hz frequency range. NREM bout duration and SWA were normalized for each animal’s baseline period (1 hour prior to chemogenetic modulation). 95% confidence intervals for resulting comparisons were estimated using jackknife resampling across mice. 1-way or 2-way ANOVAs were used to compare Fraction of Time Spent in each vigilance state, NREM bout durations, SWA, and locomotor activity across 1 hour epochs before and after injection. Lastly, transition probability matrices were estimated during the CNO period (0.5–2.5 hours post-injection) and compared to those estimated during a zeitgeber-matched baseline period 24 hours earlier. 7 hM3Dq and 8 mCherry mice were used for this analysis; excluded mice did not have baseline ZT matched recordings. These transitions matrices were compared with 2-way ANOVAs as well. To eliminate transient fluctuations, classified network states were required to persist for at least five consecutive windows.^180^

To access whether chemogenetic activation of L5 ET neurons induced NREM sleep locally or globally, EEG channels were independently classified using the neural network classifier described above. We quantified cross-channel agreement by computing classification accuracy for each channel relative to the prefrontal EEG channel (AP +2.30mm from Bregma), which was used as ground-truth labels for comparison across the remaining channels (AP +1.00 mm to -2.90 mm from Bregma).

### Analysis of Calcium Imaging Recordings

Miniscope videos were spatially decimated by a factor of 2 and temporally downsampled to a frame rate of 4 fps. Video frames were then Gaussian filtered and motion corrected using piecewise rigid alignment (NoRMCorre).^181^ Motion correction results were verified manually, and any motion related artifacts were typically less than 2 µm. Regions of interest (ROIs) corresponding to cell soma were manually selected. For each ROI, average pixel intensities were computed to obtain a single raw-fluorescence time-series. Raw calcium traces were detrended (3^rd^ order polynomial) to remove slow signal drifts and then normalized to their mean, resulting in ΔF/F values for cross cell comparison. Calcium transients were detected using the *findpeaks* function in Matlab. Peaks were required to exceed 100% ΔF/F, have a minimum prominence of 80% ΔF/F, and be separated by at least 1 second.

Two-photon calcium imaging videos were analyzed post hoc using ImageJ software version 2.1.0. All time-lapse images from each individual field of view were motion-corrected and referenced to a single template frame using cross-correlation image alignment (TurboReg plugin for ImageJ version 2.1.0). ROIs corresponding to visually identifiable somas were manually selected from the field of view. For each ROI, pixel intensities were averaged to generate a fluorescence trace. Background fluorescence was estimated as the average pixel value per frame from a region without GCaMP expression (e.g., blood vessel) and subtracted from the ROI trace. Baseline fluorescence (F₀) was calculated as the mean of inactive periods of the trace (∼2 s). Normalized ΔF/F_0_ traces were then used for calcium transient detection, defined as periods where the trace exceeded 3 standard deviations above the mean signal. Calcium traces shown throughout figures across individual groups are from single mouse to represent median activity level of respective group.

To determine whether NAP activity preceded NREM sleep onset, spontaneous Wake-NREM transitions were identified that were preceded by at least 1 min of continuous wakefulness and followed by 1 min of continuous NREM sleep. Calcium transients were aligned to the Wake-NREM transition and binned into consecutive 10second non-overlapping bins. Transient rates were estimated for each bin to quantify changed in NAP activity before and after NREM onset.

#### State Preference Estimation

Neuronal activity was estimated as the number of calcium transients (see above) per minute of recording. To quantify activity preference for a specific state *i* (e.g. NREM) we used a state preference index (*S*): 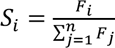 where *F*_*i*_ is the calcium transient frequency (transients/minute) observed in state of interest *(i)*, and the denominator is the sum of frequencies in all states n states in which this neuron was recorded. Thus, a value of 0 indicates a cell with no transients in state *i*, whereas a value of 1 indicates a cell with all transients exclusively in state *i*. *S*_*i*_ was computed for each neuron. The preferred state for each neuron was assigned as the state in which most transients were detected. Neurons in which no transients were observed in any state were excluded from the analysis.

To assess whether NREM and isoflurane state preference within the same neuron were coupled, a two-dimensional histogram was constructed (bin edges 0, 0.25, 0.5, 0.75, and 1 for both *S*_*iso*_ and *S*_*NREM*_). A null distribution was generated by randomly sampling NREM and isoflurane state preference indices from the same neuronal population. This preserves the marginal distribution of each state preference but destroys all inter-relationships between them. This procedure was repeated 100,000 times. For each histogram bin, the observed occupancy was compared with the corresponding permutation-derived null distribution, and empirical *p-values* were calculated as the fraction of permutations with occupancy greater than or equal to the observed value. Significance was determined using a Bonferroni-corrected threshold of alpha (0.05/16 bins).

#### c-Fos Colocalization to Layer 5 Confirmation Studies

RBP4 x Ai6 mice were exposed to 0.9% isoflurane or oxygen for 2 hours prior to brain harvest. Upon transcardial perfusion, brain harvest and overnight post fix in 4% paraformaldehyde, brains were placed in PBS with 0.02% sodium azide until ready for sectioning. A compresstome (Precisionary Instruments VF-510-0Z) was used to prepare 40µm thick sections for c-Fos and NeuN staining (1:4 series cut, 1 well stained). Sections were blocked in 2.5% normal goat serum in PBST (0.3% Triton X-100 in PBS) for 2 hours at room temperature, followed by overnight incubation at 4C in rabbit anti-c-Fos primary antibody (Cell Signaling Technologies 2250, 1:5000) and mouse anti-NeuN (abcam ab104224, 1:1000) in 2.5% normal goat serum in PBST. Sections were then washed in PBS, followed by a 2-hour incubation with Alexa Fluor 594 goat anti-rabbit (ThermoFisher 1:200) for c-Fos visualization and Alexa Fluor 647 goat anti-mouse (ThermoFisher 1:200) for NeuN visualization. Following a final wash in PBS, sections were mounted and coverslipped with mounting medium containing a DAPI counterstain (Southern Biotech 1011-20). Sections were imaged with a 20x objective lens using a confocal microscope (SP5II, Lecia Microsystems).

NeuN staining was used for automated neuronal soma detection as follows. Grayscale NeuN images underwent neural network classification using Ilastik^182^ using the NucleiSegmentationBoundaryModel from BioImage Model Zoo.^183^ Resultant probability maps were binarized in MATLAB at a threshold probability of 0.9. Region properties of the binarized cell detection map were computed and selected for further analysis using the following thresholds parameters: area threshold (in pixels) 30-1500, circularity .3-1.7, equivalent diameter 5-40. A pixel index list was created for each resulting segmented soma for each image separately.

To automatically determine whether a cell body was positive for c-Fos and/or Ai6, c-Fos and Ai6 channels were independently filtered by a 2-D gaussian smoothing kernel with a standard deviation of 2 pixels. Within identified neuronal cell bodies, presence or absence of stain was classified by a 2-state Gaussian mixture model based on median stain intensity within each NeuN-identified cell body. Posterior probabilities of greater than 0.9 were required to classify a cell as positive. Double labeled cells were identified as the union of c-Fos and ZsGreen positive. Manual regions of interest were drawn by a blinded experimenter to limit analysis to the prelimbic, infralimbic, and cingulate cortex. The number of positive neurons in each region was averaged across laterality.

Colgalt2-cre mice were bilaterally transfected with a cre-dependent tdTomato in the PFC (see coordinates and procedures above). After allowing time for viral expression, mice were exposed to 0.9% isoflurane or 100% oxygen for 2 hours prior to transcardial perfusion, brain harvest, and overnight post-fix in 4% paraformaldehyde. Coronal sections were collected as above and blocked in 2.5% normal goat serum in PBST (0.3% Triton X-100 in PBS) for 2 hours at room temperature, followed by overnight incubation at 4C in rabbit anti-c-Fos primary antibody (Cell Signaling Technologies 2250, 1:5000) in 2.5% normal goat serum in PBST. Sections were then washed in PBS, followed by a 2-hour incubation with Alexa Fluor 647 goat anti-rabbit (ThermoFisher 1:200) for c-Fos visualization. Following a final wash in PBS, sections were mounted, coverslipped, and imaged as described above.

Grayscale tdTomato or c-Fos images were independently analyzed using the “Pixel Classification + Object Classification” workflow in Ilastik for automated tdTomato-positive neuronal soma detection and c-Fos immunoreactivity, respectively. For pixel classification, all available pixel features were included for training. Foreground labels were manually assigned either tdTomato- or c-Fos-positive signal, while background labels were manually assigned to regions of nonspecific staining within the coronal section, ventricular spaces, or regions outside of brain tissue. For object classification, all available features were used to train the classifier. Resulting object identities were exported for further analysis in Matlab, where Colgalt2-tdTomato expressing neurons were classified as c-Fos positive when the Jaccard similarity coefficient between tdTomato-defined soma pixel-indices and any c-Fos object pixel index list exceeded 0.1. Manual anatomical regions were delineated by a blinded experimenter to restrict analysis to motor, cingulate, prelimbic, and infralimbic cortices. The fraction of double positive neurons within each region was quantified and compared across isoflurane and oxygen conditions.

To determine whether a subset of Colgalt2 PFC neurons is preferentially activated following isoflurane exposure, a separate cohort of Colgalt2-cre mice received unilateral injections of cre-dependent retrograde AAV into the claustrum (FLEX-tdTomato) and thalamus (FLEX-EGFP) to label the claustrum and thalamus projecting PFC Colgalt2 neurons. Isoflurane exposure, tissue processing, immunostaining, and imaging was performed in the same manner as described above. The same per channel “Pixel Classification + Object Classification” workflow was used for EGFP, tdTomato, and c-Fos expression in this dataset. Colocalization analysis between EGFP/c-Fos and tdTomato/c-Fos was performed as above. Colgalt2-expressing neurons were classified as dual-projecting if any tdTomato-defined soma pixel-indices overlapped with EGFP-defined soma pixel-indices.

### Verification of Chemogenetic Modulation with c-Fos Activity

Two hours after application of chemogenetic actuator (CNO or uPSEM), mice were euthanized and brains were perfused and harvested (as above). For c-Fos immunohistochemistry, sections were blocked in 2.5% normal goat serum in PBST (0.3% Triton X-100 in PBS) and incubated overnight at 4 °C with rabbit anti-c-Fos (Cell Signaling Technologies 2250, 1:5000). After PBS washes, sections were incubated with Alexa Fluor 647 goat anti-rabbit secondary antibody (ThermoFisher, 1:200), counterstained with DAPI, and coverslipped. Images were acquired using a Leica SP5II confocal microscope with a 20× objective.

Infralimbic subregions (∼1mm^2^) of grayscale c-Fos and viral expression images were processed separately using Ilastik, where Pixel Classification + Object Classification models were trained.^182^ The resulting object identity maps were exported and analyzed in Matlab. Co-localization was quantified by assessing the overlap of c-Fos identified ROIs with somas identified with viral expression.

### Thermal Imaging

Body temperature monitoring was performed in freely moving mice using a thermal imaging camera (FLIR Lepton 3.5, 160×120 pixels) positioned above home cages. Thermal images were acquired via the Lepton PureThermal SDK (v1.0.2) in MATLAB at 5 Hz. Cages contained standard bedding, nesting material, food pellets, and water. Mice were maintained in constant darkness to preserve free-running diurnal cycles. After 24 hr habituation, mice received IP CNO with ≥48 hr of subsequent acquisition.

Frames were down sampled to 0.1 Hz. The mouse was identified as the hottest 1% of pixels per frame, which was verified manually to capture the head and body while excluding cage surfaces. The median temperature of these pixels was computed and smoothed using a 5-min moving median to remove transient spikes caused by flat-field correction. Temperatures were normalized to a reference baseline defined as the mean over a 1 h period ∼25 h post-injection. Two time courses were analyzed: post-injection (from CNO administration) and a control period 24 h later at the same ZT to account for diurnal variation. Temperature changes between control and post-CNO periods were compared across hM3Dq and mCherry groups (50 minute windowed centered 90 min post-injection).

### Whole Brain Mapping of Neural Activity, Projections, and Monosynaptic Inputs

#### Viral Expression Timeline

To ensure sufficient expression of the anterograde viral label (tdTomato, see above) animals were euthanized two weeks following isoTRAP protocol in TRAP2-cre mice or 2 weeks following anterograde viral injection in Colgalt2-cre mice. For monosynaptic input tracing, a separate cohort of mice TRAP2-cre mice were first injected with a cre-dependent helper virus into the prefrontal cortex one week prior to isoTRAP exposure. Three weeks following isoTRAP, a genetically modified, replication-deficient rabies virus was injected into the same site to label direct presynaptic inputs of isoflurane active neurons (see above). Colgalt2-cre mice utilized for monosynaptic input tracing followed a similar viral transfection timeline of cre-dependent helper virus injected followed by a 3 week period for expression followed by injection of replication deficient rabies virus. To ensure optimal expression of neurons infected with G-deleted Rabies-EGFP, the survival time was set to five days following rabies virus injection for all mice.^105^ Following viral expression timelines for both projection and input groups, mice were exposed to 0.6% isoflurane for 4 hours. This was done to independently label anesthetic active neurons via c-Fos immunoreactivity. Following this isoflurane exposure, animals were rapidly euthanized and transcardially perfused using ice cold PBS with heparin (10 U/mL) followed by 4% paraformaldehyde. Brains were removed and post-fixed in 4% paraformaldehyde at 4°C overnight. Whole brains were then washed in PBS and shipped to LifeCanvas Technologies (Cambridge, MA) in a PBS with 0.02% sodium azide for further processing, staining, imaging, and mapping.

The c-Fos staining dataset in this study includes 18 brains stained for c-Fos in this study and an additional 10 brains subjected to identical protocol from our prior study from Wasilczuk et al.^74^ (WTM 1-5 and WTF 1-5 datasets found at https://zenodo.org/record/8384835 were incorporated in this analysis). Therefore, a total of 28 brains were used for c-Fos analyses in this paper.

#### Tissue Preservation, Clearing, Immunolabeling, and Imaging

Paraformaldehyde-fixed c-Fos and isoTRAP samples were preserved with SHIELD reagents (LifeCanvas Technologies: LCT) using the manufacturer’s instructions.^65^ Samples were delipidated using LCT Clear+ delipidation reagents. Following delipidation, samples were blocked using Antibody Blocking Solution (LCT) with 5% normal donkey serum (Jackson ImmunoResearch Laboratories Inc.) Samples were then labeled using the c-Fos optimized protocol (LCT) for primary labeling. This was followed by secondary labeling using SmartBatch+ Secondary Buffer system (LCT) with eFLASH^184^ technology which integrates stochastic electrotransport,^66^ using a SmartBatch+ device (LCT). Colgalt2 samples were not blocked with Antibody Blocking Solutions with 5% normal donkey serum, but were labeled with anti-RFP using SmartBatch+ Secondary Buffer system (LCT) with eFLASH^184^ technology which integrates stochastic electrotransport,^66^ using a SmartBatch+ device (LCT). After immunolabeling, samples were incubated in 50% EasyIndex (RI = 1.52, LCT) overnight at 37°C followed by 1 d incubation in 100% EasyIndex for refractive index matching. After index matching the samples were imaged using a SmartSPIM axially-swept light sheet microscope using a 3.6x objective (0.2 NA) (LCT).

#### Atlas Registration and Cell Detection

Samples were registered to the Allen Brain Atlas (Allen Institute: https://portal.brain-map.org/) using an automated process (alignment performed by LCT) at 25 µm x 25 µm x 25 µm resolution (1 voxel: 625µm^3^). For each brain, a chosen channel—NeuN, YoPro (nuclear dye), or autofluorescence—was registered to an average atlas of the same channel (generated by LCT from previously registered samples). Registration was performed using successive rigid, affine, and b-spline warping algorithms (SimpleElastix: https://simpleelastix.github.io/).

Automated cell detection was performed using a custom convolutional neural network through the Tensorflow python package. The cell detection was performed by two networks in sequence. First, a fully-convolutional detection network^68^ based on a U-Net architecture^68^ was used to find possible positive locations. Second, a convolutional network using a ResNet architecture^69^ was used to classify each location as positive or negative. For the isoTRAP sample subset, candidate cell centers were evaluated using watershed-based segmentation to isolate individual structures. Segmented objects were classified as cells if their total volume (including soma and proximal processes) was less than 2000 µm³—a threshold empirically determined from true positive examples. For the Colgalt2 sample subset, watershed segmentation was not implemented, but a final filtering step was performed for the anti-RFP cell detection to reduce false-positive cell candidates. Using the previously calculated Atlas Registration, each cell location was projected onto the Allen Brain Atlas in order to count the number of cells for each atlas-defined region.

#### Brain Wide c-Fos Density, Anterograde Projection, and Monosynaptic Input Estimation

c-Fos–positive neurons were assigned voxel coordinates, and density estimates (Figure 1) represent the mean number of neurons per voxel. Cortical region densities were computed by summing voxels corresponding to previously defined “Summary Structures”^70,74^ and layer-specific prefrontal densities were calculated using relevant Allen atlas structures^70^; differences were assessed with repeated measures one-way ANOVAs. Brains expressing tdTomato were similarly registered, and projection intensity estimates (Figure 3) represent mean voxel intensity. To normalize for differences in virus expression and imaging, intensity in each voxel was normalized by the total stain intensity computed over the entire brain excluding the injection site (prefrontal cortex: infralimbic, prelimbic, anterior cingulate) as well as olfactory-associated areas, cerebellum, and orbital regions. Monosynaptic input densities (Figure 3) represent mean input cell counts per voxel, calculated as the fraction of total inputs across the brain excluding the same regions. For connectivity graphs, only structures in the 90th percentile or higher of projection or input fraction were included.

### Ex Vivo Tissue Preparation

To establish cell-type properties of NAPs, ex vivo recordings from frontal cortex were obtained using 30-46 week old male and female activity tagged mice (see above). Mice were deeply anesthetized by IP administration of ketamine (130 mg/kg) and xylazine (8.8 mg/kg) mix and were perfused through the heart with chilled (2-4°C) sodium-free aCSF consisting of (in mM): 210 Sucrose, 7 d-glucose, 25 NaHCO_3,_ 2.5 KCl, 1.25 NaH_2_PO_4_, 7 MgCl_2_, 0.5 CaCl_2,_1.3 Na-ascorbate, 3 Na-pyruvate bubbled with carbogen (95% O_2_/5% CO_2_). Off-coronal slices (15 degrees tilted rostrally) 300 microns thick were generated using a Leica vibratome (VT1200, Leica, Germany) in the same sodium-free aCSF and were transferred to warmed (35°C) holding solution (in mM): 125 NaCl, 2.5 KCl, 1.25 NaH_2_PO_4_, 26 NaHCO_3_, 2 CaCl_2_, 2 MgCl2, 17 dextrose, and 1.3 sodium pyruvate bubbled with carbogen (95% O_2_/5% CO_2_). After 30 minutes of recovery, the chamber holding slices was allowed to cool to room temperature. Slices were maintained under these conditions until transferred to the recording chamber for patch-seq experiments.

#### Patch-Seq Recordings

Prior to data collection all surfaces and equipment were cleaned with RNAse Zap (Sigma-Aldrich) and nuclease-free water. Brain slices were placed in a submerged, heated (30-34°C) chamber that was continuously perfused with fresh, carbogenated aCSF consisting of (in mM): 125 NaCl, 3.0 KCl, 1.25 NaH_2_PO_4_, 26 NaHCO_3_, 2 CaCl_2_, 1 MgCl_2_, 17 dextrose, and 1.3 sodium pyruvate bubbled with carbogen (95% O_2_/5% CO_2_). Ionotropic synaptic receptor blockers were included 25 µM d-APV (Hello Bio), 20 µM DNQX (Sigma) and 100 µM picrotoxin (Tocris) in all recording solutions. Fluorescently labeled and adjacent neurons from activity-tagged mice (see above) were visualized with upright microscopes (Olympus BX51WI/Zeiss Examiner) equipped with infrared differential interference contrast (IR-DIC) or Dodt optics using 40x water immersion objectives. The somatic depth (150 to 700 µm from pia) of targeted neurons depended upon the cytoarchitecture of the cortical layers, which are more compact in ventral prefrontal cortex. Reflected fluorescent light was separated using two dichroic mirrors and through green (ET525/70 m-2P; Chroma) and red (ET650/75 m-2P; Chroma) emission filters. Whole-cell patch recordings were performed with pulled borosilicate glass pipettes (P1000, Sutter, Novato, CA) with tip openings about 1 μm and resistances of 3-7 MΩ, backfilled filled using 5-20 µl of internal recording solution consisting of (in mM): 110.0 K-gluconate, 10.0 HEPES, 0.2 EGTA, 4 KCl, 0.3 Na_2_-GTP, 10 phosphocreatine disodium salt hydrate, 1 Mg-ATP, 20 mg/mL glycogen, 0.5U/mL RNase inhibitor (Takara, 2313A), 0.5% biocytin and 0.02 Alexa 594 or 488 – pH adjusted to 7.3 with KOH. Whole cell somatic recordings were acquired using either an Axoclamp 2B or a Multiclamp 700A amplifier and custom acquisition software written in Igor Pro (MIES or using custom protocols written by Rick Gray and modified by N. Dembrow)^108^. Electrical signals were digitized at 20 or 50 kHz by an ITC-18 (HEKA) and were filtered at 10 kHz. The pipette capacitance was compensated, and the bridge was balanced (5-30 MΩ) during the entirety of the current clamp recording. Reported voltages were not corrected for the measured liquid junction potential (8 mV) and recordings with >30 MΩ series resistances or <0 mV spike overshoots were discarded. At the end of the recording, negative pressure was applied to collect the nucleus and was extruded into 10.4 µL collection buffer consisting of SMART-seq Takara lysis buffer, RNase Inhibitor (0.17 U/µl), and ERCC RNA Spike-In Mix 1 (1x10^-8^).

### Direct application of anesthetics on ex vivo slices

The direct effects of anesthetics on ET versus IT neurons were tested using 7-20 week old male and female mice from the Thy1h-eyfp, Etv1-egfp and Colgalt2-cre mouse lines prepared as described in ex vivo tissue preparation above. Patch-recordings were performed as described above (Patch-seq recordings). All experiments were performed with ionotropic synaptic blockers plus the GABA_B_ receptor antagonist 10 µM CGP 55845 (Sigma) throughout the recordings. Propofol (Sigma) was diluted to 25 mM stocks in DMSO and further diluted to working concentration the day of recording. Isoflurane (UW vet services) was administered to the slices using a custom-made sealed preincubation bottle that was continuously bubbled with carbogen (95% O2/5% CO2) delivered through an isoflurane well-fill vaporizer (Vapor 19.1, AES, Inc.). To ensure steady-state levels of isoflurane in the preincubation bottle was charged (at 1% isoflurane) for at least 45 minutes prior to administration. To calibrate appropriate concentrations of isoflurane in the recording chamber (Sup. Figure 13), samples were collected at 15 minutes intervals for HPLC analysis (see Measurement of Isoflurane below) immediately prior to switching the superfusion line from regular carbogenated aCSF to a fully pre-charged continuously bubbled isoflurane-carbogen solution for 45 minutes, followed by return to aCSF washout for an additional 30 minutes.

For all anesthetic bath application experiments, five minutes after initiating the patch recording, baseline measurements of subthreshold properties were performed at the resting membrane potential (see Patch clamp electrophysiological analyses below). Excitability was assessed using two methods. First, using 1-s long depolarizing current injections at a fixed current amplitude sufficient to drive between 5 and 9 action potentials (APs). The fold change in AP count to this fixed current injection amplitude was calculated as the ratio in the number APs elicited in the presence of anesthetic divided by the number during baseline conditions. We also measured excitability across a range of depolarizing current injection amplitudes (50-700 pA, 50 pA increment). The firing rate was calculated as the number of APs elicited from a 1 second current injection. To quantify the changes in the firing rate vs current (F-I) curves caused by anesthetic application, we subtracted baseline curves from 30 min after bath application of the drug.

### Electrophysiological Analyses

Three sets of stimuli were utilized to probe the electrophysiological properties of neurons. First, subthreshold chirp stimuli were sinusoidal current injections that increased logarithmically in frequency from 0.2 to 40 Hz over 20s. The amplitude of the chirp was adjusted for each neuron to produce a peak-to-peak deflection of 5-10 mV. Impedance amplitude profiles (ZAP) were calculated as the ratio of the Fourier transform of the voltage response to the Fourier transform of the chirp. The frequency at which the maximum impedance occurred was defined as the resonant frequency (fR). Resonance strength was defined as the ratio of the maximum impedance to the impedance at .5 Hz. The 3 dB cutoff was defined as the frequency at which the ZAP attenuated to the square root of 0.5 of the maximum impedance. The second set of stimuli comprised 11 subthreshold 1 s square wave current injections from -150 pA to 50 pA in 20 pA steps. Maximum and steady-state input resistance were calculated from the linear portion of the current voltage relationship in response to this protocol. Voltage sag was defined as the ratio of maximum to steady state input resistance. Rebound slope was defined as the slope of the rebound amplitude as a function of steady-state membrane potential. The third stimulus set consisted of a series of 1 s square wave depolarizing current injections that increased in amplitude by +50 pA/step to a maximum of 0.95 nA or until depolarization induced spike failure occurred. Threshold current was defined as the minimum current injection within this series to drive an action potential. Single action potential properties were measured from the voltage response of the first spike from at this current. This included action potential voltage threshold, defined as the voltage where the first derivative of the voltage response exceeded 20 V/s, maximum dV/dt, minimum dV/dt, and action potential width measured at half the amplitude between this threshold and the peak voltage. The first interspike interval (ISI) was measured at the minimal current step that drove at least two action potentials. Spike frequency accommodation (SFA), defined as the ratio of the second ISI to last ISI, was measured from the first spike train that produced at least 10 spikes.

### Processing of Patch-Seq Samples

Patch-seq samples were processed by standardized procedures developed by Allen Institute, as described in Lee et al. 2021^185^ and Dembrow et al. 2024.^108^ SMART-Seq v4 Ultra Low Input RNA Kit for Sequencing (Takara, 634894) and library construction using Nextera XT DNA Library Preparation Kit (Illumina FC-131-1096) were performed by Genewhiz/Azenta, Inc., following Takara kit instructions except at 0.2x reaction size. Samples were sequenced at a minimum of 17 million reads per sample using Illumina MiniSeq with 2x150 bp ends. Sequence reads were trimmed to remove possible adapter sequences and nucleotides with poor quality using Trimmomatic v.0.36. The trimmed reads were mapped to the Mus musculus GRCm38 reference genome available on ENSEMBL using the STAR aligner v.2.5.2b. The STAR aligner is a splice aligner that detects splice junctions and incorporates them to help align the entire read sequences. Unique gene hit counts were calculated by using the Subread package v.1.5.2. Transcripts per kilobase million (TPM) for each gene x sample were then mapped to the mouse 10x Whole mouse brain cell type taxonomy (CCN20230722) using both correlation and hierarchical clustering with MapMyCells (RRID:SCR_024672). Samples were included if they passed all of the following quality control metrics: >10 million mapped unique reads, >6000 genes detected, bootstrap probability at the class level of 1, and >0.45 supertype correlation score.

### Gene expression analysis

We utilized the publicly available snRNA-sequencing dataset derived from the mouse brain (WMB-10xV3)^55^ to compare gene expression of a non-exhaustive handpicked subset of anesthetic-targeted ion channels (SK, HCN, 2PK, Na leak). Reference dataset was acquired using Cv3 10x sequencing. The RNA-seq datasets were filtered to include samples from prelimbic, infralimbic, orbital (’PL-ILA-ORB’) and anterior cingulate area (‘ACA’) regions only, and infragranular layer subclasses (‘L5 ET CTX Glut’, ‘L6 IT CTX Glut’, ‘L5 IT CTX Glut’,’L4/5 IT CTX Glut’, and ’L6 CT CTX Glut’). For differential gene analysis in Figure 4 and Supplemental Figure 11, log2(counts per million (CPM) + 1), genes with at least 0.5 log2-fold difference in median gene expression between L5 ET (n=1656) versus L4-6 IT (n=11301) or L6 CT (n=7425) subclasses were compared using a two-sided Wilcoxon rank-sum test with a Benjamini-Hochberg false discovery rate correction done within each pairwise comparison. Effect size (r) was calculated as the absolute value of the Wilcoxon z-score following rank-biserial type conversion, divided by the square root of the total combined sample size. Differences labeled as statistically significant had an adjusted p value < 0.05 and effect size (r) of at least 0.3.

### Measurement of Isoflurane via HPLC

The experimental samples were stored in 2 mL gas-tight vials with minimal head space at -80 °C from the time of collection to analysis. Samples were allowed to thaw at room temperature and then vortexed to ensure a homogenous solution. They were then measured using a Beckman Coulter System Gold 126 HPLC, an Agilent Eclipse Plus C18 column (part #959990-902) equipped with a 500 μL injection loop, and a Waters 2410 Refractive Index Detector (phase: negative, sensitivity: 4, scale: 20, internal temperature: 34 °C). The mobile phase was run isocratically at 1 mL/min using the 150: 340: 510 mixture of 2-propanol: acetonitrile: 20 mM aqueous phosphate buffer (pH=4.6, tribasic sodium phosphate adjusted with *o*-phosphoric acid). The solvent system was made by adding 150 mL 2-propanol, 340 mL acetonitrile, and 102 mL 100 mM phosphate buffer stock solution to a 1000 mL graduated cylinder and diluting with water to reach a total volume of 1000 mL. This was then vacuum filtered through a 0.2 μm membrane filter (Thermo Fisher Scientific DS0215-4020). The isoflurane peak (retention time between 10 and 11 minutes) was integrated electronically (N2000 Chromatostation 3.50) and isoflurane concentration estimates were estimated from a standard curve. The standard samples (50, 100, 250, 500, 750, 1000, and 2000 μM isoflurane diluted in 1:1 methanol:water) were made fresh and stored in 2 mL gas tight vials with minimal headspace prior to analysis. A volume of 1 mL was injected for each sample to purposely overfill the injection loop (500 μL volume). All standards were run in tetraplicate, and experimental samples were run in triplicate (See Sup. Figure 13).

### Statistical Analyses

Animals were randomly assigned to experimental groups. Within each experiment, experimental and control groups were age matched. Sex was evaluated as a biological variable in the initial c-Fos mapping experiments. As no significant sex differences were detected in the overall PFC or Layer 5 c-Fos density, data from male and female mice were pooled for all subsequent analyses. The exception being experiments utilizing the Colgalt2-cre mouse line, where male mice were exclusively used given that the Colaglt2 transgene is Y-linked. Data were processed and analyzed in Matlab (2024a, Mathworks, Natick MA) using the Signal Processing, Image Processing, Curve Fitting, Deep Learning, Parallel Processing, and Statistics and Machine Learning Toolboxes. EEG spectral estimation was performed using Chronux open source software.^186^ Statistical comparisons were performed using Prism 10.6.0 (GraphPad Software, San Diego, CA, USA). Data were tested for normality, and appropriate parametric or nonparametric statistical comparison tests were used and reported within the figure legends. Reported values are (mean ± 95% CI) unless noted otherwise.

For all ex-vivo electrophysiology group comparisons, statistical testing was done using Igor Pro. Normality was tested using a Kolmogorov-Smirnov (KS) test. If the D statistic was larger than the critical value, normality was not assumed, and a Wilcoxon Rank test was used for comparison. D statistics smaller than the critical value were assessed as normally distributed, then tested for equal variances in the two samples, and then compared with a two-sample t-test. For all groups, a Cohen’s d statistic was also performed to measure the effect size of the difference between groups. P < 0.05 was considered statistically significant for all comparisons with appropriate corrections for multiple comparisons.

## Author Contributions

AZW, JC, MBK, ND, and AP conceived and designed experiments. AZW performed whole brain mapping, calcium imaging, chemogenetic, IHC imaging, behavior, and EEG experiments. DT performed miniscope surgery and recordings. JC performed 2P calcium imaging experiments and data processing. XZ assisted with whole brain mapping, chemogenetic, behavior, and EEG experiments and histology. EBB, HK assisted with miniscope imaging and analysis. JPM, MHK, WS, and ND performed patch-seq experiments, ex vivo anesthetic recordings and analysis. AK performed temperature recordings. ERW and BJK performed isoflurane concentration analysis using HPLC. AZW and AP developed coding scripts, performed general data analysis, created figures, and wrote the paper with input from all authors.

## Acknowledgments

We thank M. Fina for animal management and genotyping, M. Hudson for tissue preparation and histological processing, and D. Hunt of the Research Instrumentation Shop for technical support. This work is supported by T32GM112596-09 to AZW, AJK, and BJK, R35GM151160-01 to JC, BBRF Young Investigator Award to JC, American Society of Regional Anesthesia Chronic Pain Medicine Research grant to JC, T32NS105607 to EBB, R01 GM144377 to MBK/AP, R35GM156838 to ERW, Royalty Research Fund Grant GR032157 to ND, R01 GM151556 to AP/MBK, and the Department of Anesthesiology and Critical Care at the University of Pennsylvania. We would like to thank members of the Center for the Neuroscience of Unconsciousness and Reanimation Research Alliance for their helpful discussions about the data.

## Supplemental Figures

**Supplemental Figure 1:**
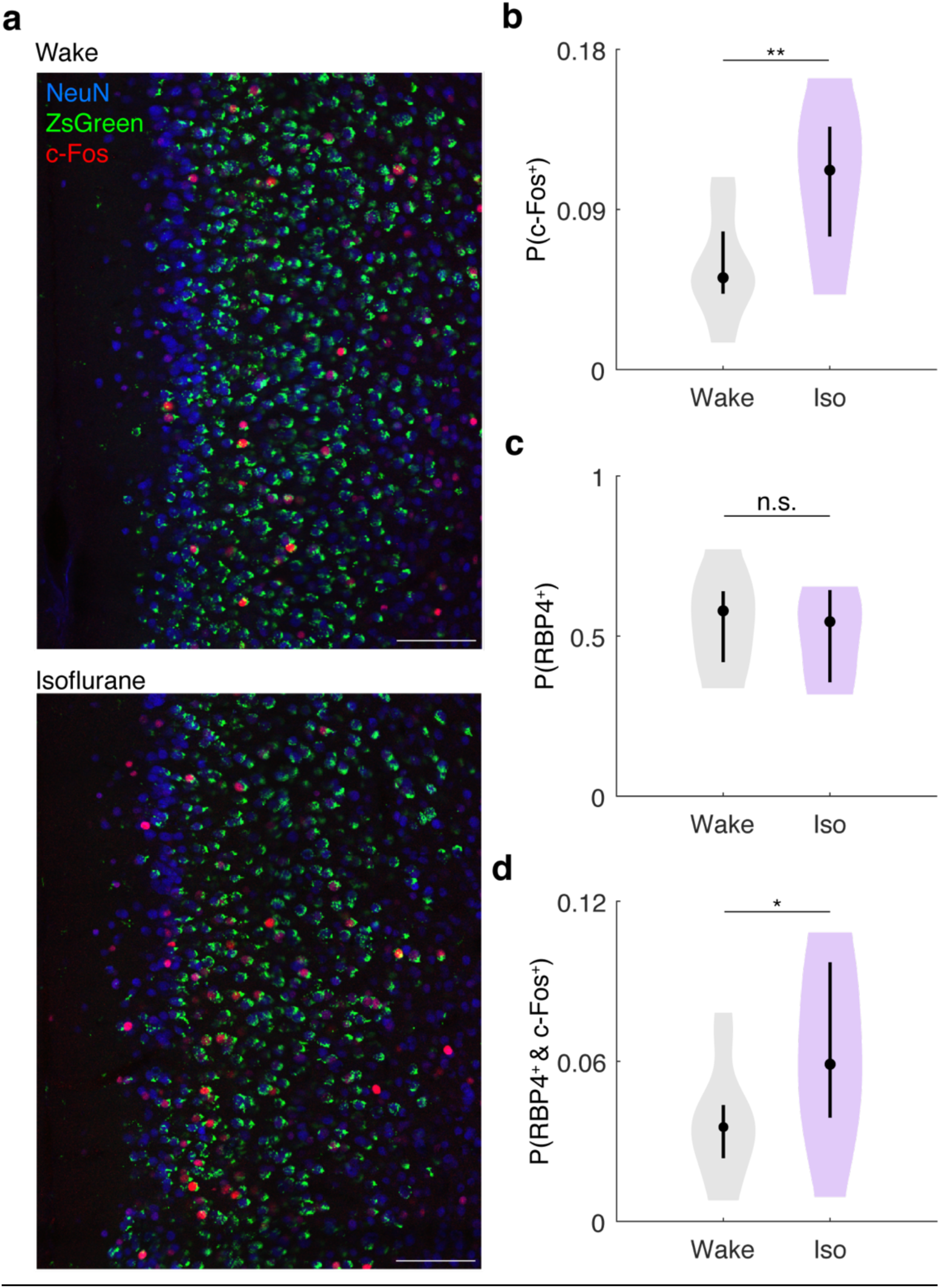
PFC Layer 5 neurons are preferentially active under isoflurane compared to wakefulness. RBP4 x Ai6 mice (n=4 per group) were exposed to either 0.9% isoflurane (iso) or 100% oxygen (wake) for 2-hours before rapid sacrifice, brain harvest and immunostaining for c-Fos (activity marker) and NeuN (neuronal marker). **a)** A representative coronal section through the PFC in the oxygen (top) and isoflurane (bottom) conditions. Calibration bar 200 μm. **b)** Fraction of PFC neurons (NeuN positive cells) expressing c-Fos during wakefulness is significantly less than under isoflurane (p=0.0021, unpaired t-test). **c)** Fraction of PFC neurons expressing ZsGreen (L5) is similar during wakefulness and under isoflurane (p=0.5977, unpaired t-test). **d)** Fraction of L5 c-Fos^+^ PFC neurons is higher under isoflurane than during wakefulness (p=0.0212, unpaired t-test). Data shown as violin plots (black circle shows the median, vertical bar shows interquartile range). Statistical significance denoted by *p<0.05, **p<0.01.

**Supplemental Figure 2:**
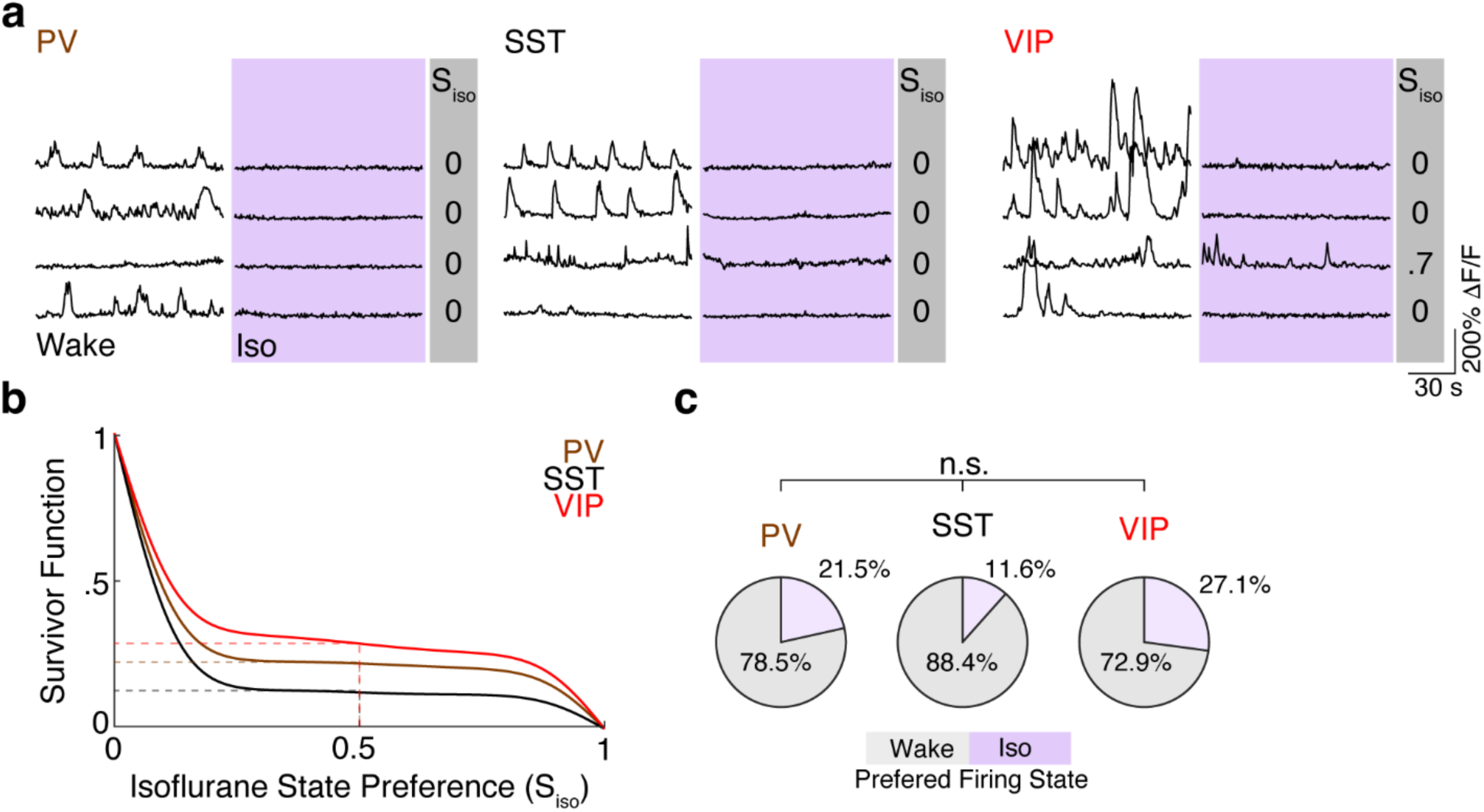
Wake-Isoflurane Activity of ACA Interneurons. **a)** Representative calcium activity traces from molecularly defined interneuron subtypes shown during wakefulness and under 0.6 % isoflurane (purple): Parvalbumin (PV), somatostatin (SST), and vasoactive intestinal polypeptide (VIP), showing state-dependent activity across wake and isoflurane conditions. Numbers shown at right margin indicate Isoflurane State Preference (S_iso_) for each neuron (PV n=4 mice, 79 neurons, SST n=3 mice, 69 neurons, VIP n=3 mice, 59 neurons). **b)** Survivor function (P) of S_iso_ across the three neuronal groups in **a.** P(0.5) reflects the fraction of neurons that exhibited higher activity under isoflurane (dashed lines). **c)** Binarized pie charts of preferred activity state. A chi-square test indicates no significant differences in the proportion of interneurons with isoflurane state preference between interneuron subtypes (χ^2^(2)=5.062, p=0.0796).

**Supplemental Figure 3:**
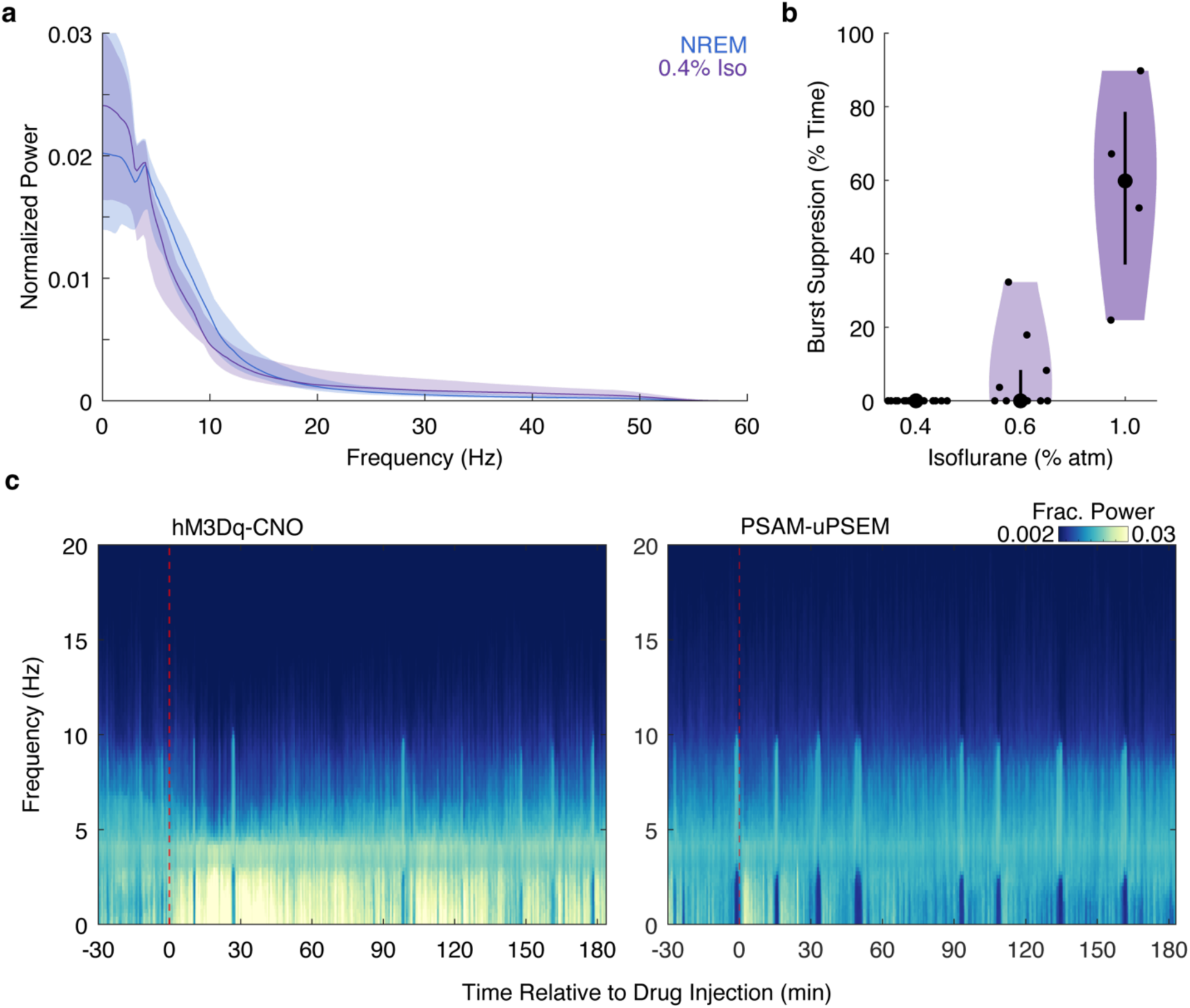
EEG signatures and chemogenetic modulation during isoflurane anesthesia. **a)** Mean EEG power spectra during NREM sleep (blue, n=48 mice: 19M 29 F) and 0.4% isoflurane anesthesia (purple, n=15 mice; 6M 9F). Mean and 95% CI across mice is shown. **b)** Fraction of time spent in burst suppression during 0.4%, 0.6%, and 1.0% isoflurane anesthesia. **c)** Representative EEG spectrograms before and after chemogenetic activation (hM3Dq + CNO, left) or inhibition (PSAM + uPSEM, right) of NAPs during steady-state 0.4% isoflurane anesthesia.

**Supplemental Figure 4:**
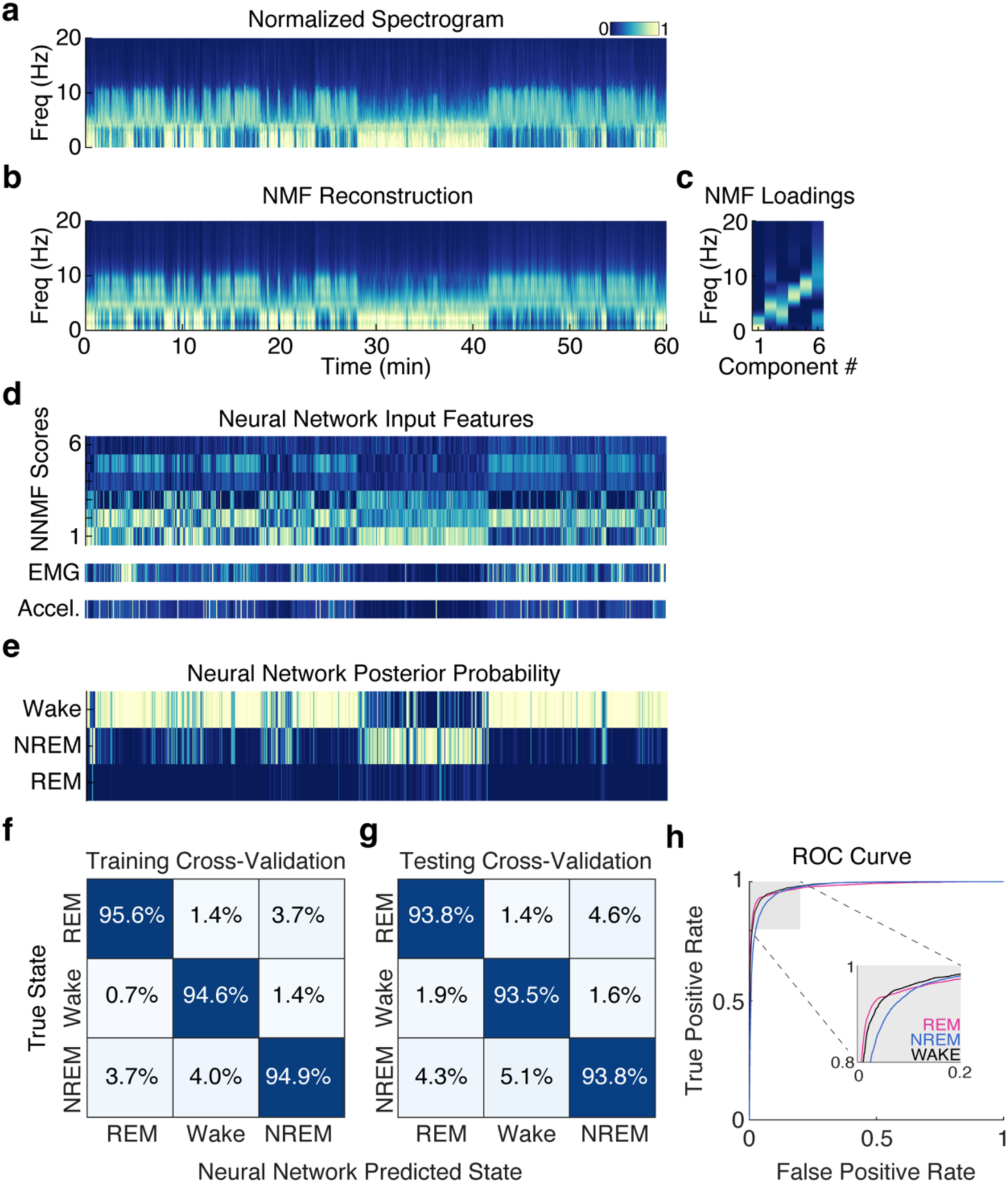
Neural network classifier performance for sleep-state prediction using NMF-derived features. **a)** Normalized EEG spectrogram over a 60 min period used as input for non-negative matrix factorization (NMF). **b)** Reconstructed spectrogram from NMF components. **c)** Frequency loading matrix showing spectral profile of each NMF component. **d)** Corresponding score matrix showing temporal activation of each component across time. The number of NMF components was optimized to explain spectral variance, with 6-components capturing 91.4% of total signal variance. These components, along with synchronized EMG and Accelerometer root-mean-square (RMS) signals were used as input features for neural network classification. **e)** Posterior probability for each predicted vigilance state. The state with the highest posterior probability at each time point was assigned the predicted label. **f-g)** Confusion matrices showing classifier concordance with expert labels (computed on 16 mice) during leave one out cross-validation during **f)** training and **g)** testing segments, demonstrating high classification accuracy across all three states. **h)** Receiver operating characteristic (ROC) curves for Wake (black), NREM (blue), and REM (pink) classification, with inset highlighting high-sensitivity region.

**Supplemental Figure 5:**
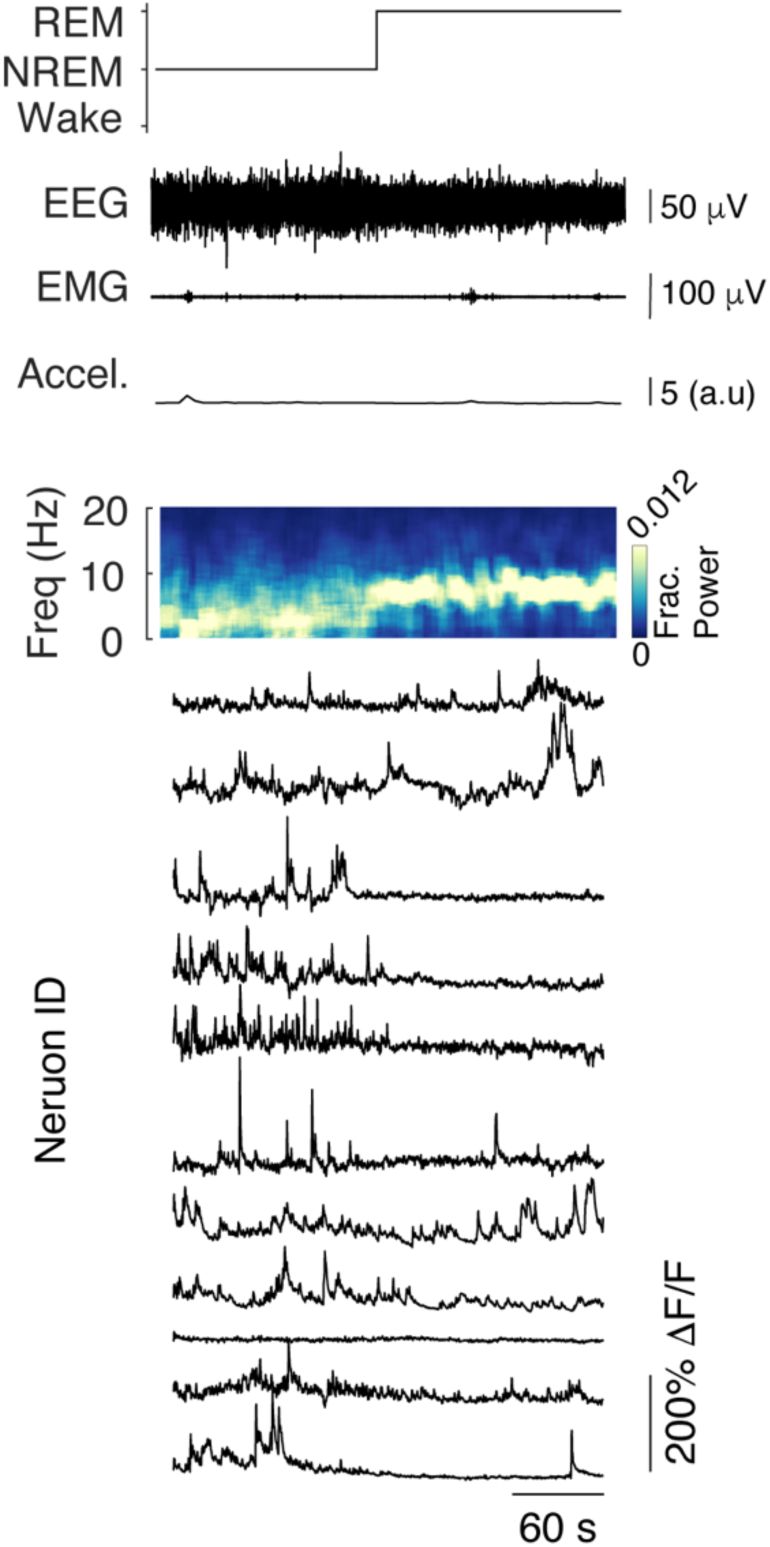
Simultaneous EEG, EMG, accelerometer, and calcium activity of NAPs during a NREM-REM transition. 5-minute recording shown. Top to Bottom: Hypnogram, EEG, muscle tone, and movement. Spectrogram is expressed as fraction of power per frequency. Calcium traces from prefrontal NAPs expressing GCaMP following isoTRAP.

**Supplemental Figure 6:**
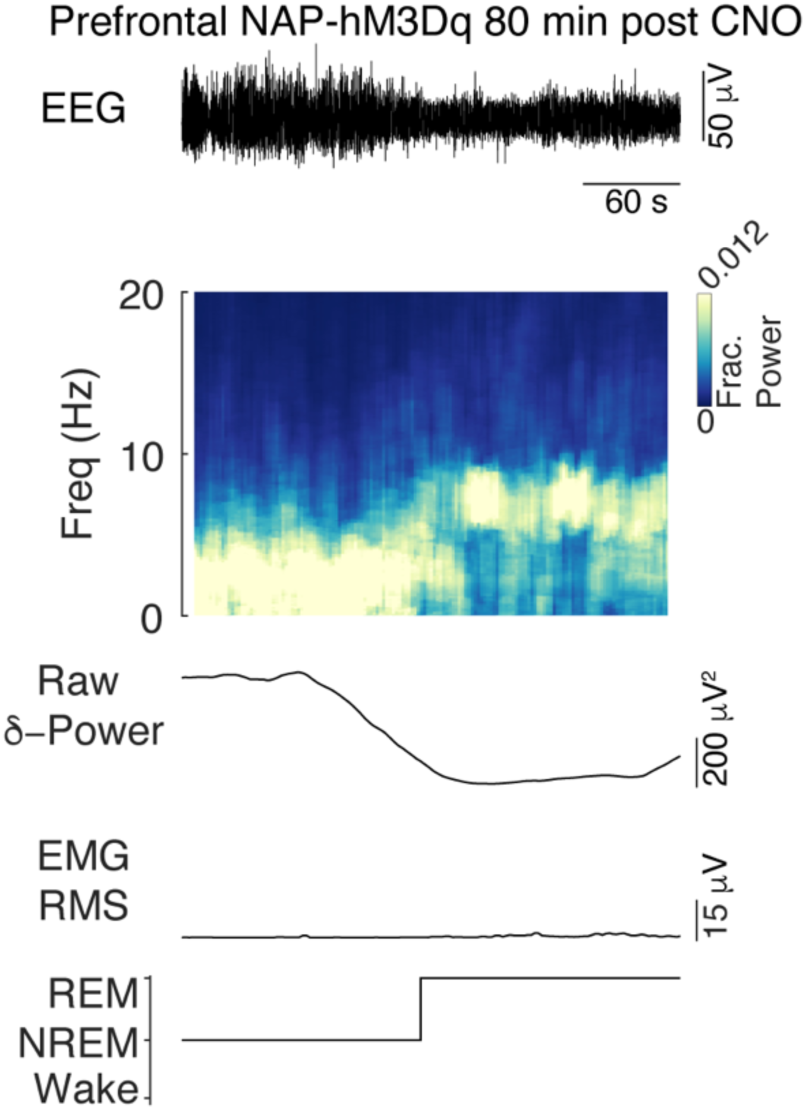
Example of NREM-REM transition following chemogenetic activation of NAPs using CNO. 5 min of data shown starting ∼80 min following CNO injection in isoTRAP hM3Dq mouse (same as in Figure 2h), illustrating NREM to REM transition. Top panel is EEG trace, second panel is normalized spectrogram, third panel δ-power, forth panel EMG RMS, and bottom panel is the hypnogram.

**Supplemental Figure 7:**
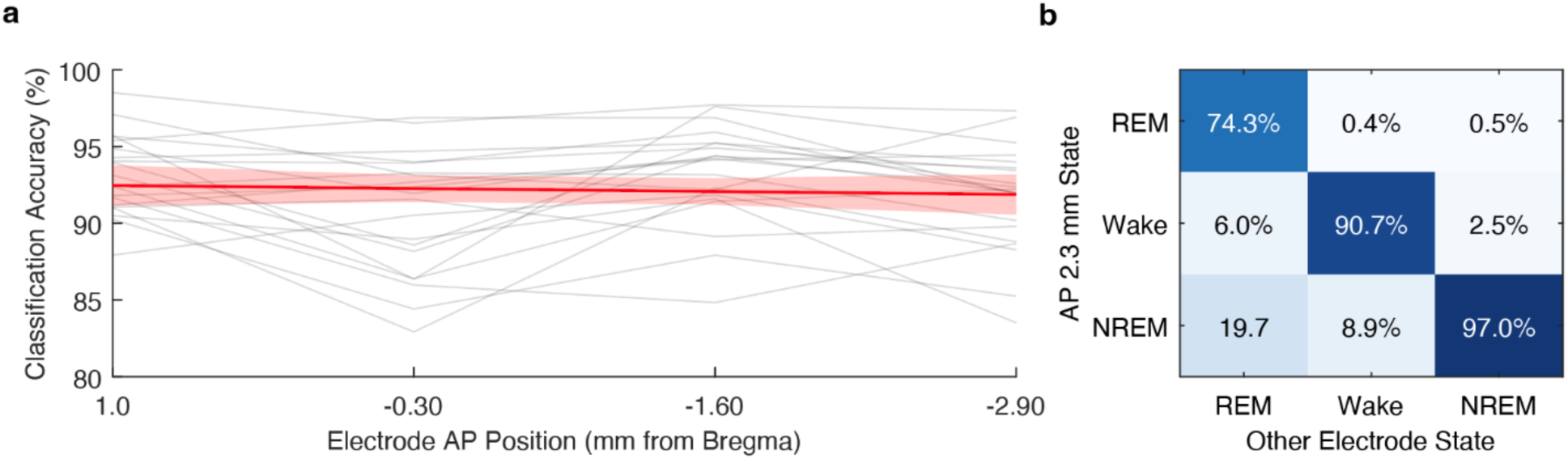
Sleep state classification across the anterior-posterior axis. **a**) Sleep-state classification was performed in 20 mice during the 4 hours following chemogenetic activation of NAP neurons. EEG signals from individual electrodes spanning the AP axis were independently classified using the neural network classifier shown in Sup. Figure 4. The same EMG and accelerometer RMS measures were used across channels for each mouse. Using the prefrontal EEG electrode (AP +2.30mm) as ground truth labels, classification accuracy was calculated for each electrode in each mouse (gray lines). Red line is best fit linear regression (Pearson’s r=-0.063, p=0.58). Classification accuracy did not significantly decrease as a function of distance from prefrontal electrode, suggesting vigilance state following NAP activation is globally coordinated across the cortex. **b)** Confusion matrix aggregated across mice during the same recording period.

**Supplemental Figure 8:**
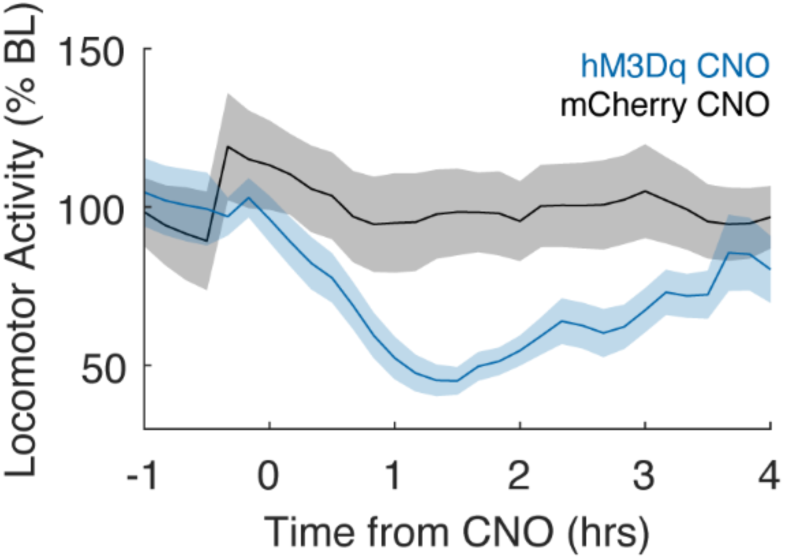
Chemogenetic activation of prefrontal NAPs reduces spontaneous locomotor activity. Time course of home cage locomotor activity following CNO injection (t=0) in mice expressing hM3Dq (blue, n=11) or mCherry (black, n=9) in prefrontal NAPs. Activity is normalized to baseline movement 1 hr prior to CNO injection Locomotor activity significantly decreased in hM3Dq, but not in mCherry group. (F(4,90)=51.43, p<0.0001, 2-way ANOVA, Šídák’s multiple comparisons test: mCherry BL vs. 0-1 p=0.9999, BL vs. 1-2 p=0.1638, BL vs. 2-3 p=0.9896, Bl vs. 3-4 p=0.4927; hM3Dq BL vs. 0-1, 1-2, 2-3, 3-4, all p<0.0001; mCherry vs hM3Dq BL p=0.2854, 0-1, 1-2, 2-3, 3-4, all p<0.0001). Solid line (shading) represents mean and 95% Jackknife CI.

**Supplemental Figure 9:**
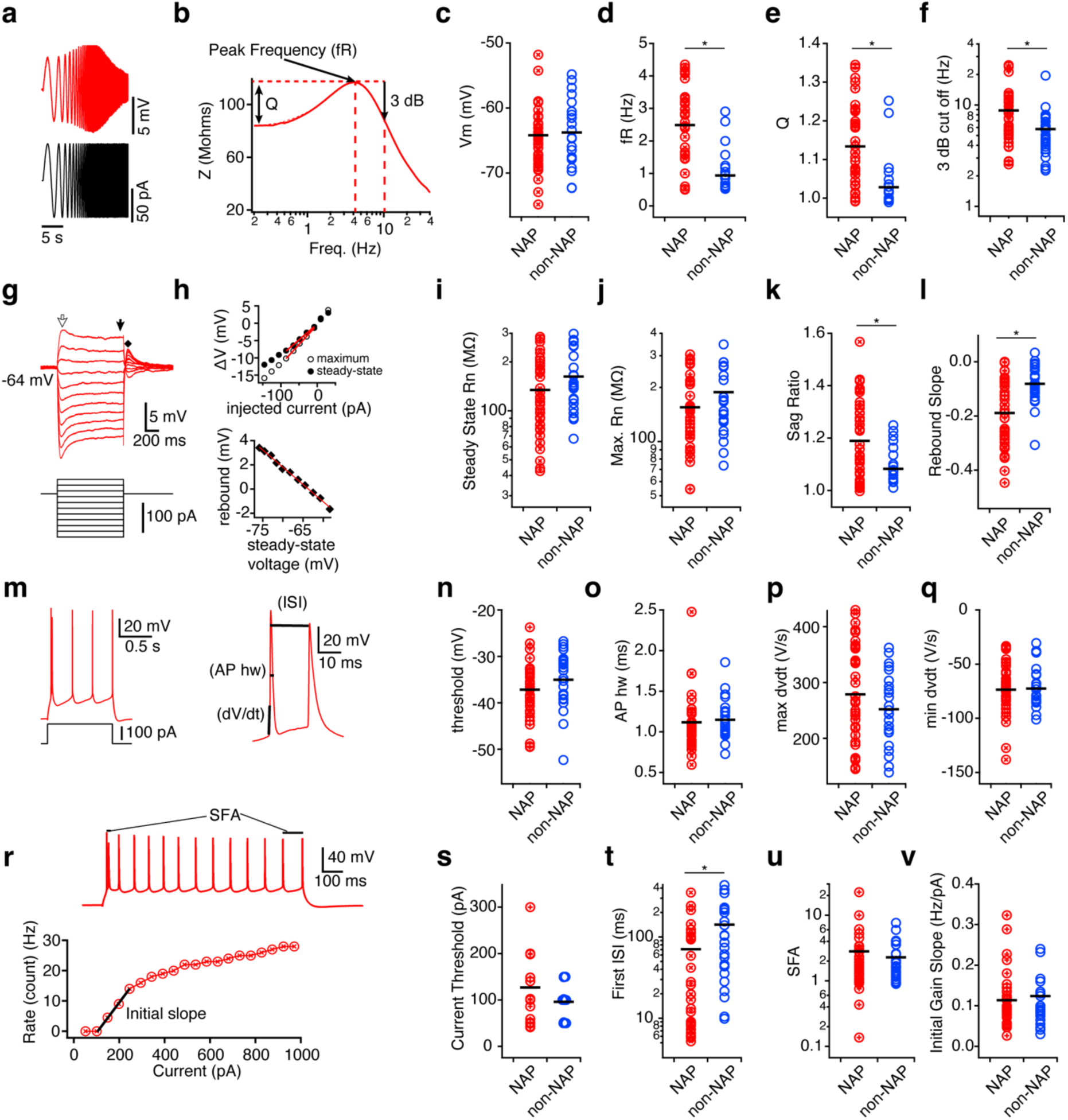
Electrophysiological properties of NAPs. Supporting to Figure 4. **a)** Voltage response (upper, red) of NAP to sinusoidal chirp current injection (black): ± 50 pA increasing from 0.2 to 40 Hz. **b)** The calculated impedance amplitude (Z) profile of **a**, with arrows denoting frequency with maximum impedance (resonant frequency, fR), relationship the resonant strength (Q), and the higher frequency at which the peak is diminished by 3 dB. **c-f)** Distribution of resting membrane potential (Vm), fR (U = 206.5, p <0.001, *d* = 1.35), Q (U = 194, p < 0.001, *d* = 1.13), and 3 dB cutoff (U=254, p < 0.001, *d* = 0.71) in NAPs (red circles, n = 37) compared to adjacent non-labeled pyramidal neurons (non-NAPs, blue circles, n = 26). Means are denoted by black bars. **g)** Representative traces of voltage response (red, upper) of NAP to subthreshold current step injections (black, below). **h)** Upper graph is the current-Voltage (I-V) plot of change in voltage in response at maximum (open circle, measurement location is open arrow in **g)** and steady-state (filled circle, measured at filled arrow in **g**). Red lines show linear slopes that are the input resistance (Rn). Lower graph shows for each steady-state voltage potential at the end of the current step, the magnitude of ‘rebound’ that occurs relative to rest after the current step is turned off. Red line shows slope from which rebound slope is calculated. **i-l)** Distribution of steady state and maximum input resistance, sag ratio (U = 256, p = 0.01, *d* = 0.77) and rebound slope (U = 196, p < 0.001, *d* = 1.06) of NAPs (n = 36) and non-NAPs (n = 22). **m)** Left, representative voltage response of NAP to the first depolarizing step current injection driving action potentials. Right, expanded initial voltage response of cell denoting single spike properties collected. **n-q)** Distribution of voltage threshold, action potential half-width (AP hw), maximum dV/dt, minimum dV/dt of NAPs (n = 34) and non-NAPs (n = 24). **r)** Upper trace, Representative voltage response of NAP to step current injection sufficient to drive at least 10 action potentials. Action potential firing rate was calculated as the number of action potentials driven at each current step for a 1s depolarization. Spike frequency adaptation (SFA) over the course of the step current injection is calculated as the ratio of the first to the last interspike interval (ISI). Lower graph, the number of spikes (plotted as frequency, F) versus injected current (I) of a NAP. **s-v)** Distributions for NAPs (n = 34) and non-NAPs (n = 24) of the minimum current required to drive at least one action potential, the first ISI after the first spike (U = 244, p <0.005, *d* = 1.06), SFA and initial slope of the FI. * Statistical significance denoted by * for samples with P < 0.01 from Wilcoxon rank test, U scores and Cohen’s d effect size reported for significant differences in throughout legend.

**Supplemental Figure 10:**
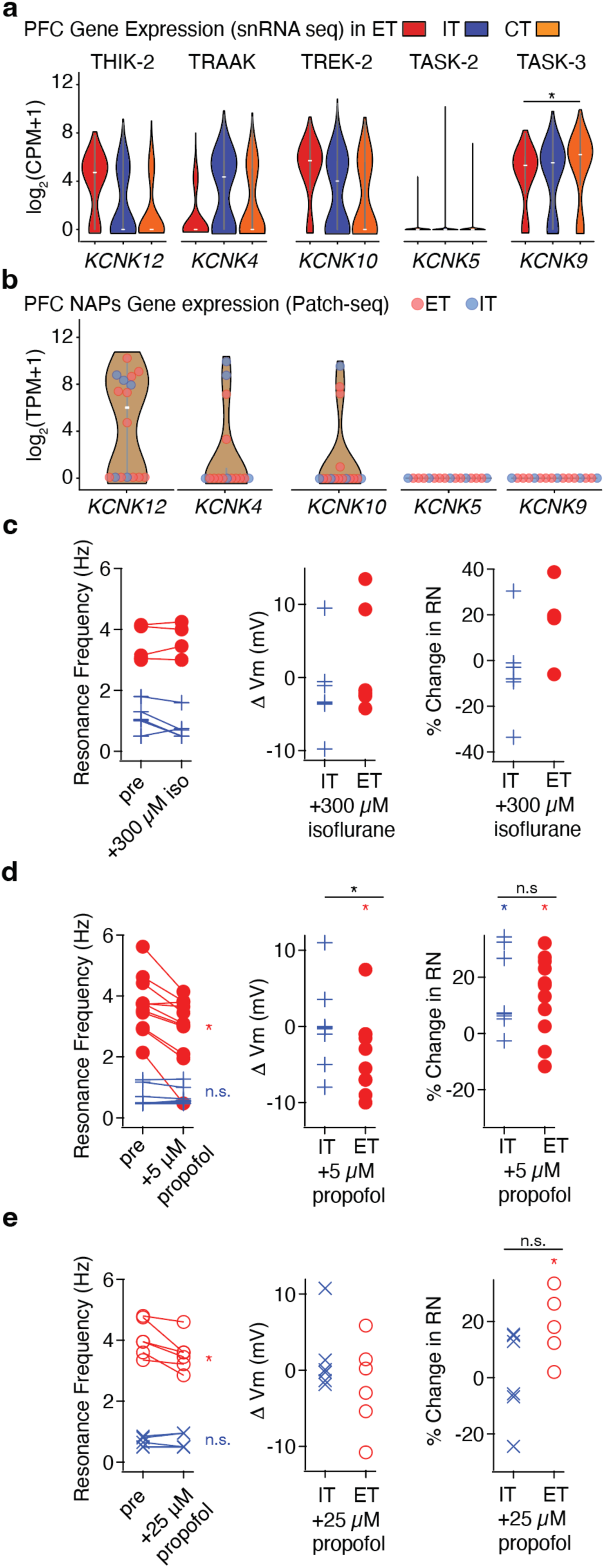
Supporting to Figure 4. Cell-type specific expression of additional 2-pore potassium channels genes in PFC pyramidal neurons and the effects of anesthetics on intrinsic physiological properties. **a)** Violin plots comparing gene expression across PFC L5 ET (n = 1656), L4-6 IT (n = 11301), and L6 CT (n = 7425) nuclei from the Allen Institute mouse brain dataset for several anesthetic-targeting 2-pore potassium channels. **b)** Violin plots of the same ion channel genes from patch-seq recordings of NAPs, showing broad concordance with the snRNA-seq data in (a). **c)** Changes caused by isoflurane (300 μM) in the frequency with the maximum impedance (resonant frequency, IT: n = 7, ET: n = 4) in response to subthreshold chirp stimuli (± 50 pA increasing from 0.2 to 40 Hz), resting membrane potential (IT: n = 7, ET: n = 6), steady state input resistance (IT: n = 7; ET: n = 5). **d)** Changes to the same features in (c) caused by 5 μM propofol (IT: n = 8; ET: n = 11) **e)** Changes to the same features in (c) caused by 25 μM propofol (IT: n = 6; ET: n = 6). Color (*) indicate significant difference from baseline (Wilcoxon Signed test, p <0.05), bars with (*) above indicate significant difference between change in ET and IT neurons caused by anesthetic (Wilcoxon Rank test, p <0.05). Significance: *p < 0.05.

**Supplemental Figure 11:**
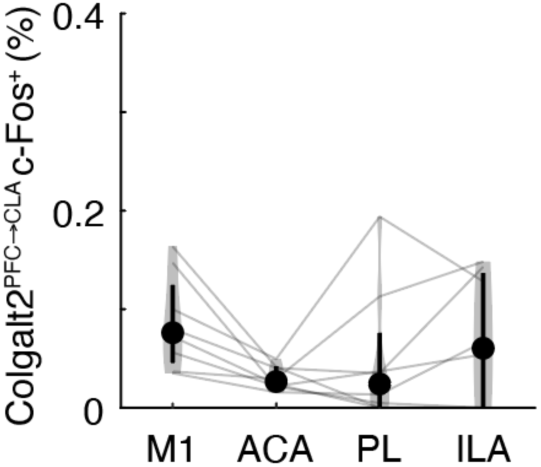
c-Fos expression in claustrum-projecting Colgalt2 neurons following isoflurane exposure. Following 2-hour 0.9% isoflurane exposure, two-way repeated measures ANOVA revealed significant effects of brain region (F(3,21)=11.27, p<0.0001) and projection target (F(1,7)=23.08, p=0.002), as well as a significant brain region x projection target interaction (F(3,21)=11.66, p=0.0001). Post hoc Tukey’s multiple comparisons demonstrate significantly greater c-Fos expression in thalamus-projecting (Figure 6e) than claustrum-projecting Colgalt2 neurons within anterior cingulate, prelimbic, and infralimbic cortices (all p<0.0001), but not within motor cortex (p=0.94). No regional differences were observed within the claustrum projecting population (all comparisons p>0.05).

**Supplementary Figure 12:**
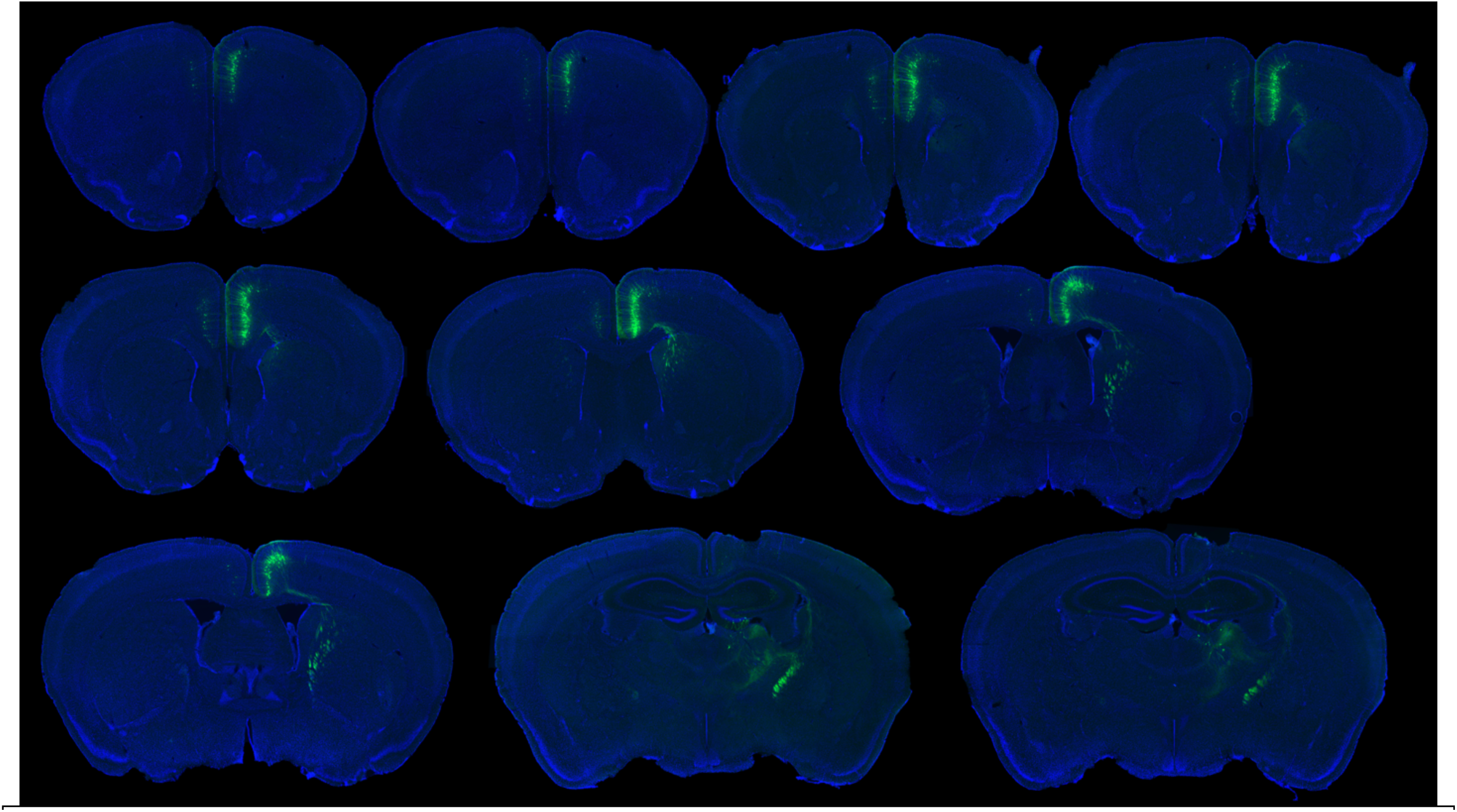
Retrograde labeling of thalamus-projecting Colgalt2 neurons. Representative serial coronal sections from a Colgalt2-Cre mouse following a unilateral right-sided injection of a Cre-dependent retrograde AAV-EGFP into the anterior thalamus. Sections arranged in anterior most to posterior most. Blue indicates DAPI counterstain and green indicates retrogradely labeled thalamus-projecting Colgalt2 neurons. Consistent with quantitative analyses in Figure 6d, labeled neurons are largely localized within the medial prefrontal cortex.

**Supplemental Figure 13.**
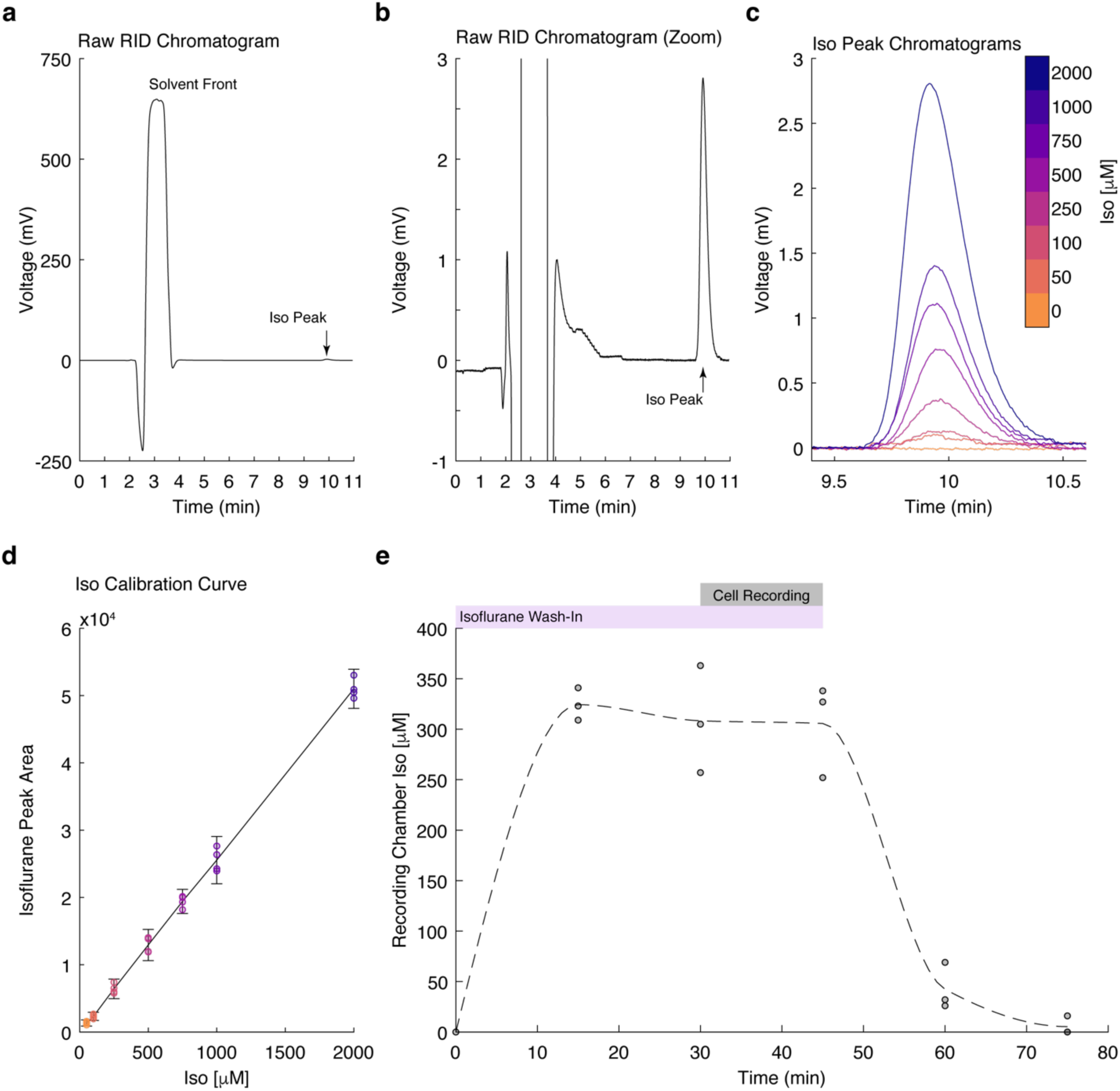
Quantification of Isoflurane by HPLC. **a**) Full HPLC RID chromatogram demonstrating an injection peak and a smaller signal from isoflurane at approximately 10 minutes elution time. b) Zoom in of the same chromatogram demonstrating clear elution of isoflurane at approximately 10 minutes. c) Chromatograms form one of four replicates of isoflurane standards used to generate the calibration curve. Samples of concentration 50-2000 μM) were used to generate the standard curve. d) Calibration curve of isoflurane calculated from tetraplicate standards. Linear regression and R2 value calculated in GraphPad Prism 11.02. (R2 =0.9966) e) Isoflurane concentration collected efflux limb of the perfusion chamber during wash-in and wash-out. Perfusion chamber samples were collected every 15 min and analyzed in triplicate (black points). Purple bar denotes the wash-in period, and gray bar denotes the period during which cell recordings were performed. The dashed line represents the estimated isoflurane concentration over time generated using MATLAB piecewise cubic Hermite interpolation (*pchip*). Experimental data used in Figure 4 and Sup. Figure 11 were collected during steady state period (gray bar) with average (± standard deviation) isoflurane concentrations of 308 μM (±53) and 306 μM (±47) respectively.

